# A transcriptional dissection of the petty spurge (*Euphorbia peplus* L.) reproductive structure

**DOI:** 10.1101/2024.05.06.592779

**Authors:** Arielle Rose Johnson, Ashley Bao, Margaret Hannah Frank

## Abstract

The unique reproductive structure of *Euphorbia* species, the cyathium, is an inflorescence that resembles a single complete flower. This structure has not yet been investigated using transcriptomics at a genomic scale. Enabled by the petty spurge (*Euphorbia peplus* L.) genome and guided by the ABCDE model of floral development, we dissected petty spurge cyathia and carried out a transcriptomic analysis of the different organs. We also constructed gene phylogenies and performed dN/dS analysis on select floral genes. Unlike in model species in the Asteraceae, which also have pseudanthia (false flowers), we did not find evidence for lineage-specific duplications of E class genes in *Euphorbia*. Rather, our transcriptomic and genomic analyses indicate that SEP1 paralogs have distinct expression patterns within the cyathium. *E. peplus* filiform structures (sterile organs within the cyathium) show upregulation of B and E class genes and transcriptomic signatures of heterochromatin formation, consistent with some form of floral identity and potentially consistent with their being reduced flowers in an inflorescence. The gene *CRABS CLAW* (*CRC*), which is involved in nectary development in both arabidopsis and petunia, is not upregulated in *E. peplus* nectaries, indicating an alternative nectary development pathway in *Euphorbia.* Finally, *Euphorbia* homologs for the inflorescence/floral meristem genes *UFO* and *LFY* and the B class genes *AP3* and *PI* have highly diverged sequences relative to other Euphorbiaceae species.

## INTRODUCTION

There are approximately 2000 species of *Euphorbia*, including horticulturally important plants such as poinsettia (*Euphorbia pulcherrima* L). The reproductive structure of *Euphorbia* species is called a cyathium. A cyathium resembles a complete flower with pistils, stamens, and a perianth; however, it is a pseudanthium (false flower) that is a condensed inflorescence consisting of a pistillate flower surrounded by staminate flowers and an involucral bract (Figure 1).

**Figure 1:**
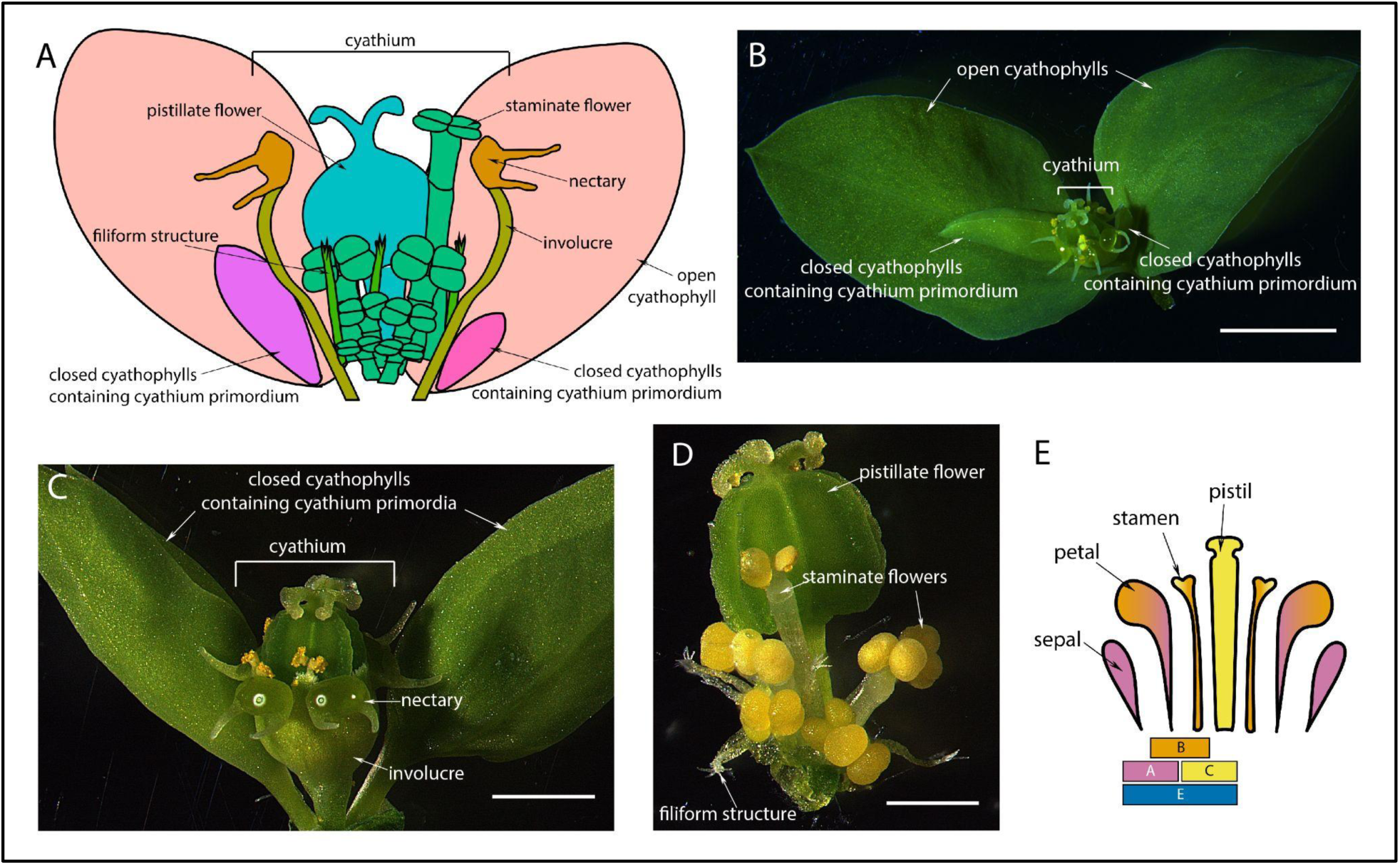
The *Euphorbia* cyathium is a unique reproductive structure. A. Diagram of a *Euphorbia peplus* cyathium; involucre is cut away to reveal filiform structures and staminate flowers inside. B. *E. peplus* cyathium and adjacent primordia and cyathophylls. Scale=2mm. C. *E. peplus* cyathium and adjacent primordia with adjacent cyathophylls removed. Scale=1mm. D. *E. peplus* cyathium with involucre and nectaries removed. Scale=500 microns. E. Diagram of the ABCDE model of floral development and the four whorls of a traditional flower; D class genes, which control ovule development, are not shown.

*Euphorbia peplus* L., commonly known as petty spurge, is an emerging model species for *Euphorbia* with a publicly available genome assembly (Czechowski et al., 2022; Johnson et al., 2023). In *E. peplus*, each immature cyathium is enclosed within two bracts, called cyathophylls. The cyathophyll contains the immature cyathium as well as multiple smaller cyathium primordia (enclosed in their own cyathophylls). The total number of primordia can vary by ecotype. Although *E. peplus* is primarily self-pollinated, the cyathium remains enclosed by cyathophylls until the stigma is receptive to pollen (indicated by maturation of stigma lobe morphology), which brings the cyathium into the open for pollination and later seed dispersal (Li et al., 2014).

The *E. peplus* cyathium itself, as in other *Euphorbia* species, is composed of a single pistillate flower in the center, surrounded by five mature staminate flowers. Each of the five mature staminate flowers is subtended by immature or nonfunctional staminate flowers in a zigzag pattern. Interspersed with the staminate flowers in the *E. peplus* cyathium are fringed filiform (threadlike) structures. These filiform structures have been referred to by various terms including “filamentous structures” (Prenner and Rudall, 2007) and “filiform bracteoles” (Bruyns, 2022). For the sake of clarity, we will refer to these structures as “filiform structures’’ in this paper, because “filamentous” implies a developmental homology between these structures and filaments, and because Prenner and Rudall 2007 also refer to cyathophylls as “bracteoles” (Prenner and Rudall, 2007). The pistillate flower, staminate flowers, and filiform structures are surrounded by a fused cuplike structure of involucral bracts, also known as the involucre. It is this cup-shaped structure that gives cyathia their name, as “cyathium” comes from the Greek word for cup, “kyathos”. In *E. peplus*, 4 or 5 crescent-shaped nectaries are present near the top of the involucre.

Although the cyathium has the superficial organization of a single flower (i.e., a pistillate structure surrounded by staminate structures that are surrounded by sterile organs), it is generally interpreted as an inflorescence based on several criteria (reviewed in Prenner and Rudall, 2007). There is a line of constriction on the staminate flower that is interpreted as the flower-stamen boundary, rather than individual stamens. Other genera in the Euphorbiaceae (e.g. *Anthostema*, Figure 2) form distinct staminate flowers that are clustered around a single pistillate flower. In addition, species within the Euphorbiaceae typically have unisexual flowers, and the bisexual composition of the cyathium would be an exception to this trend (Kaplan and Specht, 2022). Other genera within the Euphorbiaceae, such as *Dalechampia* and *Pera* have reproductive structures that have long been established to be pseudanthia (i.e., inflorescences that resemble single flowers; Gagliardi et al., 2018). Prenner and Rudall 2007 speculate that because the cyathium is an inflorescence, the sterile filiform structures that surround the staminate flowers could be interpreted either as prophylls or as highly reduced staminate flowers from a dichasial inflorescence (Prenner and Rudall, 2007). They interpret the involucre as a non-floral organ.

**Figure 2:**
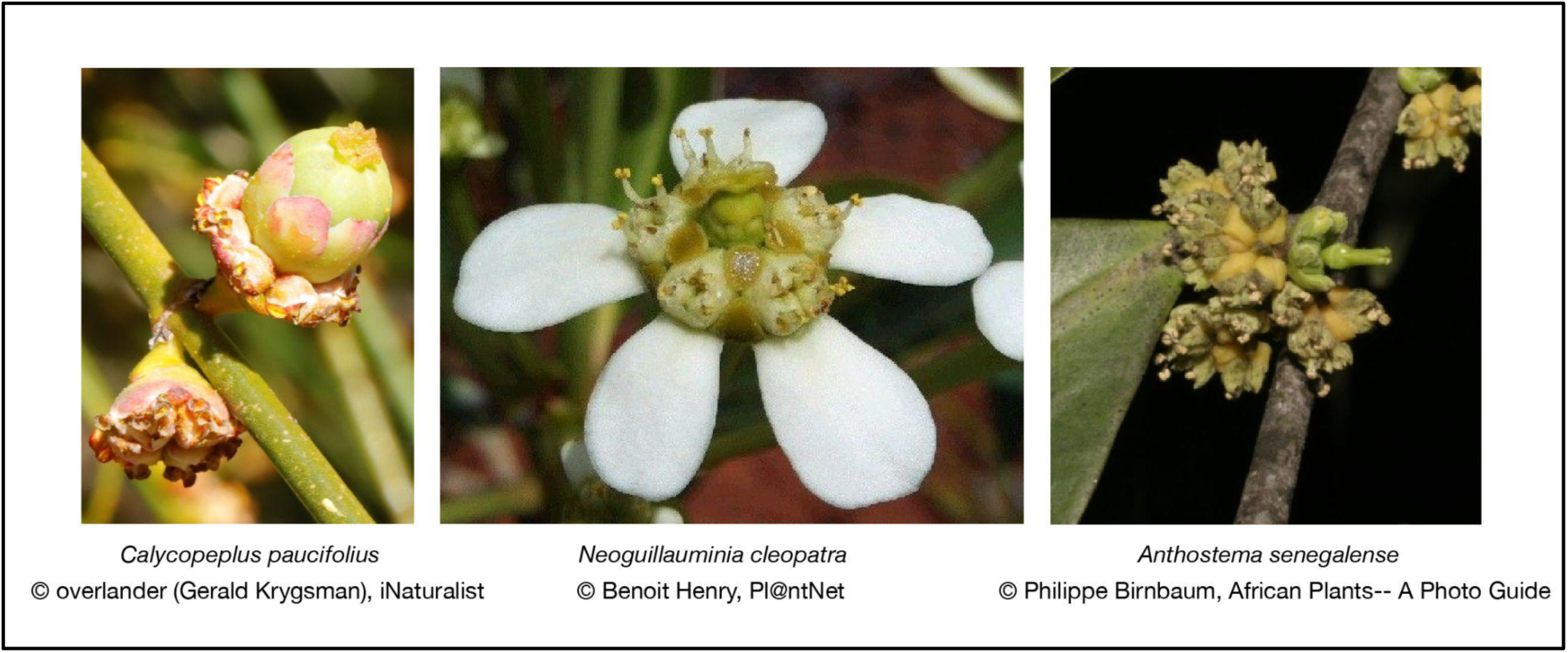
Close relatives of *Euphorbia*, *Calycopeplus* and *Neoguillauminia*, have perianths around their pistillate and staminate flowers and a condensed inflorescence similar to that of *Euphorbia*. *Anthostema*, which is less closely related, has perianths around its pistillate and staminate flowers and a less-condensed inflorescence.

As compared with its relatives such as *Anthostema* (Figure 2), the *Euphorbia* inflorescence is very compact structurally. Cyathia have a reduced axis, different placement of their flowers, and an earlier emergence of pistillate flower primordia compared with staminate flower primordia than inflorescences of related genera (Gagliardi et al., 2018). The Australian genus *Calycopeplus* and the New Caledonia tree *Neoguillauminia cleopatra* have been proposed as “missing links” towards the evolution of the *Euphorbia* cyathium from separate pistillate and staminate flowers; these close relatives to *Euphorbia* both have an inflorescence where a pistillate flower has its own small perianth that is then surrounded by staminate flowers with their own perianth whorls (Figure 2) (Western Australian Herbarium et al., n.d.; Prenner and Rudall, 2007; Prenner et al., 2008; Dressler et al., 2014; Gilbert, 1994).

Now that a high-quality genome is publicly available for two *Euphorbia* species, the cyathium can be investigated from a comparative molecular developmental perspective. This paper will use transcriptomics and evolutionary analysis to shed more light on the cyathium structure. Transcriptomics of floral development of another species in the Euphorbiaceae, *Jatropha curcas* L., showed that the ABCDE model of floral developmental regulation (Irish, 2017) is mostly conserved in *J. curcas* (Hui et al., 2017); therefore, we decided to investigate the ABCDE genes in *Euphorbia* as well.

Under the ABCDE model, different combinations of MADS-box transcription factors combine to specify the four floral whorls: the sepals, petals, stamens, and carpel (Figure 1E). E class genes are present throughout the flower and help specify all four whorls. C class transcription factors specify the carpel; B and C genes together specify stamens; B and A genes together specify the petal; and A genes alone specify the sepal. D class genes specify ovules.

Generally, B and C class genes are conserved between taxa, while A class genes are not highly conserved (Litt and Kramer, 2010). Complete flowers contain all of the floral whorls, while unisexual flowers have either functional stamens or functional carpels but not both. As defined by Mitchell and Diggle 2005, unisexual flowers can be classified based on the presence or absence of rudiments of the gynoecium or androecium (Type I = rudiments present; Type II = rudiments absent) (Mitchell and Diggle, 2005). The *Euphorbia* cyathium has Type II unisexual flowers, with no evidence of sterile carpels/stamens present. The complete disappearance of a reproductive whorl in Type II unisexual flowers is hypothesized to result from differential regulation of the MADS-box transcription factors (Feng et al., 2020). The flowers of the *Euphorbia* cyathium also lack a perianth (sepals and petals). Loss of the perianth whorls can result from edits to MADS-box transcription factors or to other transcription factors not related to whorl identity, as is seen in the arabidopsis *petal loss* (*ptl*) mutant (Brewer et al., 2004).

We also wanted to contextualize the *Euphorbia* cyathium as a pseudanthium (false flower). False flowers can form from multiple types of meristems. Claßen-Bockhoff and Bull-Hereñu 2013 use an ontogenetic framework to draw distinctions between two different types of flower-producing meristems: ‘flower unit meristems’ (FUMs) which are determinate and lack apical growth, and inflorescence meristems (IMs) which are initially indeterminate and show apical growth (Claßen-Bockhoff and Bull-Hereñu, 2013). IMs produce the basic branched inflorescence types, e.g. racemes and panicles, whereas FUMs are relatively similar to floral meristems and generate dense heads of flowers. Head meristems in Asteraceae are considered FUMs based on their mantle-core structure and lack of apical growth (Zhang and Elomaa, 2021). Morphological work on *Ricinus communis* (Euphorbiaceae), reinterpreted its “staminate flower” which forms many-branched “staminal trees” as a multi-flower unit with each flower reduced to a single anther, forming from an FUM (Claßen-Bockhoff and Frankenhäuser, 2020). The *Euphorbia* cyathium similarly has a meristem that could be interpreted as an FUM, as it initially lacks visible primordia (Prenner and Rudall, 2007). Because of their ontogenetic similarity, we compared the *Euphorbia* cyathium with the Asteraceae pseudanthium.

## MATERIALS AND METHODS

### Plant growth conditions

*E. peplus* plants were grown under long day conditions with 25 °C day/16 °C night. Seeds were germinated without any pretreatment in closed Phytatrays (P5929; Sigma-Aldrich, Saint Louis, MO, USA) in LM 1-1-1 soil mix, then week-old plants were uppotted to 4-inch pots containing LM 1-1-1 soil mix.

### Imaging

Images of cyathia were taken using a Leica M205 FCA stereo microscope with a DMC6200 camera.

### Fixation and dissection for RNA sequencing

Cyathia from twelve-week-old *E. peplus* plants were collected into ice-cold acetone then stored at -20°C until dissection. Cyathia were dissected under the dissecting scope in acetone on a cold block and dissected organs were put immediately into chilled acetone tubes. 5 biological replicates of 10 different organs were collected, with 10 cyathia dissected for each biological replicate.

### Stages used for RNA sequencing

We used primordia approximately 200 μm tall for our “small primordium” samples. We used cyathium primordia approximately 500 μm tall for our “large primordium” samples. We used staminate flower primordia approximately 200 μm tall for our “young staminate flowers” and pistillate flower primordia approximately 400 μm tall for our “young pistillate flowers”. For the remaining samples, we dissected them from nearly-mature cyathia still enclosed within unopened cyathophylls when cyathia were approximately 1.5 mm tall not including cyathophylls: nectary, involucre, mature staminate flower, mature pistillate flower, filiform structure. In our organ stages, ‘mature’ here refers to developmental maturity of form, not reproductive maturity, which roughly coincides with the opening of the cyathophylls. At reproductive maturity, cyathia are approximately 2mm tall.

### RNA extraction

RNA was extracted using the New England Biolabs Monarch Total RNA Miniprep Kit (NEB #T2010, Ipswich, MA, USA). Acetone was removed from the tubes under the dissecting scope, 2-3 beads (BioSpec Products #11079124, Bartlesville, OK, USA) were added, the tubes were placed into liquid nitrogen, and the tissue was ground in a bead beater (Qiagen TissueLyser II, Germantown, MD, USA). 400uL of 1x protection reagent from the kit was added to the ground tissue and beads, and the tissue was ground again in the protection reagent. The liquid was transferred to a new tube and the rest of the kit protocol was followed. RNA was eluted with 100uL nuclease-free water.

### RNA purification

The 100uL of RNA was purified by adding 10uL 3M sodium acetate, 0.75uL GlycoBlue (ThermoFisher #AM9515, Waltham, MA USA) and 275uL ice-cold 100% ethanol, vortexing to mix, precipitating at -80°C overnight, centrifuging at 13000 rpm at 4°C for 30 minutes, washing the pellet twice with 0.5mL ice-cold 75% ethanol, air-drying the pellet for 5 minutes, and resuspending the RNA in 20uL of nuclease-free water.

### 3’ sequencing

RNA samples were submitted to the Cornell Institute for Biotechnology Genomics Center for 3’ mRNA-seq library construction using a QuantSeq FWD kit (Lexogen, Vienna, Austria) followed by 1×75 bp sequencing on an Illumina NextSeq500 (Illumina, Hayward, CA, USA). Sequencing statistics were calculated with v0.15.0 seqkit stats with parameters -j 10 -T(Shen et al., 2016). (Supplementary File 1).

### Orthogroup inference

To assemble orthogroups, we ran OrthoFinder v2.5.4 (Emms and Kelly, 2019) on the longest isoform protein sequences from *E. peplus*, *E. lathyris*, arabidopsis, soybean, Pará rubber tree, cassava, poplar, and castor bean using the parameters -S diamond -t 40 -a 8(Buchfink et al., 2021). Orthogroups are included in Supplementary File 2.

### Identification of ABCDE floral identity genes

In order to identify floral identity genes of interest, we retrieved the sequences of genes of interest from Uniprot accessed through TAIR and ran blastp with our *E. peplus* proteins file as the query and the parameters -qcov_hsp_perc 80 e-value 1e-10(Camacho et al., 2009). Where BLAST results surfaced multiple candidates with low e-values and high percent identity, a reciprocal BLAST to arabidopsis was used to select the best candidate(s). We further confirmed the results through comparison to the clades identified by OrthoFinder. Gene IDs accessed through TAIR: AT1G69120(AP1)(Bowman et al., 1993); AT4G36920(AP2)(Jofuku et al., 1994); AT3G54340(AP3)(Jack et al., 1992); AT4G18960(AG)(Mizukami and Ma, 1997); AT2G45650(AGL6)(Ma et al., 1991); AT1G69180(CRC)(Alvarez and Smyth, 1999); AT1G65480(FT)(Kobayashi et al., 1999); AT5G60910(AGL8)(Mandel and Yanofsky, 1995); AT5G61850(LFY)(Schultz and Haughn, 1991); AT5G20240(PI)(Goto and Meyerowitz, 1994); AT5G15800(SEP1)(Pelaz et al., 2000); AT1G24260(SEP3)(Pelaz et al., 2000); AT3G58780(SHP1)(Liljegren et al., 2000); AT2G42830(SHP2)(Liljegren et al., 2000); AT2G45660(AGL20)(Lee et al., 2000); AT4G09960(STK)(Favaro et al., 2003); AT5G03840(TFL1)(Kobayashi et al., 1999); AT1G30950(UFO)(Levin and Meyerowitz, 1995).

### Transcriptomic analysis

Raw read files were filtered with fastp v0.23.2 using parameters --cut_right --cut_window_size 5 --cut_mean_quality 20 --length_required 50 (Chen et al., 2018). Transcript abundance was generated using *E. peplus* coding sequences with kallisto quant v0.42.4 with parameters -- single -b 100 -t 10 (Bray et al., 2016). The resulting data was processed in R using the DESeq2 pipeline; all genes with >20 estimated counts across all samples were considered to be expressed and were included in the analysis (Love et al., 2014). Variance-stabilized-transformed data were visualized in heatmaps using the package pheatmap with expression normalized by row and genes clustered by expression pattern (Kolde, n.d.). Of the floral genes of interest investigated (Supplementary File 3), the *E. peplus* homologs for AGL8/FUL (Ep_chr6_g18041), TFL1 (Ep_chr7_g23231) and two paralogs of FT (Ep_chr8_g25489, Ep_chr6_g20489) had expression too low (i.e. 20 or fewer counts across all samples) to be analyzed. Genes are said to be differentially expressed in this paper if padj<0.05. All DESeq filtered untransformed data is included in Supplementary File 4. Comparisons of differentially expressed genes between different tissues are included in Supplementary File 5.

### Biological Process and Molecular Function GO term enrichment analysis

GO terms were assigned to *E. peplus* genes as specified in Johnson et al 2023 (Johnson et al., 2023); GO terms assigned to each gene are included in Supplementary File 6. In order to find which GO terms were enriched in differentially expressed genes in different organs as compared with cyathophylls, we ran TopGO v.2.46.0 using the “classicCount” option, using Fisher’s test to determine significance (Alexa and Rahnenfuhrer, 2021). TopGO results are included in Supplementary File 7. We summarized and visualized the GO results using REVIGO and downloaded the R scripts to edit the color scale and axis limits (Supek et al., 2011). Terms with a dispensability higher than 0.9 were removed and clustered with other terms for ease of visualization. Only terms with a dispensability less than 0.2 were labeled with text in the visualization.

### Manual curation of OrthoFinder clades

Genes with highly truncated annotations were removed from the analysis (LFY: rubber tree KAF2311631.1, soybean Glyma.04G202002.1.p; PHYA: soybean Glyma.03G227333.1.p; PI: *E. lathyris* Casp05761.t1; SEP1: rubber tree KAF2325472.1). For PI, *E. lathyris* Casp33119.t1 and *E. peplus* Ep_chr8_g25689.t1 were included in the PI phylogenetic analysis despite being classified into their own orthogroup because they clearly were a result of a tandem duplication (Casp33120.1 and Ep_chr8_g25690.t1 were included in the orthogroup with Arabidopsis PI, AT5G20240.1) and because of high reciprocal BLAST scores with Arabidopsis PI.

### Gene alignment, gene tree construction and selection analysis

For each clade of interest (generated by OrthoFinder then manually curated as specified above; see Supplementary File 3 for gene names), a preliminary alignment was generated using MAFFT v7.522 with parameters --maxiterate 5000 --auto --adjustdirectionaccurately --thread 8 -- op 3. Then macse v2.07 alignSequences was used to generate a codon-aware alignment with default parameters. For a few alignments, macse generated exclamation points due to sequencer errors; these were converted to N. Then alignment trimming was performed in trimAl v1.4 with parameter -gappyout. Finally, sequences were exported for phylogeny construction using macse v2.07 exportAlignment -codonForFinalStop NNN -codonForInternalStop NNN - keep_gap_only_sites_ON. To generate phylogenetic trees, RAxML NG v1.2.0 was run with parameters --all --model GTR+G --threads 12 --bs-trees 1000(Kozlov et al., 2019). In order to generate dN/dS values for each branch of the tree, PAML v4.10.6 was run on the best tree from RAxML with model=1 and NSSites=0. In order to test for relaxed selection at sites within genes, PAML v4.10.6 was run on the best tree from RAxML with model=2 and NSSites=2. Alignments were visualized in ESPript 3.0 online (https://espript.ibcp.fr/ESPript/cgi-bin/ESPript.cgi)(Robert and Gouet, 2014). Sites flagged for positive selection based on the Bayes empirical Bayes method (BEB) are included in Supplementary File 8.

### Motif analysis

In order to retrieve a dataset that could be used to detect the canonical LFY binding motif, a bed file of LFY ChIPseq targets (ChIPseq MACS2 q-value 10-10, DANPOS Mnase-seq q-value 10-50) was taken from Supplementary Data 1 from Jin et al 2021 (Jin et al., 2021). This was converted to a fasta file of the target sequences using bedtools getfasta (Quinlan and Hall, 2010). Then MEME-ChIP from MEME v5.5.2 was used to detect the LFY canonical binding motif using default parameters except -maxw 25 (Machanick and Bailey, 2011). The dLUBS and mLUBS motifs were acquired from the authors of Rieu et al 2023 in pfm format and converted to meme format using jaspar2meme -pfm from MEME v4.11.2 (Bailey et al., 2015; Rieu et al., 2023b). Then FIMO from MEME v5.5.2 was run with default parameters on a fasta file of the 3kb upstream of the start of the LFY target genes AP1, AG, AP3, and PI in order to assess whether the motifs were present or not (Grant et al., 2011). MEME FIMO motif search results are included in Supplementary File 9.

### Genome assemblies used for analysis

The Johnson et al *Euphorbia peplus* genome assembly and annotation was accessed through Zenodo (Johnson et al., 2023): https://zenodo.org/records/8033914 The Wang et al. *Euphorbia lathyris* assembly and annotation was accessed through figshare (Wang et al., 2021): https://figshare.com/articles/dataset/High-quality_genome_assembly_of_the_biodiesel_plant_Euphorbia_lathyris/14909913/1 The Liu et al. *Hevea brasiliensis* (rubber tree) assembly and annotation was accessed through NCBI (Liu et al., 2020): https://www.ncbi.nlm.nih.gov/assembly/GCA_010458925.1/. The Xu et al. wild *Ricinus communis* assembly and annotation was accessed through NCBI (Xu et al., 2021): https://www.ncbi.nlm.nih.gov/datasets/genome/GCF_019578655.1/. The Bredeson et al. *Manihot esculenta* v8.1 assembly and annotation was accessed through Phytozome: https://phytozome-next.jgi.doe.gov/info/Mesculenta_v8_1 The arabidopsis Araport 11 genome release was accessed through TAIR (Cheng et al., 2017): https://www.arabidopsis.org/download/index-auto.jsp?dir=%2Fdownload_files%2FGenes%2FAraport11_genome_release%2FAraport11_blastsets. The *Glycine max* (soybean) Wm82.a4.v1 assembly and annotation was accessed through Phytozome (Valliyodan et al., 2019): https://phytozome-next.jgi.doe.gov/info/Gmax_Wm82_a4_v1. The *Populus trichocarpa* v4.1 assembly and annotation was accessed through Phytozome (Tuskan et al., 2006): https://phytozome-next.jgi.doe.gov/info/Ptrichocarpa_v4_1

## RESULTS

### Analysis of ABCDE gene expression in cyathium components

We dissected the different organs of the cyathium and sequenced their RNA (Figure 3 A-D). Of 25,473 annotated *E. peplus* genes, 17,164 were expressed in the cyathium. Staminate flowers had the most distinct expression pattern from other organs; a clear separation is visible between the mature staminate flowers, young staminate flowers, and other cyathium organs in the first principal component (PC) of the data (Figure 3E). This data is consistent with many other flowering plant species’ transcriptomic pattern of distinct staminate flowers compared with other organs, or distinct stamens compared with other floral organs in complete flowers (Klepikova et al., 2015; Xanthopoulou et al., 2021).

**Figure 3:**
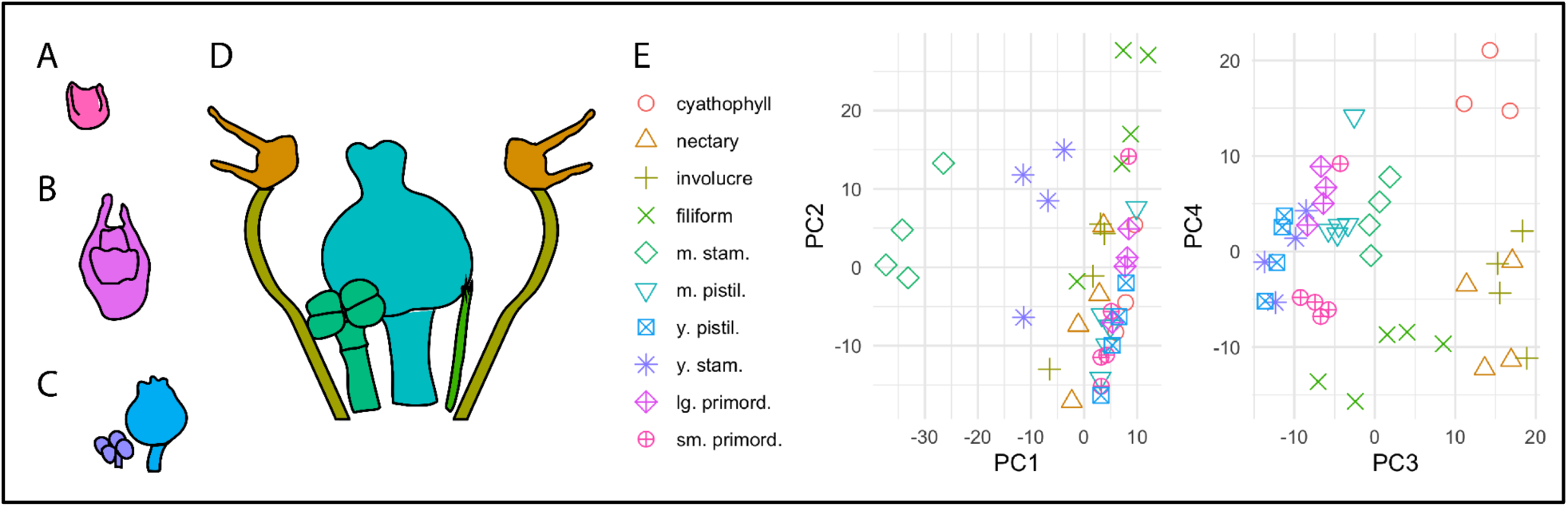
Staminate flowers have a distinct expression pattern. A-D. Illustration of cyathium parts used for RNAseq sampling. A. Small primordium. B. Large primordium. C. Young staminate flower and young pistillate flower. D. Mature organs: nectary, involucre, staminate flower, pistillate flower, filiform structure, involucre, nectary. (Duplicates of organs are omitted for simplicity. Cyathophyll is not shown.) E. PCA plot of variance-stabilized-transformed data for the 500 genes with the most variance. PC1 explains 22.4% of the variance, PC2 explains 18.9% of the variance, PC3 explains 16.7% of the variance, and PC4 explains 10.6% of the variance.

Using BLAST and OrthoFinder, we identified *E. peplus* homologs for all classes of the ABCDE genes and visualized their expression pattern across different organs (Figure 4).

**Figure 4:**
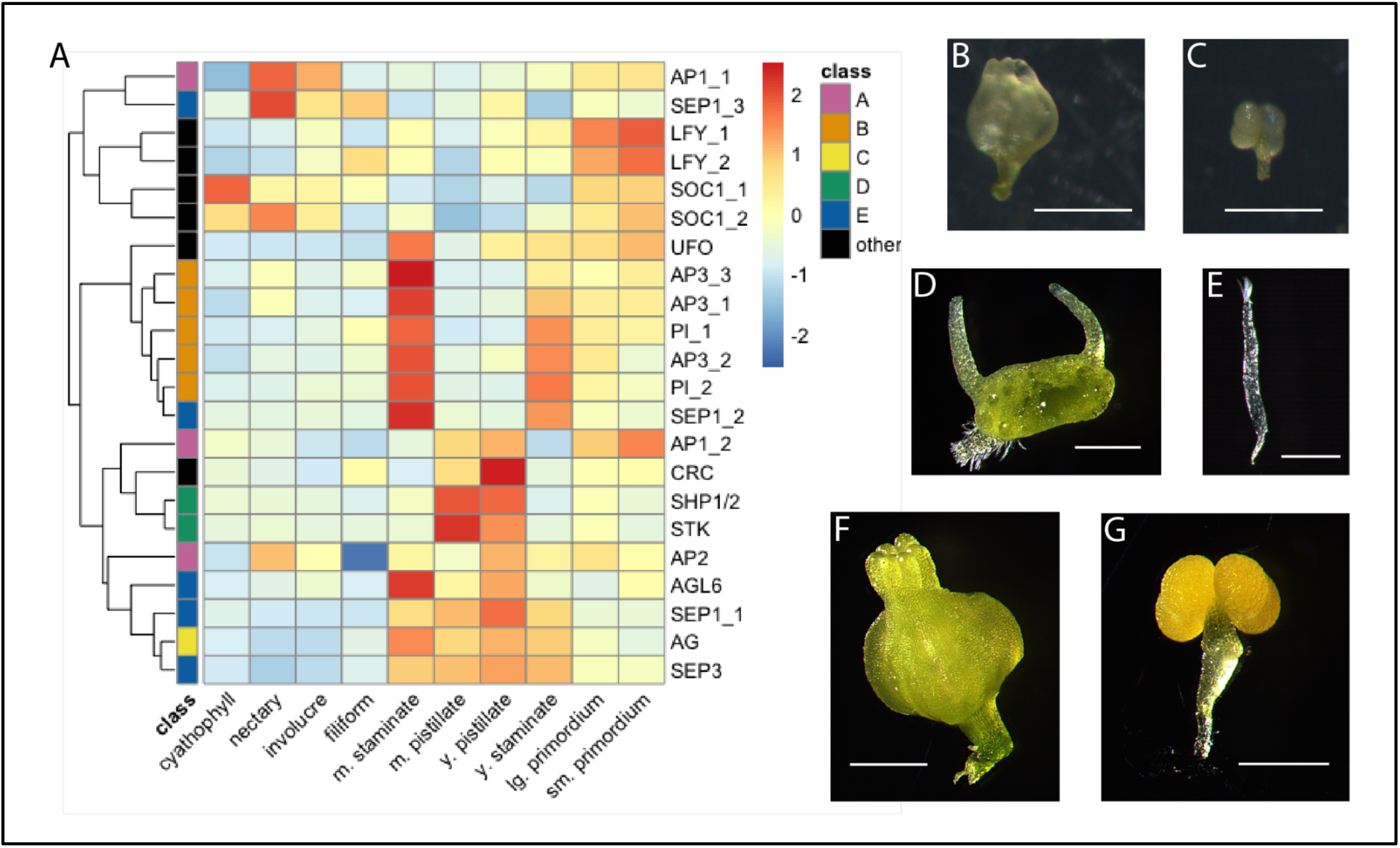
E class gene SEP3 is highly expressed in the pistillate and staminate flowers. A. Heatmap shows expression of ABCDE homologs and other floral identity genes in different organs of the *E. peplus* cyathium. B-N. Images of the organs sampled. B. Young pistillate flower. C. Young staminate flower. D. Nectary. E. Filiform structure. F. Mature pistillate flower. G. Mature staminate flower. Scale bars = 250 microns.

In the ABCDE model, the E class genes SEPALLATA1 (SEP1) and SEPALLATA3 (SEP3) are partially redundant in function and help specify all four floral whorls; a closely related E class gene, AGAMOUS-LIKE 6 (AGL6), also helps specify all of the floral organs and is also involved floral meristem regulation (Dreni and Zhang, 2016; Morel et al., 2019). There are three orthologs of SEP1 in *Euphorbia*, which is consistent with other Euphorbiaceae (Supplementary Figure 1). *E. peplus* SEP1_1 is significantly upregulated in both stages of pistillate flowers compared with cyathophylls, SEP1_2 is significantly upregulated in staminate flowers as compared with pistillate flowers, and SEP 1_3 is significantly upregulated in the filiform structures as compared with cyathophylls (Figure 4, Supplementary File 5). *Euphorbia*’s sole SEP3 ortholog is significantly upregulated in both stages of the pistillate and staminate flowers, both stages of the primordia, and the filiform structures as compared with cyathophylls. AGL6 is not significantly expressed in any tissue as compared with the cyathophylls due to low expression overall (Supplementary File 4, Supplementary File 5).

The B class genes APETALA3 (AP3) and PISTILLATA (PI) specify petal and stamen identity; this role is conserved across bisexual and unisexual flowers (Litt and Kramer, 2010; Zhang et al., 2022). All three *E. peplus* AP3 paralogs and both PI paralogs are significantly upregulated in staminate flowers compared with pistillate flowers at both stages studied, in mature staminate flowers as compared with cyathophylls, in young staminate flowers as compared with cyathophylls (except AP3_3), and in filiform structures as compared with cyathophylls (Figure 4, Supplementary File 5). Of the AP3 and PI homologs, only AP3_2 is significantly expressed in the involucres as compared with the cyathophylls.

The C class gene AGAMOUS (AG) is deeply conserved across evolution and helps specify both pistillate and staminate development (Dreni and Kater, 2014). The *E. peplus* ortholog of AG is significantly upregulated in both stages of both pistillate flowers and staminate flowers as compared with cyathophylls, and not significantly upregulated in any other parts of the cyathium (Figure 4, Supplementary File 5).

In arabidopsis, the A class genes APETALA1 (AP1) and APETALA2 (AP2) help specify sepals and help specify petals in conjunction with B class genes, in addition to other functions in floral meristem identity and determinacy (Han et al, 2014). Notably, these genes’ functions are not conserved outside the Brassicaceae and are unlikely to be the same in *Euphorbia* as in arabidopsis (Litt and Kramer, 2010; Zumajo-Cardona and Pabón-Mora, 2016). There are two homologs of arabidopsis AP1 in *E. peplus*. AP1_1, which is most similar in sequence to arabidopsis AP1 and CAULIFLOWER (Supplementary Figure 2), is upregulated in the nectary, involucre, filiform structures, both stages of primordia, both stages of staminate flowers, and young pistillate flowers as compared with cyathophylls (Figure 4, Supplementary File 5). *E. peplus* AP1_2 and AP2 are not significantly differentially expressed in any cyathium part as compared with cyathophylls, likely due to low gene expression (Supplementary File 4, Supplementary File 5).

We were also interested in the genes LEAFY (LFY) and UNUSUAL FLORAL ORGANS (UFO) which are known to directly activate many of the ABCDE genes. LFY is a transcription factor that is a key regulator of floral meristem and floral organ identity (Moyroud et al., 2010); it opens up the chromatin around the ABCDE genes, enabling their transcription (Jin et al., 2021). In arabidopsis, UFO interacts with LFY to activate the B class genes AP3 and PI as well as other targets (Rieu et al., 2023b). In some species such as petunia, UFO is also required in addition to LFY to promote floral meristem identity (Souer et al., 2008). *E. peplus* has two homologs of LFY: LFY1_1 is significantly upregulated in both stages of primordia as compared with cyathophylls, and LFY_2 is significantly upregulated in small primordia and filiform structures as compared with cyathophylls (Figure 4, Supplementary File 5). *E. peplus* UFO is upregulated in both stages of primordia, both stages of staminate flowers, young pistillate flowers, and in filiform structures, as compared with cyathophylls. It is significantly upregulated in mature staminate flowers as compared with mature pistillate flowers.

The gene CRABS CLAW (CRC), which is necessary for nectary development in both arabidopsis and petunia (Morel et al., 2018; Slavković et al., 2021), is not upregulated in *E. peplus* nectaries as compared with cyathophylls (Figure 4, Supplementary File 5), suggesting an alternative nectary development pathway in *E. peplus*. It is significantly upregulated in both stages of pistillate flowers as well as in filiform structures compared with cyathophylls, and significantly upregulated in both stages of pistillate flowers compared with staminate flowers. The upregulation of CRC in pistillate flowers in *E. peplus* is consistent with CRC’s other functions in carpel development and meristem termination and with results from *J. curcas*, where CRC is highly upregulated during developmental stages of pistillate flowers (Xu et al., 2016).

### Transcriptomic analysis of differentially expressed genes in cyathium components

We also wanted to investigate what made different parts of the cyathium unique beyond the curated list of floral genes we had selected. Therefore, we performed a gene ontology (GO) term analysis on the genes significantly upregulated in mature staminate flowers, mature pistillate flowers, young staminate flowers, young pistillate flowers, involucres, filiform structures, and nectaries as compared with cyathophylls (Supplementary File 7). We visualized the result for nectaries, involucres, and filiform structures based on semantic similarity of the GO terms (Figure 5).

**Figure 5:**
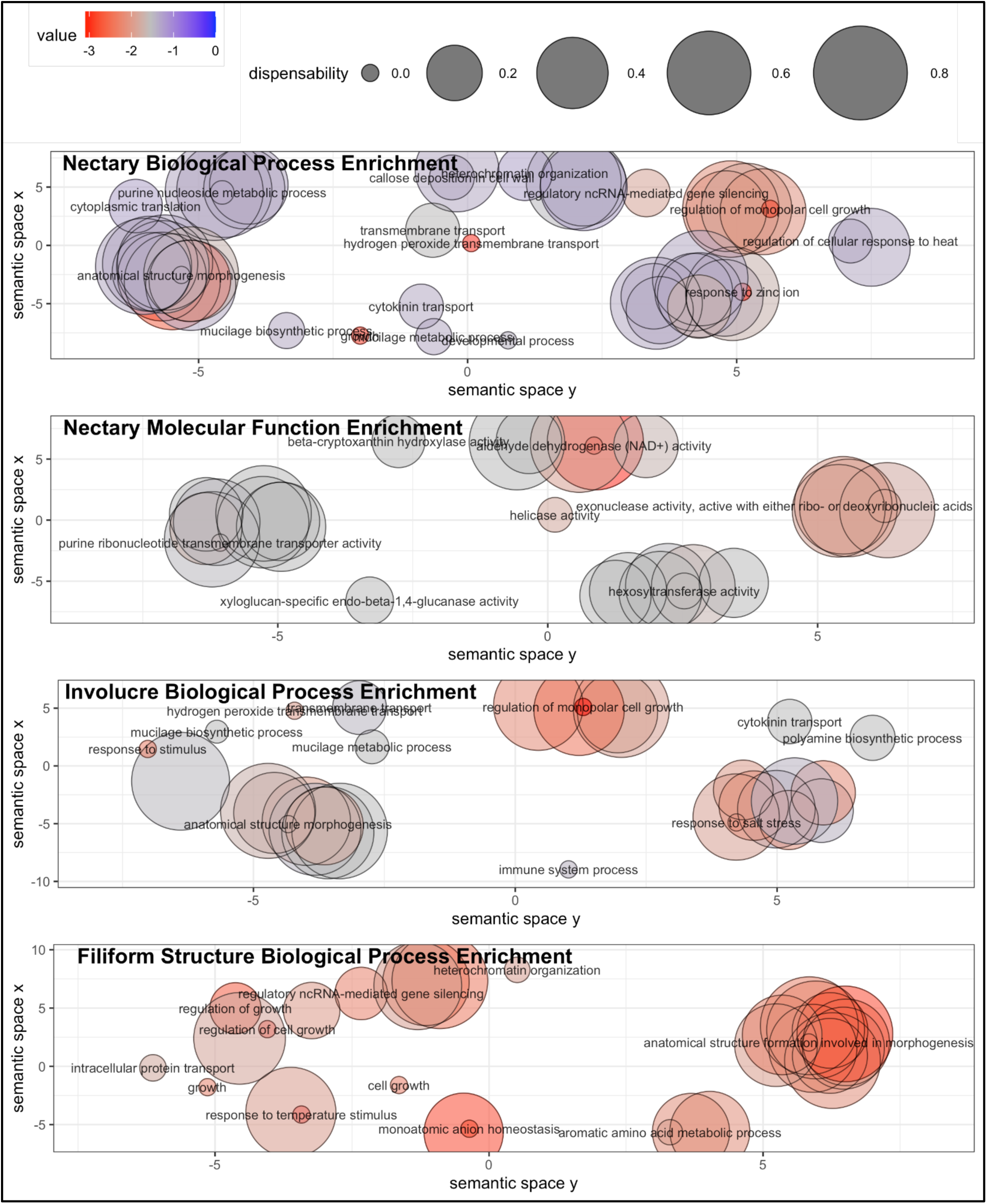
Biological Process GO terms for genes significantly upregulated in the nectary, involucre, and filiform structures as compared with the cyathophyll and Molecular Function GO terms for genes significantly upregulated in the nectary as compared with the cyathophyll are summarized using REVIGO. “Value” is the log10(p-value); a lower value, towards the red side of the color scale, is more significant. The size of the points, the dispensability, represents the semantic similarity to other terms in the dataset.

We found that genes upregulated in the nectary as compared with cyathophylls were significantly enriched for multiple Biological Process GO terms relating to transmembrane transport and specialized metabolism (Figure 5, Supplementary File 7). GO terms relating to gene silencing were also enriched. The Molecular Function GO terms enriched in the nectary included multiple significant terms relating to transmembrane transporter activity and glycosyltransferase activity, similarly to the Biological Process GO terms. ATP and NADH metabolism-related terms were also significantly enriched in the nectaries, suggesting an enrichment in energy-generating redox activities.

Genes upregulated in involucres as compared with cyathophylls were also significantly enriched in genes with Biological Process GO terms relating to specialized metabolite biosynthesis and transmembrane transport (Figure 5, Supplementary File 7). This could potentially indicate that some of the compounds present in the nectaries are at least partially biosynthesized in the involucre tissue connected to the nectaries. The involucre also had several significant GO terms relating to stress, immune responses, and defense responses. The defense GO category may be enriched because of shared hormonal signaling between defense and development; for example, the gene Ep_chr4_g10976, which is upregulated in involucres, is in an orthogroup with AT1G31770(ABCG14), a transmembrane cytokinin transporter. Cytokinin is involved in responses to osmotic stress, salt stress, and other environmental factors but also regulates developmental processes such as inflorescence branching in *J. curcas* (Chen et al., 2019).

The involucre-upregulated gene with the highest p-value, Ep_chr5_g16200, is in an orthogroup with AT5G03680, PETAL LOSS (PTL), which generally represses tissue growth by limiting cell division (Supplementary File 2). (Despite its name, PTL does not regulate petal organ identity, and its expression is not limited to petals– *ptl* mutants’ lack of petals may be due to indirect effects of tissue overgrowth disrupting auxin maxima (O’Brien et al., 2015; Takeda et al., 2022).)

This gene is upregulated in filiform structures, young pistillate flowers, mature pistillate flowers, nectaries, and both stages of primordia as compared with cyathophylls (Supplementary File 5), suggesting functions in patterning across multiple cyathium components.

For the genes upregulated in filiform structures as compared with cyathophylls, stamen development Biological Process GO terms were significantly enriched (Figure 5, Supplementary File 7). Heterochromatin formation, gene silencing, and DNA methylation-related terms were also all significantly enriched.

The mature staminate flowers had multiple significant Biological Process GO terms for stamen identity and morphogenesis as well as ATP and NADPH related terms (Supplementary File 7). The young staminate flowers had multiple significant GO terms associated with stamen and pollen development but did not have significant ATP or NADPH related terms. The mature pistillate flowers had two GO terms associated with petal development; the pistillate flower-enriched gene causing this significant GO hit, Ep_chr1_g00250, is in an orthogroup with arabidopsis AT4G16850, KronbladStorleK (KSK; the name is Swedish for petal size) which influences auxin responses during petal development (Miller et al., 2020). Compared with the mature pistillate flower, the younger pistillate flower was enriched for more GO terms relating to reproductive structure development, suggesting that the patterning of the pistillate flower is more active in this earlier stage. Both stages of both the pistillate and staminate flowers had significant GO terms relating to gene silencing; the top five most significant mature pistillate flower terms were all related to gene silencing.

The zinc finger motif-containing gene Ep_chr4_g09148 was in the top 10 most significant upregulated genes in the involucre, nectary, and filiform structures, and was significantly upregulated in every other cyathium tissue studied compared with the involucre (Supplementary File 5). This gene shares an orthogroup with AT2G33835, FRIGIDA-ESSENTIAL 1 (FES1) which is part of a complex that represses flowering in arabidopsis (Schmitz et al., 2005).

### Evolutionary analysis of ABCDE floral identity gene sequences

Changes in MADS-box binding partners are a driver of floral morphological diversity across many plant families (Bartlett, 2017). For example, duplication and neofunctionalization of SEP genes is involved in the generation of pseudanthia in the Asteraceae (Zhang et al., 2017). Therefore, in addition to analyzing gene expression, we analyzed gene sequence evolution in order to see whether there were changes in *Euphorbia* gene sequence and/or copy number that might reflect changes in binding and function. We performed a codon-aware alignment, tree construction, and dN/dS analysis using the gene sequences from *E. peplus*, *Euphorbia lathyris*, several other Euphorbiaceae species, and the model systems arabidopsis, soybean, and poplar (Figure 6, Supplementary Figure 1, Supplementary Figure 2, Supplementary Figure 3).

**Figure 6:**
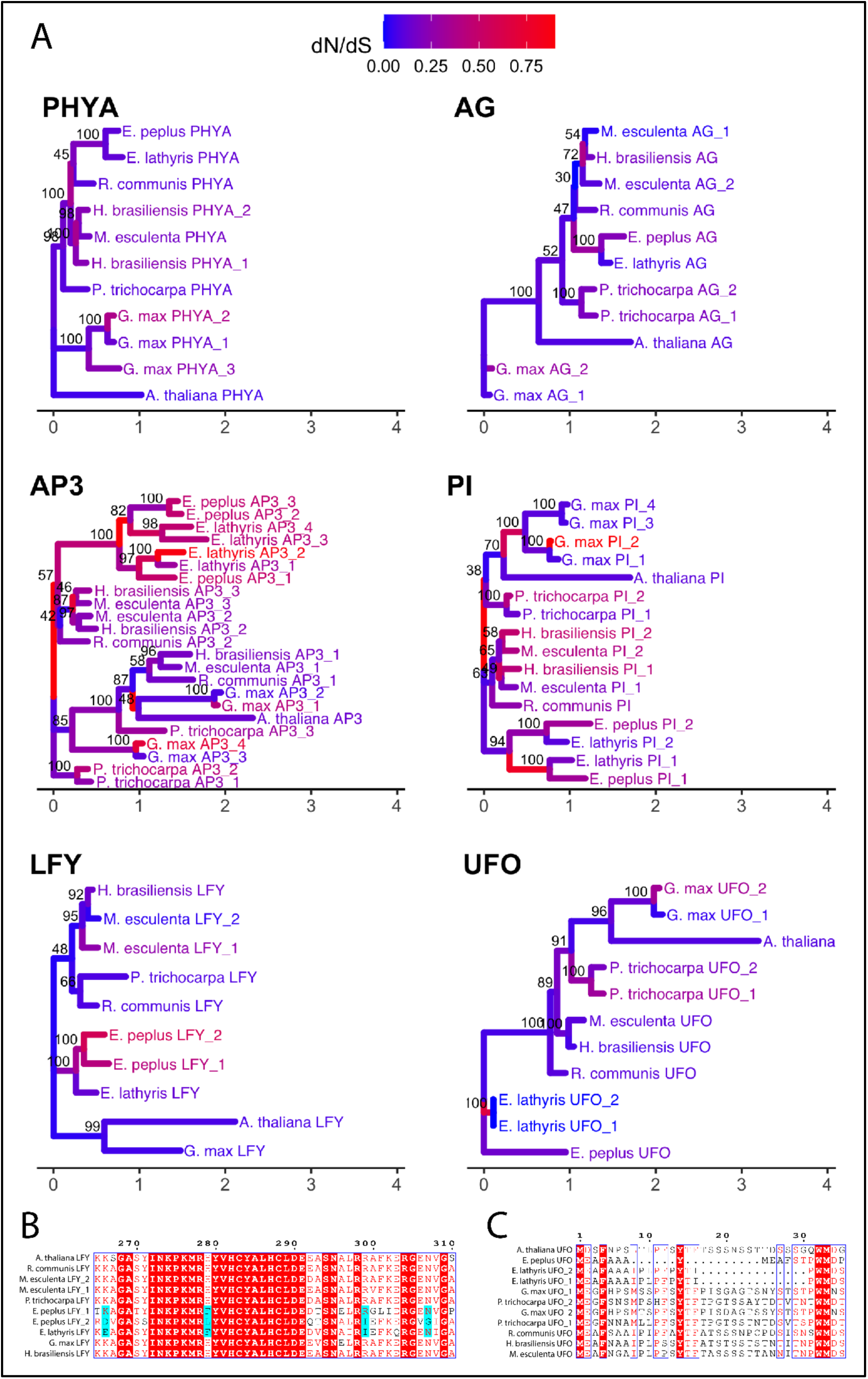
LFY, UFO, and B class genes show signs of divergence and relaxed selection in *Euphorbia*. A. dN/dS values shown on gene trees of floral genes of interest; RAxML support value from 1000 bootstraps is shown at nodes. PHYA is shown as a non-floral “control”. B. Visualization of amino acid alignment in a highly conserved region of LFY; site 279 is a DNA binding site that is highly conserved (except in *Euphorbia*). Cyan color shows sites in the LFY protein alignment rated as “highly significant” for positive selection in the *Euphorbia* clade by PAML. C. Visualization of amino acid alignment at the start of the UFO protein showing deletions in *Euphorbia* (“Ep” and “Casp” genes). Gene names and their gene ID equivalents are listed in Supplementary File 3.

LFY, UFO, AP3, and PI show high sequence divergence in *Euphorbia* (Figure 6A). Compared with the non-floral gene PHYTOCHROME A (PHYA) and with the C class gene AG, the genes LFY, AP3, and PI have high dN/dS on their *Euphorbia* branches. The *E. peplus* and *E. lathyris* homologs for LFY have an amino acid change from a positive amino acid to a hydrophobic amino acid in a DNA-binding site that is otherwise deeply conserved in vascular plant lineages dating back to the lycophytes (Figure 6B) (Rodríguez-Pelayo et al., 2022). The *E. peplus* and *E. lathyris* homologs for UFO also differ greatly in sequence from other Euphorbiaceae homologs of UFO, and one reason is that the *Euphorbia* versions have deletions near the start of the protein (Figure 6C). Interestingly, many of the AP3 residues flagged as “highly significant” for positive selection in *Euphorbia* are in the K domain in a region involved in tetramer formation (Lai et al., 2019), perhaps indicating a change in binding partner (Supplementary Figure 3).

Because LFY and UFO showed a high level of sequence divergence, in order to investigate whether LFY and UFO binding targets were conserved, we looked for LFY binding motifs and LFY-UFO binding motifs in the putative promoters of homologs of the LFY targets AP3, PI, AG, and AP1 in *E. peplus*, *H. brasiliensis*, and arabidopsis. (AP3 and PI are targets of the LFY-UFO complex as well.) We hypothesized that the binding motifs might not be conserved in *E. peplus*. Both canonical LFY binding motifs and UFO-LFY motifs (mLUBS and dLUBS) are present in the 3kb upstream of these target genes in *E. peplus* (Supplementary File 9), suggesting that at least some of the binding motifs of LFY and LFY-UFO are conserved despite the high divergence in sequence.

## DISCUSSION

Few pseudanthia have been well-characterized at the molecular level. Several studies have examined transcriptional patterns and key developmental genes in the pseudanthium of gerbera (*Gerbera hybrida*, Asteraceae), which has a flower unit meristem (FUM) that lacks apical growth, initially lacks visible primordia, and fractionates into many flowers in a short developmental span—similarly to *Euphorbia*’s cyathium FUM. In the ABCDE model, E-function SEP genes generally promote floral identity. Gerbera has eight SEP homologs due to lineage-specific gene duplications in both the SEP1 and SEP3 subclades; and genes in both subclades have functionally diversified in terms of which floral organs they help specify in the gerbera pseudanthium (Zhang et al, 2017). Interestingly, *Euphorbia* only has four SEP homologs and they are not the result of *Euphorbia*-specific duplications (Supplementary Figure 1), indicating that SEP gene family duplication may not have contributed as much to the development of a pseudanthium in *Euphorbia* as in the Asteraceae. However, the *Euphorbia* SEP homologs do show distinct expression patterns within the cyathium, with the three *Euphorbia* SEP1 homologs all showing different expression patterns from each other. This is consistent with a broader trend that SEP1 orthologs have been repurposed in many species to regulate bract identity and inflorescence architecture (Dreni and Ferrándiz, 2022).

LFY and UFO are generally important in specifying floral meristem identity (Rieu et al., 2023a; Weigel et al., 1992; Levin and Meyerowitz, 1995, Lippman et al., 2008), although this is not necessarily true for pseudanthia. in gerbera, GhLFY is uniformly expressed in the undifferentiated FUM and regulates its determinacy, while GhUFO is expressed in emerging flower primordia and regulates floral meristem identity (Zhao et al, 2016). The LFY homolog in *J. curcas* (Euphorbiaceae) does promote floral meristem identity; its overexpression causes early flowering, and its co-suppression delays flower formation, promotes more inflorescence branching, and the formation of sepal-like organs (Tang et al., 2016). Although our dataset lacks the spatial and temporal resolution needed at the earliest time points to distinguish which genes are expressed in the cyathium FUM as opposed to the floral meristems, both LFY and UFO are expressed highly in our small primordium and large primordium stage suggesting that they are both important in floral meristem and/or cyathium FUM identity.

In our analysis of gene sequence evolution, *Euphorbia* LFY, UFO, and the B-class genes AP3 and PI show signs of divergence based on phylogenetic distance and relaxed selection. Except for UFO (which may have lacked signs of relaxed selection due to the gene tree topology), these genes also had multiple sites that were classified as highly significant for positive selection in Bayes Empirical Bayes (BEB) analysis (Supplementary File 8). The increased rates of molecular evolution of AP3 and PI in *Euphorbia* is consistent with increased rates of molecular evolution of B class genes in other lineages that have lost petals, for example the Piperaceae and Saururaceae compared with other Piperales (Jaramillo and Kramer, 2007). This likely reflects a release of evolutionary constraints when B class genes no longer need to promote petal identity.

One interesting observation regarding the expression of UFO is that in *J. curcas*, UFO is downregulated in both mature staminate flowers and mature pistillate flowers compared to younger stages (Chen et al., 2016), whereas in our *E. peplus* dataset we saw the highest expression of UFO in mature staminate flowers. This either suggests a repurposing of UFO, or that developmentally mature but not reproductively mature (i.e. not elongated) staminate flowers in the *E. peplus* cyathium aren’t yet transcriptomically mature compared with pistillate flowers.

As UFO functions in the development of individual perianth organs as well as in the broader activation of the B class genes (Takeda et al., 2022), some of the changes in UFO expression and sequence in *Euphorbia* could be related to the suppression of perianth development in addition to potential functions in defining the floral meristem.

Genes in the AP1 clade can also be involved in floral meristem specification– and are also highly diverged in *Euphorbia*. In arabidopsis, AGL79 interacts with SUPPRESSOR OF OVEREXPRESSION OF CONSTANS1 (SOC1) to regulate the expression of TERMINAL FLOWER 1 (TFL1) which controls inflorescence meristem identity (Yang et al., 2023). This AP1 (AGL79) homolog in *Euphorbia* appears to have diverged, and now groups in a separate clade within the AP1 clade “missing” its AP1-CAL homolog in the Euphorbiaceae AGL79 clade in our gene tree because that gene, AP1_2, is so diverged that it appears elsewhere in the tree (Supplementary Figure 2). One question we had about the cyathium was which other organs might have homology with the filiform structures. In the filiform structures, we observed significant expression of B-class and E-class genes, as well as one homolog of LFY, potentially consistent with floral identity. The filiform structures had low levels of RNA overall compared with other structures in the cyathium (Supplementary File 1), which could be explained by the gene ontology analysis showing significant GO terms relating to heterochromatin formation. Our data therefore agrees with a potential interpretation of the filiform structures as reduced flowers in an inflorescence. Overall, our GO term analysis showed that gene silencing seems to be involved in the creation of multiple cyathium organs.

The C class gene AGAMOUS is highly expressed in both pistillate flowers and staminate flowers, as would be expected given its function in pistillate and staminate development. We did not see transcriptomic signs of abortion of other whorls in the pistillate and staminate flowers (e.g. hypothetically, seeing B class expression in the young pistillate flower could have been an example of that), besides the upregulation of the putative homolog of the arabidopsis perianth gene KSK in the pistillate flower. Organ abortion can happen at many different stages, however, and unisexual flowers do not always show evidence of their missing whorls (Mitchell and Diggle, 2005).

In future studies, it would be fruitful to expand this analysis to closely related species given Prenner and Rudall’s interpretation of the Australian genus *Calycopeplus* and the New Caledonia tree *Neoguillauminia cleopatra* as “missing links” towards the *Euphorbia* cyathium; future studies could sequence the floral genes of *Calycopeplus*, *Neoguillauminia*, and *Anthostema* to study the sequence evolution of LFY, UFO, AP1 and B class genes with those taxa included to see how much of the sequence divergence occurred in *Euphorbia* alone and how much is shared with these other taxa.

In addition to expanding our analyses to more taxa, future studies could experimentally investigate the function of the UFO-LFY complex and its DNA binding targets in *Euphorbia*, e.g. through electrophoretic mobility shift assays (EMSA), microscale thermophoresis (MST), or DAP-seq. Additionally, yeast two-hybrid or similar techniques could be used to check the transcription factor binding partners of *Euphorbia*’s B-class genes. It would also be interesting to use ATAC-seq or another method to determine chromatin accessibility across the genome and speculate on what might be being silenced in the different cyathium structures. *In situ* hybridization, if optimized in this system, could also potentially reveal more spatially and temporally resolved patterns of gene expression, including more detailed expression patterns of LFY and UFO. We are excited to see what future work reveals about the developmental genetics involved in creating the complex structure of the cyathium.

## Supporting information

All Supplemental Files

## ACKNOWLEDGEMENTS

Thank you to Jacob Landis for extremely helpful advice on running PAML. Thank you to François Parcy, Jérémy Lucas and Laura Turchi for graciously sending the motif files for mLUBS and dLUBS. Thank you to the Cornell Weill growth chamber plant maintenance staff, especially Corbin Lay, for their excellent upkeep of the plants used. Thank you to the Frank Lab for very helpful comments on the manuscript. This work was funded by a Cornell Institute of Biotechnology Seed Grant, National Science Foundation Integrative Organismal Systems (NSF IOS) award 1942437, and startup funds from the College of Agriculture and Life Sciences at Cornell University to Margaret Frank.

## AUTHOR CONTRIBUTIONS

A.R.J., A.B., and M.H.F. all conceived of the study. Under the supervision of A.R.J., A.B. collected the images of cyathia and dissected cyathia components, and performed preliminary dissections and RNA extractions that contributed to the experimental design for the RNAseq experiment. A.R.J. performed the dissection for the RNAseq experiment. A.B. and A.R.J. both contributed to the RNA extraction for the RNAseq experiment. A.B. and A.R.J. collaborated on the initial sequence alignment and phylogeny construction to assess conservation of sequence similarity in floral genes. A.R.J. performed all other data analysis. A.R.J. wrote the first draft of the manuscript, after which all authors contributed to reviewing and editing the manuscript.

## DATA AVAILABILITY

The raw RNAseq data generated in this study are available from NCBI GenBank under the accession PRJNA1063728. All code used in the data analysis is available on Github at https://github.com/ariellerjohnson/Cyathium-2024.

