## Supplementary material for "A transcriptional dissection of the petty spurge (*Euphorbia peplus* L.) reproductive structure": All Supplemental Files: Supplementary Figures.pdf

Supplementary Figure 1

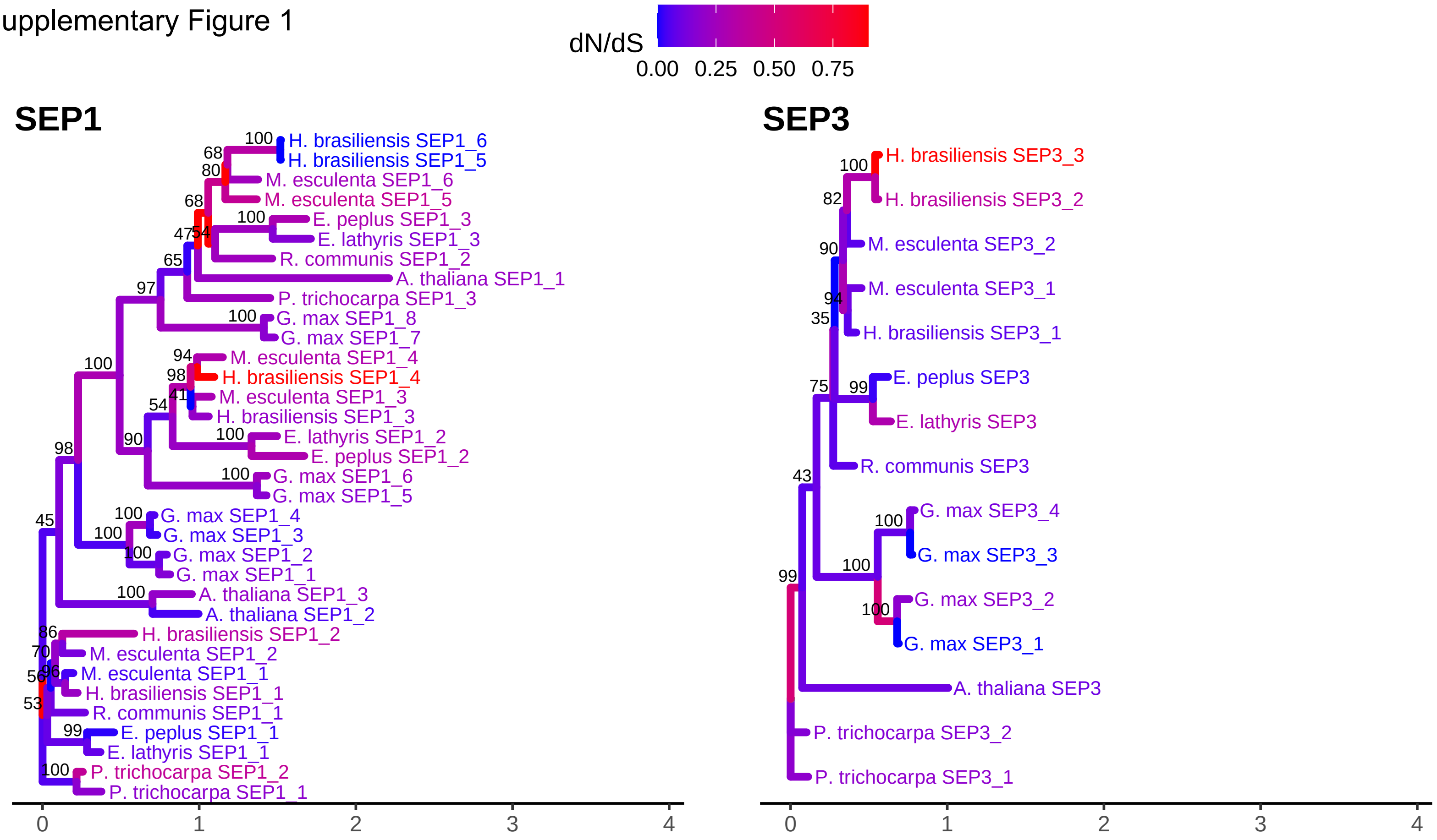

Supplementary Figure 2

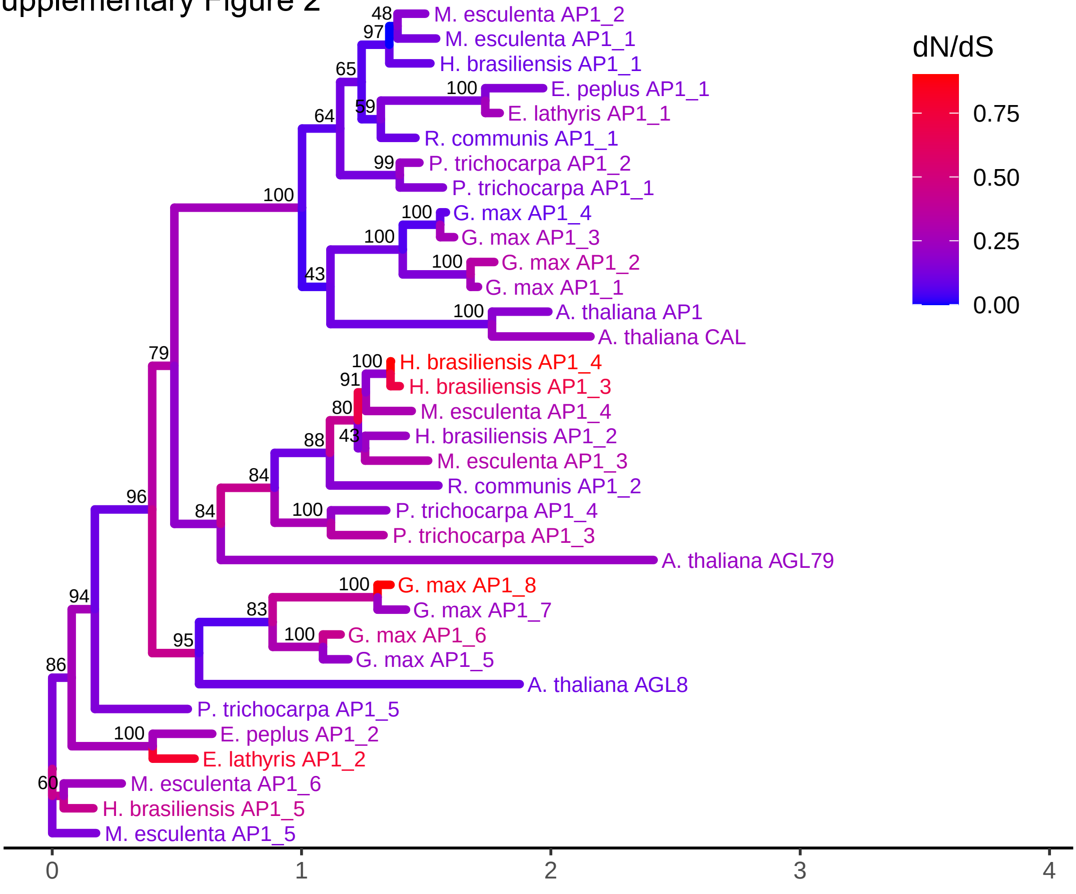

Supplementary Figure 3

# AG

```
                        1        10        20        30        40        50        60
Casp22688.t1            MSYPSDSRETSPQRKMGRGKIEIKRIENTTNRQVTFCKRRNGLLKKAYELSVLCDAEVAL
XP_015576689.1          MEYQGDSRDDSPQRKMGRGKIEIKRIENTTNRQVTFCKRRNGLLKKAYELSVLCDAEVAL
Ep_chr4_g09342.t1       MSYPSDERDESPQRKLGRGKIEIKRIENTTNRQVTFCKRRNGLLKKAYELSVLCDAEVAL
KAF2314627.1            ...............MGRGKIEIKRIENTTNRQVTFCKRRNGLLKKAYELSVLCDAEVAL
Manes.17G062000.1.p     MAYQSESRETSPQRKMGRGKIEIKRIENTTNRQVTFCKRRNGLLKKAYELSVLCDAEVAL
Manes.15G115500.3.p     MAYPSNSGETSLQRKIGRGKIQIKRIENTTNRQVTFCKRRNGLLKKAYELSVLCDAEVAL
Potri.011G075800.2.p    MAYQNEPQESSPLRKLGRGKVEIKRIENTTNRQVTFCKRRNGLLKKAYELSVLCDAEVAL
Potri.004G064300.3.p    MEYQNESLESSPLRKLGRGKVEIKRIENTTNRQVTFCKRRSGLLKKAYELSVLCDAEVAL
AT4G18960.1             TAYQSEGGDSSPLRKSGRGKIEIKRIENTTNRQVTFCKRRNGLLKKAYELSVLCDAEVAL
Glyma.15G088600.1.p     MVFPNPSMSVSPQKKMGGGKIEIKRIENTTNRQVTFCKRRNGLLKKAYELSVLCDAEVAL
Glyma.13G223300.1.p     MAFPDPSMSVSPQRKMGRGKIEIKRIENTTNRQVTFCKRRNGLLKKAYELSVLCDAEVAL

                                 70        80        90       100       110       120
Casp22688.t1            IVFSSRGRLYEYANNSVKTTIERYKKATADASNTGSVSEANVQYYQQESAKLRQQINSLQ
XP_015576689.1          IVFSSRGRLYEYANNSVRSTIDRYKKVSTDSSTTGSVSEANAQFYQQEAAKLRVQIGNLQ
Ep_chr4_g09342.t1       IVFSSRGRLYEYANNSVKTTIDRYKKATTDASSTGSVSEANVQYYQQESDKLRKEITNLQ
KAF2314627.1            IVFSSRGRLYEYANNSVKSTIERYKKACADSSSTGSVSEANAQYYQQEAAKLRVQISNLQ
Manes.17G062000.1.p     IVFSSRGRLYEYANNSVKSTIERYKKACADSSNTGSVSEANAQYYQQEAAKLRVQISNLQ
Manes.15G115500.3.p     IVFSTRGRLYEYANNSVKSTIERYKKASADSSNTGSVSEVNAQFYQQEAAKLRVQISNLQ
Potri.011G075800.2.p    IVFSSRGRLYEYSNNSVKSTIERYKKACADSSNNGSVSEANAQFYQQEAAKLRSQIGNLQ
Potri.004G064300.3.p    IVFSSRGRLYEYSNDSVKSTIERYKKASADSSNTGSVSEANAQYYQQEAAKLRSQIGNLQ
AT4G18960.1             IVFSSRGRLYEYSNNSVKGTIERYKKAISDNSNTGSVAEINAQYYQQESAKLRQQIISIQ
Glyma.15G088600.1.p     IVFSSRGRLYEYANNSVKATIERYKKACSDSSGAGSASEANAQFYQQEADKLRAQISSLQ
Glyma.13G223300.1.p     IVFSSRGRLYEYANNSVKATIERYKKASSDSSGAGSASEANAQFYQQEADKLRQQISNLQ

                                130       140       150       160       170       180
Casp22688.t1            THNRQMMGEGIGSMHQRELKSLESRLEKGISRIRSKKNELLFAEIEYMQKREIDLHNNNQ
XP_015576689.1          NSNRHMLGESLGALNLKDLKGLEGRLEKGISRIRSKKNELLFAEIEYMQKREIDLHNNNQ
Ep_chr4_g09342.t1       ARNREMMGEGIGSMQHRDLKSLEGKLEKGINRIRSKKNELLFSEIEYMQKREVDLHNNNQ
KAF2314627.1            NSNGNMLGESLGALNVKELKSLEIRLEKGISRIRSKKNELLFAEIEYMQKREIDLHSNNQ
Manes.17G062000.1.p     NSNRHMLGESLGALTVKELKSLEIRLEKGISRIRSKKNELLFAEIEYMQKREIDLHNNNQ
Manes.15G115500.3.p     NSNRHMLGESLGALNVKELKSLEIRLEKGISRIRSKKNELLFAEIEYMQKREIDLHNNNQ
Potri.011G075800.2.p    NSNRNMLGESLSALSVKELKSLEIKLEKGIGRIRSKKNELLFAEIEYMQKREIDLHNNNQ
Potri.004G064300.3.p    NSNRHMLGEALSSLSVKELKSLEIRLEKGISRIRSKKNELLFAEIEYMQKREVDLHNNNQ
AT4G18960.1             NSNRQLMGETIGSMSPKELRNLEGRLERSITRIRSKKNELLFSEIDYMQKREVDLHNDNQ
Glyma.15G088600.1.p     NNNRQMMGESLGPLTAKELKNLETKLEKGISRIRSKKNELLFAEIEYMQKREIDLHNNNQ
Glyma.13G223300.1.p     NNNRQMMGDSLGSLTAKDLKNLETKLEKGISRIRSKKNELLFAEIEYMQKREIDLHNNNQ

                                190       200       210       220       230       240
Casp22688.t1            LLRAKIAENERKQQNMNLMPGSDNYEMIQSQTYDNRTFFQVNSLQPTNHYSHQDQMSLQL
XP_015576689.1          LLRAKIAENERK.QNMNLMPGGGNYEIMQSQPFDNRNYFQVNALQSTNHYPHQDQMALQL
Ep_chr4_g09342.t1       LLRAKIAENERKQQNMNLMPGSSNYEMIQSQTYDNRTFFQVNSLQPTNHYSHQDPMALQL
KAF2314627.1            LLRAKIAENERKQQNMNMMPGGGNYEITQSQTFDNRNYFQVNALQPTNHYPHQDQMALQI
Manes.17G062000.1.p     LLRAKIADNERKQQNMNMMPGGGNYEIMQSQPFDNRNYFQVNALQPTNHYPHQDQMALQL
Manes.15G115500.3.p     LIRAKIAENERKQQNMNLMPGGGDYEMMQPQPFDNRNYFQVNALQPTNHYPHQDQMALQL
Potri.011G075800.2.p    LLRAKIAENERKRQHMNLMPGGVNFEIMQSQPFDSRNYSQVNGLPPANHYPHEDQL...F
Potri.004G064300.3.p    LLRAKISENERKRQSMNLMPGGADFEIVQSQPYDSRNYSQVNGLQPASHYSHQDQMALQL
AT4G18960.1             ILRAKIAENERNNPSISLMPGGSNYEQLQSQPFDSRNYFQVAALQPNNHHGRQDQTALQL
Glyma.15G088600.1.p     LLRAKIAEGERNHHNLAVLPGGSNYDSLQTQQFDSRGYFQVTGLQPNNQYARQDQMSLQL
Glyma.13G223300.1.p     LLRAKIAESERNHHNMAVLPGGSNYDSMQSQQFDSRGYFQVTGLQPNNQYARQDQMSLQL

Casp22688.t1            VX
XP_015576689.1          VX
Ep_chr4_g09342.t1       VX
KAF2314627.1            VX
Manes.17G062000.1.p     VX
Manes.15G115500.3.p     VX
Potri.011G075800.2.p    SX
Potri.004G064300.3.p    VX
AT4G18960.1             VX
Glyma.15G088600.1.p     VX
Glyma.13G223300.1.p     VX
```

# AP1

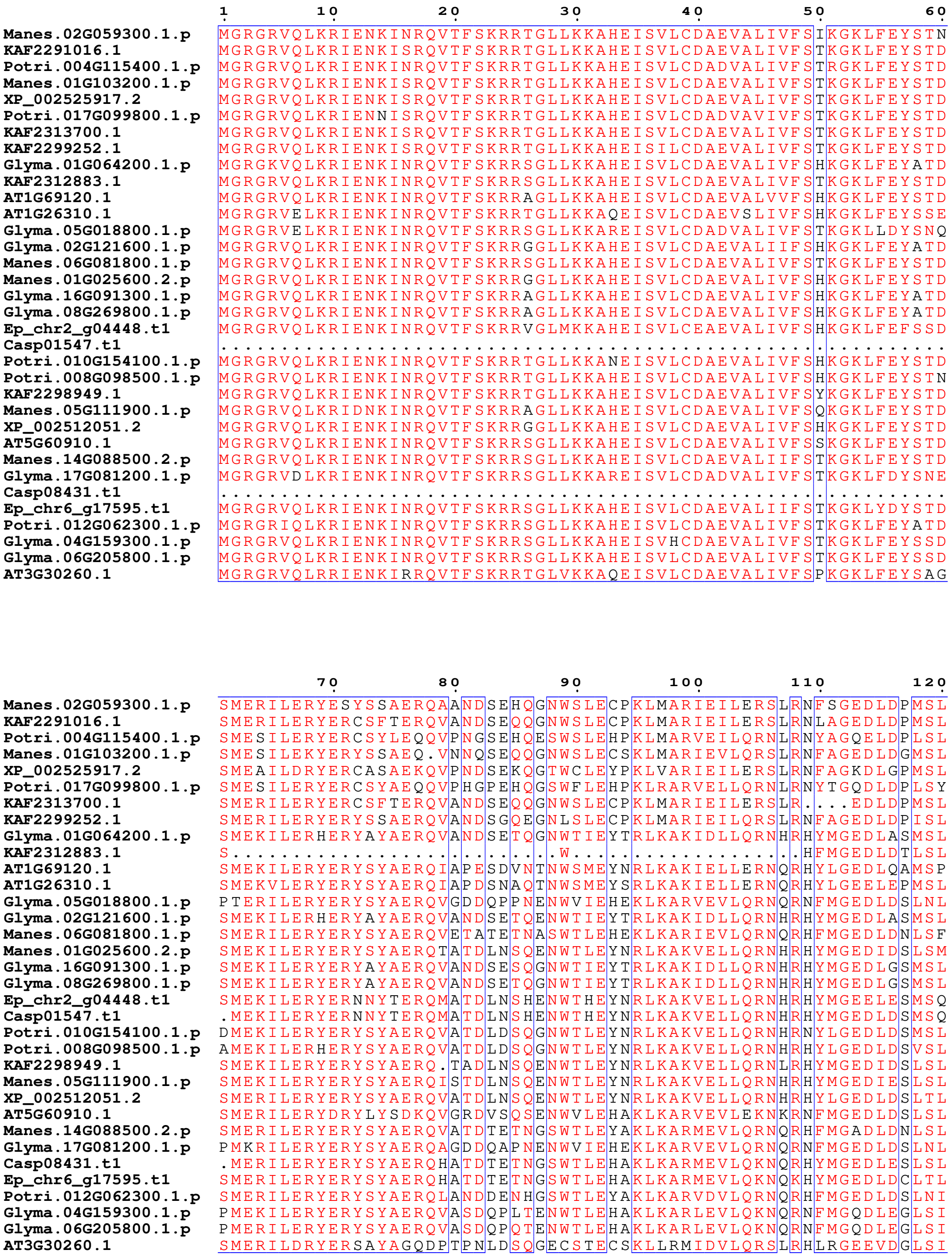
Manes.02G059300.1.p
KAF2291016.1
Potri.004G115400.1.p
Manes.01G103200.1.p
XP_002525917.2
Potri.017G099800.1.p
KAF2313700.1
KAF2299252.1
Glyma.01G064200.1.p
KAF2312883.1
AT1G69120.1
AT1G26310.1
Glyma.05G018800.1.p
Glyma.02G121600.1.p
Manes.06G081800.1.p
Manes.01G025600.2.p
Glyma.16G091300.1.p
Glyma.08G269800.1.p
Ep_chr2_g04448.t1
Casp01547.t1
Potri.010G154100.1.p
Potri.008G098500.1.p
KAF2298949.1
Manes.05G111900.1.p
XP_002512051.2
AT5G60910.1
Manes.14G088500.2.p
Glyma.17G081200.1.p
Casp08431.t1
Ep_chr6_g17595.t1
Potri.012G062300.1.p
Glyma.04G159300.1.p
Glyma.06G205800.1.p
AT3G30260.1

```
                              130       140       150       160       170       180
Manes.02G059300.1.p    RELQHLEQQIANGLKRVRARKNQLYHESISGLQKKERALHEQNNMLSKKLKENEKTTAEQ
KAF2291016.1           RELQHLEQQIDNGLKRVRARKNQLYNQSISELQKKERALHEQNNMLSKKVKENEKTMAEQ
Potri.004G115400.1.p   KELQYLEQQIDTALKRIRSRKNQLIHESLNELRKKEKELQEQNNILAEQVKENEKSLTEQ
Manes.01G103200.1.p    RELQHLEQQIDTGLKRVRTRKNQLYNESISELQKKEKALHEQNNMLSKQLEQNEKTMAEG
XP_002525917.2         RELQHLEQQIDTALKRVRSRKNQLYHESLAALQKNERSLHEQNNMLAKQLKENEKTLTER
Potri.017G099800.1.p   KELQHLEQKIDTALKSVRSRKNQLVHESLAEMQKKEKALQDQNNILVEQVKKKLKALTEQ
KAF2313700.1           RELQHLEQQIDNGLKRVRARKNQLYNQSISELQKKERALHEQNNMLSKS...........
KAF2299252.1           RELQHLEQQIDNGLKRVRARKNQLYHESLSELQKKDKALHEQNSMLSKQLKQNEKSMAEH
Glyma.01G064200.1.p    KELQSLEQQLDTAIKNIRTRRNDLMYASISELQKKEKMIQEQNNILAKKIKEKEQAVAQQ
KAF2312883.1           KELQSVEQQIDSALKHIRSRKNQLMYESIAELQKK..............VKEKEKEIAQQ
AT1G69120.1            KELQNLEQQLDTALKHIRTRKNQLMYESINELQKKEKAIQEQNSMLSKQIKEREKILAQQ
AT1G26310.1            KDLQNLEQQLETALKHIRSRKNQLMNESLNHLQRKEKEIQEENSMLTKQIKERENILTKQ
Glyma.05G018800.1.p    RGLQSLEQQLDSALKHIRSRKNQAMNESISELQKKDRTLREHNNLLSKKIKEKEKELTPQ
Glyma.02G121600.1.p    KELQSLEQQLVTGIKNIRTRRNDLMSESISELQKKEKRIQEENNTLAKKIKEKEQAVAQQ
Manes.06G081800.1.p    KELQSVEHQIDSALKHIRTKKNQLMYESISELQKKDKALQEQNNQLAKKVKEKEKEIAQQ
Manes.01G025600.2.p    KELQNLEQQLDTALKHIRTRKNQLMYESISELQKKEKAIQEQNNMLAKQIKENEKAVAQQ
Glyma.16G091300.1.p    KELQSLEQQLDTALKQIRTRRNQLMYESISELQKKEKVIQEQNNMLAKKIKEKEKVAAQQ
Glyma.08G269800.1.p    KELQSLEQQLDTALKQIRTRRNQLMYESISELEKKEKVIQEQNNMLAKKIKEKEKVAASQ
Ep_chr2_g04448.t1      KELGNLEQQLENALKHIRLRKNQLLADYITDLQKKEKAILEENNMLEKQLKDNQKAVAQQ
Casp01547.t1           KELANLEQQLENALKHIRSRKNQLIGDYISELQKKEKAILEENSMLTKQIKEKEKAVAQQ
Potri.010G154100.1.p   KELQNLEQQIDTALKHIRARKNHLMSQSISELQRKEKAIQVQNNMLVKQIKEKEKALAQP
Potri.008G098500.1.p   KELQNLEQQIDTALKLIRERKNHLMYQSISELQIKEKAIKEQNNMLVKQIKEKEKALAQP
KAF2298949.1           KELQNLEQQLETALKHIRTRKNQLMYESISELQKKEKAIQEQNSMLAKQIKENEKAVAQQ
Manes.05G111900.1.p    KELQNLEQQLDTALKHIRARKNQLMYESISELQKKEKAIQEQNSMLTKQIKEKEKAVAQQ
XP_002512051.2         KELQNLEQQLDTALKHIRTRKNQLMFESISELQKKEKAIQEQNNMLSKQIKEKEKAVAQQ
AT5G60910.1            KELQSLEHQLDAAIKSIRSRKNQAMFESISALQKKDKALQDHNNSLLKKIKEREKKTGQQ
Manes.14G088500.2.p    KELQSLEQQIDSALKHIRSRKNQMMHESISELQKKDKALQDQNNQLAKKVKEKEKEMAQQ
Glyma.17G081200.1.p    RGLQSLEQQLDSALKLIRSRKNQAMNESISALQKKDKSLREHNNLLSKKIKDKEKELAPQ
Casp08431.t1           KDLQSLEQQIDSALKHIRSRKNNLMAENLTDLQKKDKIMEEERNQLLKKVKEKEKEIAAQ
Ep_chr6_g17595.t1      KELQSLEQQIDSALKHIRSRKNQLLYEQLTDMQKRDKLMEEANKQLHRQVKEKEKELAEQ
Potri.012G062300.1.p   KELQNLEHQIDSALKHVRSRKNQLMYESISELQKKDKALQEQNNMLAKKVKEKEKAIAQQ
Glyma.04G159300.1.p    KELQNLEHQLESALKHIRSRKNQLMYESISELHKKDKALQEQNNTLAKKIKEKEKALAQQ
Glyma.06G205800.1.p    KELQNLEHQLDSALKHIRSRKNQIMHESISELHKKDKVLQEQNNTLAKKIKEKEKALAQH
AT3G30260.1            RDLQGVEMQLDTALKKTRSRKNQLMVESIAQLQKKEKELKELKKQLTKKAGEREDFQTQN
```

```
                              190       200       210       220
Manes.02G059300.1.p    ANCEQQNGPNTSFMPLLSISGPNENEEARAQIHPDTLPPWMLHVNE
KAF2291016.1           ANWEQQNGQNTSFMPLLTIGGTNEDEEAIAQIQPNTIPPWMLHVNE
Potri.004G115400.1.p   AQWEQRNGQNSSFMPMLTIGDSNGNENVEVQTPPSTMPSWMLHVND
Manes.01G103200.1.p    SNWEQQNGQNTSFMPLPAIGGTKENEEGMAQEEPNTMPSWMLHVNE
XP_002525917.2         EQWEQQNSQNAAFFPLPTTCGNNENEESTAHAQPNTMPPWMLHLNE
Potri.017G099800.1.p   AQWEQQNGQNSSFMPLPTIGGSNGNKDVEVQTQPSTMPHWMLHVND
KAF2313700.1           ..................ASGTNEDEEAIAQIQPNTIPPWMLHVNE
KAF2299252.1           ANWEQRNGQNTYFMPLLTTGSTNEKEEATAQIQLNTMPPWMLHVNE
Glyma.01G064200.1.p    AQWEQPNRVDTSFMPLLNIGGNEAAPELGRNGLDLTLPLYSCHLGC
KAF2312883.1           NQKEQQNIIDSTVLPMLNIRGNDEDETTPMQNLANAVPSWMIYLNE
AT1G69120.1            EQWDQQNGHNPPPLPSLNMGGLDDPMAMRRNDLELTLPVYNCNLGC
AT1G26310.1            TQCEQLNSVDQPFQTSLNMGGLEDQTAMRRNNLDLTLPIYN.YLGC
Glyma.05G018800.1.p    EQEGLQNNMDSSVLPLLTIGGSKSNEETPTSCRPKTIPPLMPPTNE
Glyma.02G121600.1.p    AQWEQPNRVDTSFMPLLNIGNNEAAPELGRNGLDLTLPLYSCNLGC
Manes.06G081800.1.p    TQKEQQSTSVSSTIPMLNLRGNGEDETTPMQHRSNTLPAWLLHMNE
Manes.01G025600.2.p    ALWEQHNGTNSPFLPLLNIGGNEAPEAARRNELDLTLPIYSCHLGC
Glyma.16G091300.1.p    AQWEHPNGVNASFLPLLNMGGNEEASEVGRNELDLTLPLYSCHLGC
Glyma.08G269800.1.p    AQWEHPNGVNASFLPLLNMGGNDEAPEVGRHELDLTLPLYSCHLGC
Ep_chr2_g04448.t1      AQWEQQNNNNSPFLPLLNMGGNEAVPEGRRSELDLTLPIYSCHLGC
Casp01547.t1           AQWEQQNNNNSQFLPLLNIGGNEAVPEVRRNELDLTLPIYSCHLGC
Potri.010G154100.1.p   AFWDLQDGPNSSFLPLLNIGGSEEAPEARRNELGHTLPIYSFHLGG
Potri.008G098500.1.p   ALWDQQDGPNSSFLPLLNI..SEEDPEARRNELDLTLPIYSCHLGC
KAF2298949.1           ALWEQHNGTNSPFLPLPNIGX.........................
Manes.05G111900.1.p    ALWEQHNGTNSPFLPLLNIGVNEEAPEVRRNELDLTLPLYSCHLGC
XP_002512051.2         ALWEQQNGNNSPFLPLLNIGGTEEGQEVRRNELDLTLPIYSCHLGC
AT5G60910.1            E..GQLVCSNSSVLYCSSRDGFGENGGASSLTEPNSLPAWMLTTNE
Manes.14G088500.2.p    TQKEQQNGADSTVLP.FILRGNGEDETTPIQDRGNAVPAWMLHLNE
Glyma.17G081200.1.p    EQDGLQNNMDTSVLPPLTIGGFKCNEETPTSSRPKTIPPWMPPTNE
Casp08431.t1           MDWEQQNGVDSPPLPSANISSSGEEDRTGVQQRGNVLPAWMLHLNX
Ep_chr6_g17595.t1      TEREQQNSADSSSLPIPNTSSAGEDERIGVQQRGNVLPAWMLHLNX
Potri.012G062300.1.p   ASWEQQNDLDPTILPMLNISSSIEEEPAPIQHRANALPAWMLYLNE
Glyma.04G159300.1.p    AQLERLGEVDSSALPLSNIRTSGDNEGTPTPTQANAIPPWMLPTNE
Glyma.06G205800.1.p    AQMEQRGEMDSSALPLSNIRESGDNEGTPTPTRANAIPSWMLPSNE
AT3G30260.1            LSHDLASATP.PFEPPLSSGDTGEVAAGTLIRRTNATPHWMPLTGE
```

# AP3

```
                     1         10        20        30        40        50        60
Manes.02G100400.1.p  MGRGKIEIKRIENATNRQVTYSKRRNGIFKKAQELTVLCDAKVSLIMFSNTGKFHEFISP
Glyma.01G169600.1.p  MGRGKIEIKLIENPTNRQVTYSKRRNGIFKKAHELSVLCDAKVSLIMFSKNNKMHEYISP
KAF2290821.1         MGRGKIEIKRIENPTNRQVTYSKRRNGLFKKARELTVLCDAKVSLIMFSNTGKFHEFISP
Potri.005G118000.2.p MGRGKIEIKKIENPTNRQVTYSKRRNGIFKKAQELTVLCDAKVSLIMFSNTNKFHEYISP
AT3G54340.1          MARGKIQIKRIENQTNRQVTYSKRRNGLFKKAHELTVLCDARVSIIMFSSSNKLHEYISP
Glyma.04G027200.1.p  MARGKIQIKRIENNTNRQVTYSKRRNGLFKKANELTVLCDAKVSIIMFSSTGKLHQYISP
Glyma.06G027200.1.p  MARGKIQIKRIENTTNRQVTYSKRRNGLFKKANELTVLCDAKVSIIMFSSTGKLHEYISP
KAF2310046.1         MGRGKIQIKRIENSTNRQVTYSKRRNGLFKKANELTVLCDAKVSIIMFSSTGKLHEYVSP
Rc02T004810.1        MARGKIQIKRIENSTNRQVTYSKRRNGLFKKAHELTVLCDAKVSIIMFSSNDKLRQYISP
Casp07377.t1         MGRGKIEMKKIDNATNMQVTFSKRRSGIFKKAEQLSVLCDAQISLIMSSSNNKFYEFVTP
Casp07378.t1         M.......KKIDNAANMEVTFSKRRSGIFKQAEILSLLSDAQISLIMCFSNNKFYDFVTP
Ep_chr8_g24861.t1    MGRGKVLYKKIENPTSLQVTFSKRRSGIFKKAHELSVLCDAQISIIMRSSTGKFYACVSP
Ep_chr8_g24862.t1    MGRGKVMYKKIENPTSLQVTFSKRRSGIFKKAHELSVLCDAQISIIMHSSTGKFYECVSP
Casp10977.t1         MGRGKIAYKKIDNPTSMQI..................C.....LIMRSSTGRFYEFTTP
Ep_chr8_g24863.t1    MGRGKIEMKRIENTTNKQVTFSKRRNGLLKKAQELSILCDADISLLVCSSTSKFSEFHTP
Casp07379.t1         MGRGKIVYKKIDNPTSMQVTFSKRRSGIRKKAEELSILCDAQICLIMRSSTGRFYEFTTP
Manes.05G153000.1.p  MARGKIQIKRIENSTNRQVTYSKRRNGLFKKANELTVLCDARVSIIMFSTSGKLHEYISP
KAF2312359.1         MGRGKIEIKRIENPTNRQVTYSKRRNGIFKKAQELTVLCDAKVSLIMFSNTGKFHEFISP
Manes.01G141900.1.p  MGRGKIEIKRIENSTNRQVTYSKRRNGIFKKAQELTVLCDAKVSLIMFSNTGKFHEFISP
Potri.007G017000.1.p MGRGKIEIKKIENPTNRQVTYSKRRNGIFKKAQELTVLCDAKVSLIMFSNTNKLNEYISP
Glyma.11G073700.5.p  MGRGKIEIKLIENPTNRQVTYSKRRNGIFKKAHELSVLCDAKVSLIMFSKNNKMHEYISP
Rc05T012679.2        MGRGKIEIKKIENPTNRQVTYSKRRNGIFKKAQELTVLCDAKVSLIMFSNSGKFHEFISP
Potri.002G028400.1.p MARGKIQIKKIENSTNRQVTYSKRRNGLFKKAHELTVLCDAEVSLVMVSCTDKVHDYTSP

                               70        80        90        100       110       120
Manes.02G100400.1.p  TTTTKKVFDQYQETLGIDLWKTHYERMQEHLRKLREINNKLRRDIRQRMGEDLDDLSMDE
Glyma.01G169600.1.p  GLTTKKIIDQYQKTLGIDLWHSHYEKMLENLKKLKDINNKLRRQIRHRIGEDMDDMSFQQ
KAF2290821.1         TTTTKNVFDQYQKALGIDLWKTHYERMQEHLRKLREKNNKLRREIRQRMGEDLNDLSMDE
Potri.005G118000.2.p STTTKKIYDQYQKALGIDLWSAQYEKMQEQLRKLKDINHKLKKEIRQRIGEDLNELSIDH
AT3G54340.1          NTTTKEIVDLYQTISDVDVWATQYERMQETKRKLLETNRNLRTQIKQRLGECLDELDIQE
Glyma.04G027200.1.p  STSTKQFFDQYQMTLGVDLWNSHYENMQENLKKLKEVNRNLRKEIRQRMGDCLNELGMED
Glyma.06G027200.1.p  STSTKQFFDQYQMTLGVDLWNSHYENMQENLKKLKDVNRNLRKEIRQRMGDCLNDLGMED
KAF2310046.1         STSTKQMFDQYQKTLGVDIWSPQYEKMNETLRKLKEVNGNLRREIRQRMGECLNDLSFEN
Rc02T004810.1        STTTKQLIDQYQKTLGVDIWISQYEKMQDHLKKLKDVNRNLRTEIGQRMGECLSDLSFED
Casp07377.t1         TTTHKKIFDRYQKDSGVDLWQPQYQKMQEELRNLRDTNNKLGRAIRQRMGKDLHDMRFDE
Casp07378.t1         NTTHKEIFHRYQKASGVDLWQTQYQKMQEELRNLRDTNNKLGRAIRQRMGKDLHDMRFDE
Ep_chr8_g24861.t1    ETTTKKIFDIYQKEKKPDLWHIQYERMQEELKRLRDKNNMLKREIRHRVGKDLHDLSLGE
Ep_chr8_g24862.t1    ETTTKKIFDIYQKEKKSDLWHIQYERMQEELKRLRDKNNMLKGEIRHRVGKDLHDLSLGE
Casp10977.t1         TTTAKKMFDRYQKERDTDLWHNQYERMQEELKRPREKNHMLKRGIW.........LNFPQ
Ep_chr8_g24863.t1    STTHKKIIDRYQKASGIDLWDIKYQKMQEELRKLNDTNYKLTRAIRQRMGEDLNGMRCDE
Casp07379.t1         TTTAKKMIDRYQKTKKIDLWHTLYQKMKEDLEKLTERNDMLKREIGQRRGKDLDGLSFAQ
Manes.05G153000.1.p  STTTKQVFDQYQKTLGVDIWSSQYEKMQETLKKLKEANMNLRREIRKRMGECLNDMSFED
KAF2312359.1         TTTTKTMFDQYQNTLGIDLWSTHYERMQEHLRKLREINNKLRRDIRQRMGEDLNDVSFDE
Manes.01G141900.1.p  TITTKKMFDQYQKALGIDLWSTHYERMQEHLGKLREINNKLRRDIRQRMGEDLNDVSILE
Potri.007G017000.1.p STSTKKIYDQYQNALGIDLWGTQYEKMQEHLRKLNDINHKLRQEIRQRRGEGLNDLSIDH
Glyma.11G073700.5.p  GLTTKRIIDQYQKTLGIDLWRSHYEKMLENLKKLIDINNKLRRQIRHRIGEDMDDMSFQQ
Rc05T012679.2        GTTTKKIFDQYQKTLGIDLWITHYERMQDHLKKLNDINNKLRRDIRQRMGEDLNDLSFDE
Potri.002G028400.1.p STTTKRIFDQYQQTKGIDLWSSHYEIMKENLEKLKEVNMKIRREMRQRMGQCLNGLSFQD

                              130       140       150       160       170       180
Manes.02G100400.1.p  LFVLEQRMDSALELIRDRKYHVIKTQTETCKKKVKNLEERHGDLLLEYEAKYEDPQYGLV
Glyma.01G169600.1.p  LRTLEEDMVSSIGKIRERKFHVIKTRTDTCRKKVKSLKQMNGNLLLELKEKCVHPQFLLH
KAF2290821.1         LHGLEQRMASALEVVRERKYHVIKTQTDTHKKKVKNLEERHGDLVLEYEAKCEDPHCGLV
Potri.005G118000.2.p LRVLEQNMTEALNGVRGRKYHVIKTQTETYKKKVRSLEERHGNLWMEYEAKMEDPRYGLV
AT3G54340.1          LRRLEDEMENTFKLVRERKFKSLGNQIETTKKKNKSQQDIQKNLIHELELRAEDPHYGLV
Glyma.04G027200.1.p  LKLLEEEMDKAAKVVRERKYKVITNQIDTQRKKFNNEKEVHNRLLHDLDAKAEDPRFALI
Glyma.06G027200.1.p  LKLLEEEMDKAAKVVRERKYKVITNQIDTQRKKFNNEKEVHNRLLRDLDARAEDPRFALI
KAF2310046.1         LRGLEQDMDTALNLIRERKNRVIANQIVTWNKKLRNAEERYRNLLHELEARHEDPHYGLV
Rc02T004810.1        LRGLEQQMDNAIKVIRERKNKVVSNQIEKFNRKLRNLEKIQKNLLDEFEARVEDPHYGLV
Casp07377.t1         IFELEQTMISALDHVSKEKVRAIKRKKNTTGKKVRGMENEQRQMINMIEARYEE......
Casp07378.t1         IFELEQTMISALDHLNKEKVKIIKRMENTLDKKVSGMENEQREMI.....NME.......
Ep_chr8_g24861.t1    LIQLEETMISSQKVVHEAKLKDMHKKETTSKKKVRSLGQEQSTLAYHIGAKYEE..YGLL
Ep_chr8_g24862.t1    LIQLEETMISSQKVVHEAKVKDMHKKETTSKKKVNGMLKHQPQMIYP.DAKYEE..YGLL
Casp10977.t1         V.........................................SLRIPY...LX.......
Ep_chr8_g24863.t1    LFELEQTMTSALDHVSKERVKAIKRRKNTLSKKVNGMLKHQPQMIHP.DAKYEE..YGLL
Casp07379.t1         LFQLEQEMNSTLDAVRLLQVRDVKVKTYTLHKQVTSLDGTHERLKHEI.AILE.......
Manes.05G153000.1.p  LRGLEQDMDTALNVIRERKNRVIANQIVTSTKKLRNVEEVHRNLLHEYEARAEDPHYGLV
KAF2312359.1         LRGLEERMDSALELVRER..............KVRNLEERHGDLLLEYEAKCEDLQYGLV
Manes.01G141900.1.p  LRDLEERMASALEFVRDRKYHVIKTQTETCRKKVRNLEERHGDLLLEFEAKCEDLQYGLV
Potri.007G017000.1.p LRGLEQHMTEALNGVRGRKYHVIKTQNETYRKKVKNLEERHGNLLMEYEAKLEDRQYGLV
Glyma.11G073700.5.p  LRTLEEDLVSSIGKIRERKFHVIKTRTDTCRKKVKSLEQMNRDLLFELKEKCAHPQFILH
Rc05T012679.2        LRGLEQRMAPALHMIRERKYHFIETQTNTKRKKERNLVERHRELLQRYEANCEDPQYGLV
Potri.002G028400.1.p LQSLESDMESAWRVIHDRADRVLTNQIETSKKKARNVEQINRKLQVELEAMDQDP.YGLV
```

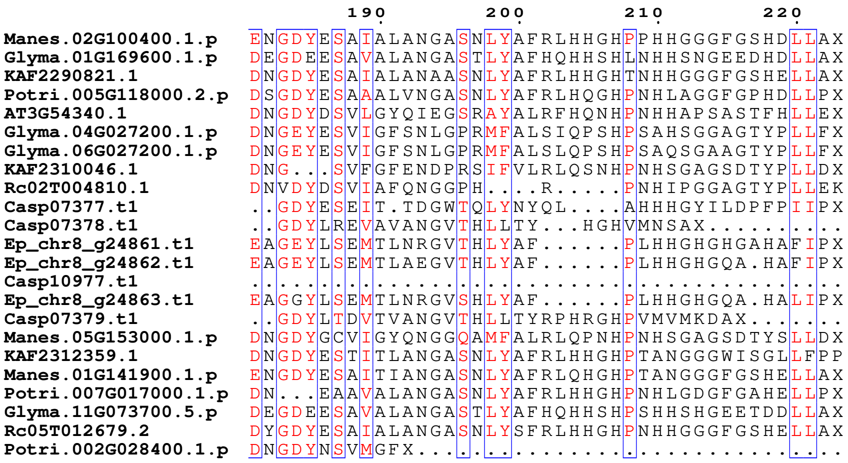
190 200 210 220
Manes.02G100400.1.p ENGDYESAIALANGASNLYAFRLHHGHPPHHGGGFGSHDLLAX
Glyma.01G169600.1.p DEGDEESAVALANGASTLYAFHQHHSHLNHHSNGEEDHDLLAX
KAF2290821.1 DNGDYESAIALANAASNLYAFRLHHGHTNHHGGGFGSHELLAX
Potri.005G118000.2.p DSGDYESAAALVNGASNLYAFRLHQGHPNHLAGGFGPHDLLPX
AT3G54340.1 DNGDYDSVLGYQIEGSRAYALRFHQNHPNHHAPSASTFHLLEX
Glyma.04G027200.1.p DNGEYESVIGFSNLGPRMFALSIQPSHPSAHSGGAGTYPLLFX
Glyma.06G027200.1.p DNGEYESVIGFSNLGPRMFALSLQPSHPSAQSGAAGTYPLLFX
KAF2310046.1 DNG...SVFGFENDPRSIFVLRLQSNHPNHSGAGSDTYPLLDX
Rc02T004810.1 DNVDYDSVIAFQNGGPH....R.....PNHIPGGAGTYPLLEK
Casp07377.t1 ..GDYESEIT.TDGWTQLYNYQL....AHHHGYILDPFPIIPX
Casp07378.t1 ..GDYLREVAVANGVTHLLTY...HGHVMNSAX..........
Ep_chr8_g24861.t1 EAGEYLSEMTLNRGVTHLYAF......PLHHGHGHGAHAFIPX
Ep_chr8_g24862.t1 EAGEYLSEMTLAEGVTHLYAF......PLHHGHGQA.HAFIPX
Casp10977.t1 ...........................................
Ep_chr8_g24863.t1 EAGGYLSEMTLNRGVSHLYAF......PLHHGHGQA.HALIPX
Casp07379.t1 ..GDYLTDVTVANGVTHLLTYRPHRGHPVMVMKDAX.......
Manes.05G153000.1.p DNGDYGCVIGYQNGGQAMFALRLQPNHPNHSGAGSDTYSLLDX
KAF2312359.1 DNGDYESTITLANGASNLYAFRLHHGHPTANGGGWISGLLFPP
Manes.01G141900.1.p ENGDYESAITIANGASNLYAFRLQHGHPTANGGGFGSHELLAX
Potri.007G017000.1.p DN...EAAVALANGASNLYAFRLHHGHPNHLGDGFGAHELLPX
Glyma.11G073700.5.p DEGDEESAVALANGASTLYAFHQHHSHPSHHSHGEETDDLLAX
Rc05T012679.2 DYGDYESAIALANGASNLYSFRLHHGHPNHHGGGFGSHELLAX
Potri.002G028400.1.p DNGDYNSVMGFX...............................

### LFY

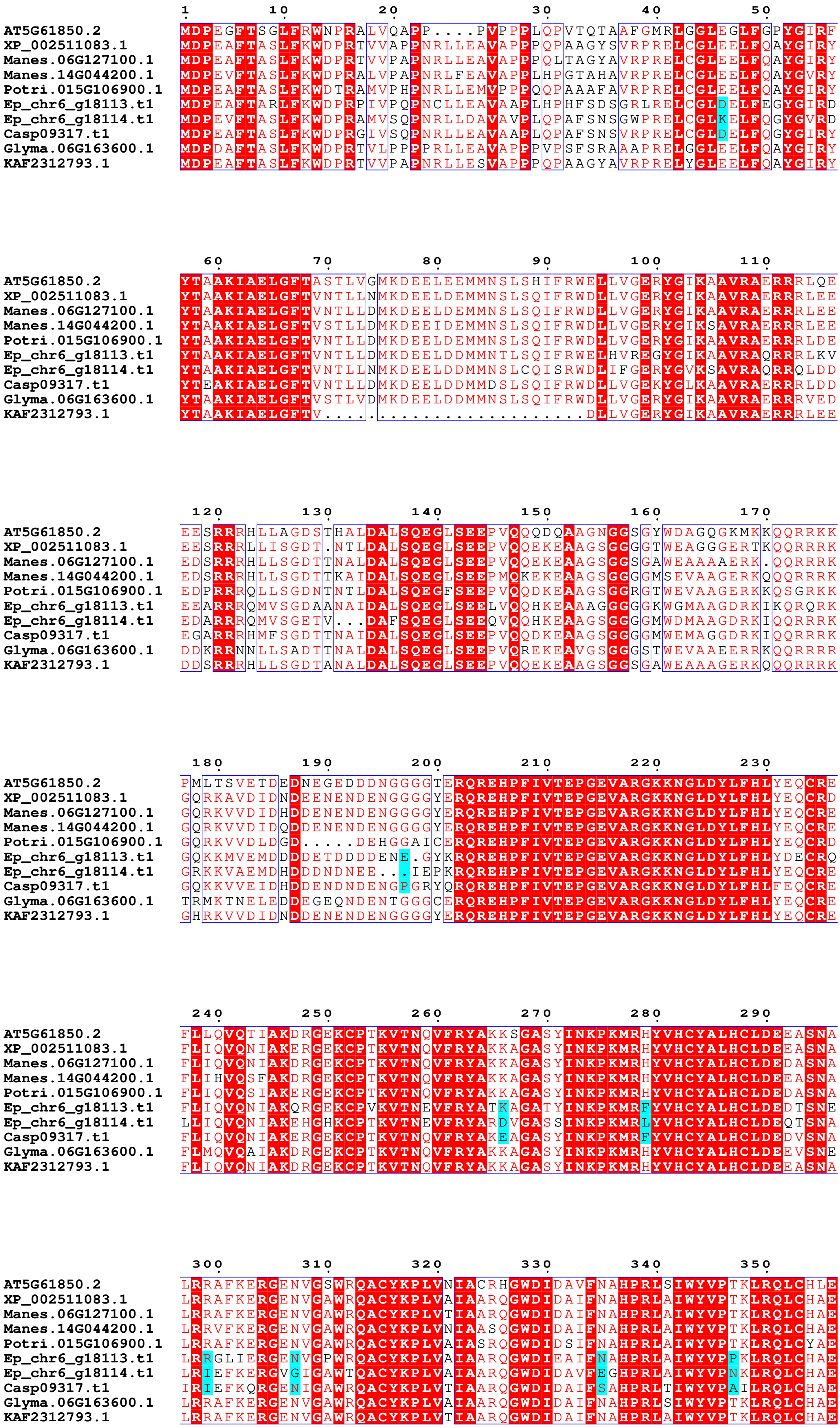
1 10 20 30 40 50
AT5G61850.2 MDPEGFTSGLFRWNPRALVQAPP....PVPPPLQPVTQTAAFGMRLGGLEGLFGPYGIRF
XP_002511083.1 MDPEAFTASLFKWDPRTVVAPPNRLLEAVAPPPQPAAGYSVRPRELCGLEELFQAYGIRY
Manes.06G127100.1 MDPEAFTASLFKWDPRTVVPAPNRLLEAVAPPPQLTAGYAVRPRELCGLEELFQAYGIRY
Manes.14G044200.1 MDPEVFTASLFKWDPRALVPAPNRLFEAVAPPLHPGTAHAVRPRELCGLEELFQAYGVRY
Potri.015G106900.1 MDPEAFTASLFKWDTRAMVPHPNRLLEMVPPPQQPAAAFAVRPRELCGLEELFQAYGIRY
Ep_chr6_g18113.t1 MDPEAFTARLFKWDPRPIVPQPNCLLEAVAAPLHPHFSDSGRLRELCGLDELFEGYGIRD
Ep_chr6_g18114.t1 MDPEVFTASLFKWDPRAMVSQPNRLLDAVAVPLQPAFSNSGWPRELCGLKELFQGYGVRD
Casp09317.t1 MDPEAFTASLFKWDPRGIVSQPNRLLEAVAAPLQPAFSNSVRPRELCGLDELFQGYGIRY
Glyma.06G163600.1 MDPDAFTASLFKWDPRTVLPPPPRLLEAVAPPPVPSFSRAAAPRELGGLEELFQAYGIRY
KAF2312793.1 MDPEAFTASLFKWDPRTVVPAPNRLLESVAPPPQPAAGYAVRPRELYGLEELFQAYGIRY
60 70 80 90 100 110
AT5G61850.2 YTAAKIAELGFTASTLVGMKDEELEEMMNSLSHIFRWELLVGERYGIKAAVRAERRRLQE
XP_002511083.1 YTAAKIAELGFTVNTLLNMKDEELDEMMNSLSQIFRWDLLVGERYGIKAAVRAERRRLEE
Manes.06G127100.1 YTAAKIAELGFTVNTLLDMKDEELDEMMNSLSQIFRWDLLVGERYGIKAAVRAERRRLEE
Manes.14G044200.1 YTAAKIAELGFTVSTLLDMKDEEIDEMMNSLSQIFRWDLLVGERYGIKSAVRAERRRLEE
Potri.015G106900.1 YTAAKIAELGFTVNTLLDMKDEELDEMMNSLSQIFRWDLLVGERYGIKAAVRAERRRLDE
Ep_chr6_g18113.t1 YTAAKIAELGFTVNTLLDMKDEELDDMMNTLSQIFRWELHVREGYGIKAAVRAQRRRLKV
Ep_chr6_g18114.t1 YTAAKIAELGFTVNTLLNMKDEELDDMMNSLCQISRWDLIFGERYGVKSAVRAQRRQLDD
Casp09317.t1 YTEAKIAELGFTVNTLLDMKDEELDDMMDSLSQIFRWDLLVGEKYGLKAAVRAERRRLDD
Glyma.06G163600.1 YTAAKIAELGFTVSTLVDMKDEELDDMMNSLSQIFRWDLLVGERYGIKAAVRAERRRVED
KAF2312793.1 YTAAKIAELGFTV.........................DLLVGERYGIKAAVRAERRRLEE
120 130 140 150 160 170
AT5G61850.2 EESRRRHLLAGDSTHALDALSQEGLSEEPVQQQDQAAGNGGSGYWDAGQGKMKKQQRRKK
XP_002511083.1 EESRRRLLISGDT.NTLDALSQEGLSEEPVQQEKEAAGSGGGGTWEAGGGERTKQQRRRK
Manes.06G127100.1 EDSRRRHLLSGDTTNALDALSQEGLSEEPVQQEKEAAGSGGSGAWEAAAAERK.QQRRRK
Manes.14G044200.1 EDSRRRHLLSGDTTKAIDALSQEGLSEEPMQKEKEAAGSGGGGMSEVAAGERKQQQRRRK
Potri.015G106900.1 EDPRRRQLLSGDNTNTLDALSQEGFSEEPVQQDKEAAGSGGRGTWEVAAGERKKQSGRKK
Ep_chr6_g18113.t1 EEARRRQMVSGDAANAIDALSQEGLSEELVQQHKEAAAGGGGGKWGMAAGDRKIKQRQRK
Ep_chr6_g18114.t1 EDARRRQMVSGETV...DAFSQEGLSEEQVQQHKEAAGSGGGGMWDMAAGDRKIQQRRRK
Casp09317.t1 EGARRRHMFSGDTTNAIDALSQEGLSEEPVQQDKEAAGSGGGGMWEMAGGDRKIQQRRRK
Glyma.06G163600.1 DDKRRNNLLSADTTNALDALSQEGLSEEPVQREKEAVGSGGGSTWEVAAEERRKQQRRRR
KAF2312793.1 DDSRRRHLLSGDTANALDALSQEGLSEEPVQQEKEAAGSGGSGAWEAAAGERKQQQRRRK
180 190 200 210 220 230
AT5G61850.2 PMLTSVETDEDNEGEDDDNGGGTERQREHPFIVTEPGEVARGKKNGLDYLFHLYEQCRE
XP_002511083.1 GQRKAVDIDNDEENENDENGGGGYERQREHPFIVTEPGEVARGKKNGLDYLFHLYEQCRD
Manes.06G127100.1 GQRKVVDIDHDDENENDENGGGGYERQREHPFIVTEPGEVARGKKNGLDYLFHLYEQCRE
Manes.14G044200.1 GQRKVVDIDQDDENENDENGGGGYERQREHPFIVTEPGEVARGKKNGLDYLFHLYEQCRE
Potri.015G106900.1 GQRKVVDLDGD.....DEHGGAICERQREHPFIVTEPGEVARGKKNGLDYLFHLYEQCRD
Ep_chr6_g18113.t1 GQKKMVEMDDDDETDDDDENE.GYKRQREHPFIVTEPGEVARGKKNGLDYLFHLYDECRQ
Ep_chr6_g18114.t1 GRKKVAEMDHDDDNDNEE...IEPKRQREHPFIVTEPGEVARGKKNGLDYLFHLYEQCRE
Casp09317.t1 GQKKVVEIDHDDENDNDENGPGRYQRQREHPFIVTEPGEVARGKKNGLDYLFHLFEQCRE
Glyma.06G163600.1 TRMKTNELEDDEGEQNDENTGGGCERQREHPFIVTEPGEVARGKKNGLDYLFHLYEQCRE
KAF2312793.1 GHRKVVDIDNDDENENDENGGGGYERQREHPFIVTEPGEVARGKKNGLDYLFHLYEQCRE
240 250 260 270 280 290
AT5G61850.2 FLLQVQTIAKDRGEKCPTKVTNQVFRYAKKSGASYINKPKMRHYVHCYALHCLDEEASNA
XP_002511083.1 FLIQVQNIAKERGEKCPTKVTNQVFRYAKKAGASYINKPKMRHYVHCYALHCLDEEASNA
Manes.06G127100.1 FLIQVQNIAKDRGEKCPTKVTNQVFRYAKKAGASYINKPKMRHYVHCYALHCLDEDASNA
Manes.14G044200.1 FLIHVQSFAKDRGEKCPTKVTNQVFRYAKKAGASYINKPKMRHYVHCYALHCLDEEASNA
Potri.015G106900.1 FLIQVQSIAKERGEKCPTKVTNQVFRYAKKAGASYINKPKMRHYVHCYALHCLDEDASNA
Ep_chr6_g18113.t1 FLIQVQNIAKQRGEKCPVKVTNEVFRYATKAGATYINKPKMRFYVHCYALHCLDEDTSNE
Ep_chr6_g18114.t1 LLIQVQNIAKEHGHKCPTKVTNEVFRYARDVGASSINKPKMRLYVHCYALHCLDEQTSNA
Casp09317.t1 FLIQVQNIAKERGEKCPTKVTNQVFRYAKEAGASYINKPKMRFYVHCYALHCLDEDVSNA
Glyma.06G163600.1 FLMQVQAIAKDRGEKCPTKVTNQVFRYAKKAGASYINKPKMRHYVHCYALHCLDEEVSNE
KAF2312793.1 FLIQVQNIAKDRGEKCPTKVTNQVFRYAKKAGASYINKPKMRHYVHCYALHCLDEEASNA
300 310 320 330 340 350
AT5G61850.2 LRRAFKERGENVGSWRQACYKPLVNIACRHGWDIDAVFNAHPRLSIWYVPTKLRQLCHLE
XP_002511083.1 LRRAFKERGENVGAWRQACYKPLVAIAARQGWDIDAIFNAHPRLAIWYVPTKLRQLCHAE
Manes.06G127100.1 LRRAFKERGENVGAWRQACYKPLVTIAARQGWDIDAIFNAHPRLAIWYVPTKLRQLCHAE
Manes.14G044200.1 LRRVFKERGENVGAWRQACYKPLVNIAASQGWDIDAIFNAHPRLAIWYVPTKLRQLCHAE
Potri.015G106900.1 LRRAFKERGENVGAWRQACYKPLVAIASRQGWDIDSIFNAHPRLAIWYVPTKLRQLCYAE
Ep_chr6_g18113.t1 LRRGLIERGENVGPWRQACYKPLVAIAARQGWDIEAIFNAHPRLAIWYVPPKLRQLCHAE
Ep_chr6_g18114.t1 LRIEFKERGVGIGAWTQACYKPLVAIAARQGWDIDAVFEGHPRLAIWYVPNKLRQLCHAE
Casp09317.t1 IRIEFKQRGENIGAWRQACYKPLVTIAARQGWDIDAIFSAHPRLTIWYVPAILRQLCHAE
Glyma.06G163600.1 LRRAFKERGENVGAWRQACYKPLVAIAARQGWDIDAIFNAHPRLSIWYVPTKLRQLCHAE
KAF2312793.1 LRRAFKERGENVGAWRQACYKPLVTIAARQGWDIDAIFNAHPRLAIWYVPTKLRQLCHAE

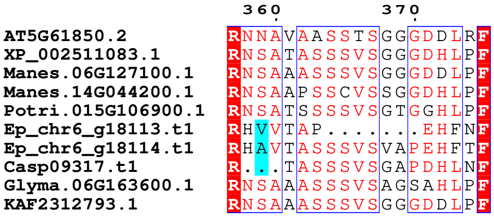
360 370
AT5G61850.2 RNNAVAASSTSGGGDDLRF
XP_002511083.1 RNSATASSSVSGGGDHLPF
Manes.06G127100.1 RNSAAASSSVSGGGDDLPF
Manes.14G044200.1 RNSAAPSSCVSSGGDHLPF
Potri.015G106900.1 RNSATSSSSVSGTGGHLPF
Ep_chr6_g18113.t1 RHVVTAP........EHFNF
Ep_chr6_g18114.t1 RHAVTASSSVSVAPEHFTF
Casp09317.t1 R...TASSSVSGAPDHLNF
Glyma.06G163600.1 RNSAAASSSVSAGSAHLPF
KAF2312793.1 RNSAAASSSVSGGGDDLPF

### PHYA

```
                          1        10        20        30        40        50        60
AT1G09570.3           MSGSRPTQSSEGSRRSRHSARIIAQTTVDAKLHADFEESGSSFDYSTSVRVTGPVENQPP
Ep_chr4_g08890.t1     MSSSRPSQSATSSSRSKHSARIIAQTTVDAKLHAEFETSGSSFDYSNSVRVTSSFGVQPP
KAF2324094.1          MSSSRPSHSSSNSVRSRHSTRIIAQTTVDAKLHADFEESGSSFDYSNLVRITSSVGDQPP
Potri.013G000300.2.p  MSSSRPSHSSSNSARSRHSARIIAQTTVDAKLHADFEESGSSFDYSSSVRVTDSVGDQPP
Glyma.19G224200.1.p   MSSSRPSQSSSNSGRSRTSARRMAQTTLDAKLHATFEESGSSFDYSSSVRMSGTVGDHQP
Glyma.20G090000.3.p   MSTSRPSQSSSNSGRSRRSARAMALATVDAKLHATFEESGSSFDYSSSVRISGTAGVNQP
Glyma.10G141400.1.p   MSTSRPSQSSSNSRRSRHSAR.MAQATVDAKIHATFEESGSSFDYSSSVRVSGTAGVNQP
KAF2307360.1          ............................................................
Manes.09G182500.1.p   MSSSRPSHSSSSSVRSRHSARIIAQTTVDAKLHAEFEESGSSFDYSNSVRVTNSVGDQLP
Casp30444.t1          MSSSRPAHSSSSSTRSKHSARIIAQTTVDAKLHAEFESSGSSFDYSNSVRVTSSFGIQPP
XP_002512596.1        MSSSRPSHSSSNSVRSRHSARIISQTAVDAKLHADFEESGSSFDYSNSVHVTSSTLDHAP

                                70        80        90       100       110       120
AT1G09570.3           RSDKVTTTYLHHIQKGKLIQPFGCLLALDEKTFKVIAYSENASELLTMASHAVPSVGEHP
Ep_chr4_g08890.t1     RSDKVTTAYLHHIQKGKTIQPFGCLLALDEKTLKVIAYSENAPEMLTMVSHAVPSVGENP
KAF2324094.1          RSDKVTKAYLHQIQKGKLIQPFGCLLALDEKTLKVIAYSENAPEMLTTVSHAVPSVGDHP
Potri.013G000300.2.p  RSDKVTTTYLHHIQKGKLIQPFGCLLALDEKTFKVVAYSENAPELLTMVSHAVPSVGEHP
Glyma.19G224200.1.p   RSDRATSSYLHQTQKIKLIQPFGCLLALDEKTCKVIAYSENAPEMLTMVSHAVPSVGDHP
Glyma.20G090000.3.p   RHDKVTTAYLHHMQKGKMIQPFGCLLALDEKTCKVIAYSENAPEMLTMVSHAVPSVGDHP
Glyma.10G141400.1.p   RSDKVTTAYLNHMQRGKMIQPFGCLLAIDEKTCKVIAYSENAPEMLTMVSHAVPSVGDHP
KAF2307360.1          ............................................................
Manes.09G182500.1.p   RSDKVTSAYLHQIQKGKLIQPFGCLLALDEKTFKVIAYSENAPEMLTTVSHAVPSVGDQL
Casp30444.t1          RSDKVTTAYLHHIQKGKMIQSFGCLLALDDKTLKVIAYSENAPEMLTMVSHAVPSVGENP
XP_002512596.1        RSDKVTTAYLHHIQKGKLIQPFGCLLALDEKTYKVIAYSENAPEMLTMVSHAVPSVGDHP

                               130       140       150       160       170       180
AT1G09570.3           VLGIGTDIRSLFTAPSASALQKALGFGDVSLLNPILVHCRTSAKPFYAIIHRVTGSIIID
Ep_chr4_g08890.t1     VLGFNSDVKTIFTAPSAIALQKALGFEDVSLLNPVLVHCKSSGRPFYAIIHRVTGSFIID
KAF2324094.1          VLGIGTDIKTIFTAPSASALQKALGFGDVSLLNPILVHCKTSGKPFYAIVHRVTGSLIID
Potri.013G000300.2.p  VLGIGTDIRTIFTAPSASALQKAMGFGDVSLLNPILVHCKTSGKPFYAIVHRVTGSLIID
Glyma.19G224200.1.p   ALGIGTDIRTIFTAPSSAAIQKALRFGDVSLHNPILVHCKTSGKPFYAIIHRVTGSVIID
Glyma.20G090000.3.p   ALGIGTDIKTLFTAPSASALQKALGFAEVLLLNPVLIHCKTSGKPFYAIIHRVTGSMIID
Glyma.10G141400.1.p   ALGIGTDIKTLFTAPSVSGLQKALGCADVSLLNPILVHCKTSGKPFYAIVHRVTGSLIVD
KAF2307360.1          ............................................................
Manes.09G182500.1.p   VLGIGTDIRTIFTAPSASALQKALGFGDVSLLNPILVHCKTSGKPFYAIVHRVTGSFIVD
Casp30444.t1          VLGFNSDVKNIFTAPSATALLKALGFGDVSLLNPVLVHCKTSGRPFYAIIHRVTGGFIID
XP_002512596.1        VLGIGTDIRTIFTAPSASALQKALGFGDVSLLNPILVHCKTSGKPFYAIVHRVTGSFIID

                               190       200       210       220       230       240
AT1G09570.3           FEPVKPYEVPMTAAGALQSYKLAAKAITRLQSLPSGSMERLCDTMVQEVFELTGYDRVMA
Ep_chr4_g08890.t1     FEPVKPYEVPMTASGALQSYKLAAKAISRLQSLPSGSMGRLCDTMVQEVFELTGYDRVMT
KAF2324094.1          FEPVKPYEVPMTAAGALQSYKLAAKAITRLQSLPSGSMERLCDTMVQEVSELTGYDRVMT
Potri.013G000300.2.p  FEPVKPYEVPMTAAGALQSYKLAAKAITRLQSLPSGSMERLCDTMVQEVFELTGYDRAMA
Glyma.19G224200.1.p   FEPVKPHEVPMTASGALQSYKLAAKAITRLESLTTGNMETLCNTMVREVFELTGYDRVMA
Glyma.20G090000.3.p   FEPVKPYEVPMTAAGALQSYKLAAKAITRLQSLPSGSMERLCDTMVQEVFELTGYDRVMA
Glyma.10G141400.1.p   FEPVKPYEVPMTAAGALQSYKLAAKAITRLQSLPSGNMERLCDTMVQEVFELTGYDRVMA
KAF2307360.1          ............................................................
Manes.09G182500.1.p   FEPVKPYEVPMTAAGALQSYKLAAKAITRLQSLPSGSMERLCDTMVQEVFELTGYDRVMT
Casp30444.t1          FEPVKPYEVPMTASGALQSYKLAAKAIARLQSLPSGSIERLCDTMVQEVFELTGYDRVMT
XP_002512596.1        FEPVKPYEVPMTAAGALQSYKLAAKAISRLQSLPSGSMERLCDTMVQEVFELTGYDRVMT

                               250       260       270       280       290       300
AT1G09570.3           YKFHEDDHGEVVSEVTKPGLEPYLGLHYPATDIPQAARFLFMKNKVRMIVDCNAKHARVL
Ep_chr4_g08890.t1     YKFHDDDHGEVISEITKPGLEPYLGLHYPATDIPQASRFLFMKNKVRMIVDCNARNVKVL
KAF2324094.1          YKFHDDDHGEVIAESTKPGLEPYLGLHYPATDIPQSARFLFMKNKVRMIVDCRAKHVKVL
Potri.013G000300.2.p  YKFHDDDHGEVVSEVTKPGLEPYLGLHYPATDIPQASRFLFMKNKVRMIVDCHAKHVKVL
Glyma.19G224200.1.p   YKFHEDDHGEVIAEVKRPGLEPYLGLHYPATDIPQATRFLFMKNKVRMIVDCCAKHVNVL
Glyma.20G090000.3.p   YKFHEDDHGEVIAEITKPGLEPYLGLHYPATDIPQASRFLFMKNKVRMIVDCHAKHVRVL
Glyma.10G141400.1.p   YKFHEDDHGEVIAEITKPGLEPYLGLHYPATDIPQASRFLFRKNKVRMIVDCHAKHVRVL
KAF2307360.1          ............................................................
Manes.09G182500.1.p   YKFHDDDHGEVISEITKPGLEPYLGLHYPATDIPQSARFLFMKNKVRMIVDCRAKQVKVL
Casp30444.t1          YKFHDDDHGEVISEITKPGLEPYLGLHYPATDIPQASRFLFMKNKVRMIVDCNAKNVKVL
XP_002512596.1        YKFHDDDHGEVISEVTKPGLEPYLGLHYPATDIPQAARFLFMKNKVRMIVDCRAKHVKVL

                               310       320       330       340       350       360
AT1G09570.3           QDEKLSFDLTLCGSTLRAPHSCHLQYMANMDSIASLVMAVVVNEDGEGDADATQPQKRK
Ep_chr4_g08890.t1     QDEKLPFELTLCGSTLRAPHSCHLQYMDNMTSIASLVMAVIINEGDDDDDASGQPQKRK
KAF2324094.1          QDEKLSFDLTLCGSTLRAPHSCHLQYMENMDSIASLVMAVVINEGDEDDSTSVQSQK.E
Potri.013G000300.2.p  QDEKLPFDLTLCGSTLRAPHSCHLQYMENMNSIASLVMAVVVNDGDEDGTDSVNPQKRK
Glyma.19G224200.1.p   QDKKIPFDLTLCGSTLRAAHSCHLQYMENMNSSASLVMAVVVNDNDEDGSDAVQPQKSK
Glyma.20G090000.3.p   QDEKLPFDLTLCGSTLRAPHSCHAQYMANMDSIASLVMAVVVNDNEEDGTDAIQPQKRK
Glyma.10G141400.1.p   QDEKLQFDLILCGSTLRAPHSCHAQYMANMDSIASLVLAVVVNDNEEDGTDAVQPQKRE
KAF2307360.1          ...........................................................
Manes.09G182500.1.p   QDEKLPFDLTLCGSTLRAPHSCHLQYMENMDSIASLVMAVVINEGDEDDSTSVQTQKRK
Casp30444.t1          QDEKLPMDLTLCGSTLRAPHSCHLQYMDNMTSIASLVMAVIINEGDDDDDASGQPQKRK
XP_002512596.1        QDEKLPLELTLCGSTLRAPHSCHLQYMENMDSVASLVMAVVVNEGDEDDSTSVQPQKRK
```

```
                       370       380       390       400       410       420
AT1G09570.3            RLWGLVVCHNTTPRFVPFPLRYACEFLAQVFAIHVNKEVELDNQMVEKNILRTQTLLCDM
Ep_chr4_g08890.t1      RLWGLVVCHNTTPRFVPFPLRYACEFLAQVFAIHVNKELELENQIVEKNILRTQTLLCDM
KAF2324094.1           RDFG..............................VYKELELENQIVEKT.LANQTLLCDM
Potri.013G000300.2.p   RLWGLVVCHNTSPRFVPFPLRYACEFLAQVFAIHVNKELELENQIVEKNILRTQTLLCDM
Glyma.19G224200.1.p    RLWGLVVCHHTTPRFVPFPLRYACQFLAQVFAVHVSKELEIEYQIIEKNILQTQTLLCDM
Glyma.20G090000.3.p    RLWGLVVCHNTTPRFVPFPLRYACEFLAQVFAIHVNKEIELEYQIIEKNILRTQTLLCDL
Glyma.10G141400.1.p    RLWGLVVCHNTTPRFVPFPLRYACEFLAQVFAVHVHKEIELEYQIIEKNILRTQALLC.M
KAF2307360.1           ...........................................................M
Manes.09G182500.1.p    RLWGLVVCHNTTPRFVPFPLRYACEFLVQVFAIHVNKELELENQIVEKNILRTQTLLCDM
Casp30444.t1           RLWGLVVCHNTSPRFVPFPLRYACEFLAQVFAIHVSKELELENQIVEKNILRTQTLLCDM
XP_002512596.1         RLWGLVVCHNTTPRFVPFPLRYACEFLAQVFAIHVNKELELENQIVEKNILRTQTLLCDM
```

```
                       430       440       450       460       470       480
AT1G09570.3            LMRDAPLGIVSQSPNIMDLVKCDGAALLYKDKIWKLGTTPSEFHLQEIASWLCEYHMDST
Ep_chr4_g08890.t1      LMRDVPLGIVSQHPNIMDLVKCDGAVLLYQNKLWRLGVTPDECQIRDIAVWLSDYHKDST
KAF2324094.1           LMRDAPLGIVTQSANIMDLVKCDGAALLYKNKIWRLGVTPSDRQIQDIAVWLSEYHMDST
Potri.013G000300.2.p   LMRDAPLGIVTQSPNIMDLVKCDGAVLFYRNKIWRLGITPSDLQLQDIAFWLSEYHMDST
Glyma.19G224200.1.p    LVQGEPLGIVSQSPNIMDLVKCDGAALLYKNKVWRLGVTPSESQIKEIALWLFECHEDST
Glyma.20G090000.3.p    VMRDAPLGIVSESPNIMDLVKCDGAALIYKNKVWRLGVTPSESQIREIAFWLSEYHMDST
Glyma.10G141400.1.p    LMRDAPLGIVSESPNIMDLVKCDGAALIYRNKVWRLGVTPSEPQIREIALWLSEYHMDST
KAF2307360.1           LMRDAPLGIVMQSPNIMDLVKCDGAALFYKNKLWRLGVTPSDLQIQDIAVWLSEYHMDST
Manes.09G182500.1.p    LMRDAPLGIIVQSPNIMDLVKCDGAALLYKNKIWRLGATPSDLQIEDIAVWLSEYHMDST
Casp30444.t1           LMRDAPLGIVSQHPNIMDLVKCDGAVLFYQNKLWRLGATPDERQIRDIAIWLSDYHKDST
XP_002512596.1         LLRDAPLGILTQSPNITDLVKCDGAALLYKNKIWRLGVTPSDLQIRDIAVWLSEYHMDST
```

```
                       490       500       510       520       530       540
AT1G09570.3            GLSTDSLHDAGFPRALSLGDSVCGMAAVRISSKDMIFWFRSHTAGEVRWGGAKHDPDDRD
Ep_chr4_g08890.t1      GLSTDSLHDAGFPGALSFGDVVCGMAAVRIASRDMLFWFRSPTAAEVRWGGAKHEPGERD
KAF2324094.1           GLSTDSLFDAGYPGALTLGDVVCGMAAVRITSKDMLFWFRSQTAAEIRWGGAKHEPGEKD
Potri.013G000300.2.p   GLSTDSLYDAGYPGALALGDVVCGMAAVRITSKDMLFWFRSQTAAEIRWGGAKHEPGEKD
Glyma.19G224200.1.p    GFCTDSLSDAGFPGAAALGDIACGMAAARIASKDILFWFRSHTASEIRWGGAKHEPGERD
Glyma.20G090000.3.p    GFSTDSLSDAGFPSALSLGDVVCGMAAVRVTAKDVVFWFRSHTAAEIRWGGAKHEAGEKD
Glyma.10G141400.1.p    GFSTDSLFDAGFPSALSLGDVVCGMASVRVTAKDMVFWFRSHTAAEIRWGGAKHEAGEKD
KAF2307360.1           GLSTDSLYDAGYPGALALGDVVCGMAAVRITAKDMLFWFRSQTAAEIRWGGAKHEPGEKD
Manes.09G182500.1.p    GLSTDSLYDAGYPAALALGDVVCGMAAVRITSKDMLFWFRCQTAAEIRWGGAKHEPGEKD
Casp30444.t1           GLSTDSLHDAGFPGALSFSDVVCGMAAVRISSKDMLFWFRSPASAEVRWGGAKHEPGERD
XP_002512596.1         GLSTDSLYDAGYSAALSLEDVVCGMAAVRITSKDMLFWFRAPTAAEIRWGGAKHEPGEKD
```

```
                       550       560       570       580       590       600
AT1G09570.3            DARRMHPRSSFKAFLEVVKTRSLPWKDYEMDAIHSLQLILRNAFKDSETTDVNTKVIYSK
Ep_chr4_g08890.t1      DNRRMHPRSSFKAFLEVVKARSAPWKDYEMDAIHSLQLILRNAFKDLEGGGADSR.IRTR
KAF2324094.1           D...........................................................
Potri.013G000300.2.p   DGRRMHPRSSFKAFLEVVKTRSLPWKDYEMDAIHSLQLILRNAFKDIETMDVDTKTIHAR
Glyma.19G224200.1.p    DGRRVHPRSSFKAFLEVVKTRSLPWKTYETDAIHSLQLILRDAFKETQSMEISTYAIDTR
Glyma.20G090000.3.p    DGRRMHPRSSFKVFLDVVKARSLPWKEYEIDAMHSLQLILRNAFKDTESMDLNTKAINTR
Glyma.10G141400.1.p    DSRRMHPRSSFKAFLEVVKARSLPWKEYEMDAIHSLQIILRNAFKDTESLDLNAKAINTR
KAF2307360.1           DGRRMHPRSSFKAFLEVAKTRSLPWKDYEMDAIHSLQLILRNAFKDVETMDTHTKTIHSR
Manes.09G182500.1.p    DGRRMHPRSSFKAFLEVSKTRSLPWKDYEMDAIHSLQLILRNAFKDIESVDMHTKTIHSR
Casp30444.t1           DNRKMQPRSSFKAFLEVVKARSSPWKDFEMDAIHSLQLILRNAFKDRESGGGDSVMIRSR
XP_002512596.1         DGRKMHPRSSFKAFLEVVKTRSLPWKDYEMDAIHSLQLILRNAFKDAETMDADAKAIHSR
```

```
                       610       620       630       640       650       660
AT1G09570.3            LNDLKIDGIQELEAVTSEMVRLIETATVPILAVDSDGLVNGWNTKIAELTGLSVDEAIGK
Ep_chr4_g08890.t1      LNELKIEGMQELEAVTSEMVRLIETATVPILAVDVDGLVNGWNTKIAELTGLPVDHAIGK
KAF2324094.1           ................................VDVDGLVNGWNTKIAELTGLPVDKAIGK
Potri.013G000300.2.p   LSDLKIEGMQELEAVTSEMVRLIETATVPILAVDVDGLVNGWNTKISELTGLLVDKAIGK
Glyma.19G224200.1.p    LGDLKIEGMQELDAVTSEVVRLIETATVPILAVDVNGMINGWNTKIAELTGLPVDEAIGK
Glyma.20G090000.3.p    LSDLKIEGMQELEAVTSEIVRLIETATVPILAVDVDGLVNGWNIKIAELTGLPVGEAMGK
Glyma.10G141400.1.p    LSDLKIEGMQELEAVTSEIVRLIDTATVPILAVDVDGLVNGWNIKIAELTGLPIGEATGK
KAF2307360.1           LSDLKIEGMQELEAVTSEMVRLIETATVPILAVDVDGLVNGWNTKIAELTGLPVDKAIGK
Manes.09G182500.1.p    LSDLKIEGMQELEAVTSEMVRLIETATVPILAVDVDGLVNGWNTKIADLTGLPVDKAIGK
Casp30444.t1           LNELKIEGMQELEAVTSEMVRLIETASVPILAVDVDGLVNGWNTKLSELTGLSVDNAIGK
XP_002512596.1         LSDLKIEGMQELEAVTSEMVRLIETATVPILAVDVDGLVNGWNTKIAELTGLPVDKAIGK
```

```
                       670       680       690       700       710       720
AT1G09570.3            HFLTLVEDSSVEIVKRMLENALEGTEEQNVQFEIKTHLSRADAGPISLVVNACASRDLHE
Ep_chr4_g08890.t1      HLLTLVEDSSLDIVKSMLFSALQGKEEQNIQFEIKTYSSKAESGPVSLVVNACASKDVQA
KAF2324094.1           HLLTLVEESSIDIVKKMLFLALQGKEEQNIQFEIKTHGSKVEGGPISLVVNACASRDIHE
Potri.013G000300.2.p   HLLTLVEDSSVDIVKRMLFLALQGKEEQNIQFEIKTHGSKSECGPICLVVNACASRDLHE
Glyma.19G224200.1.p    HLLTLVEDFSVDRVKKMLDMALQGEEERNVQFEIQTHHMKIDSGPISLVVNACASRDLQD
Glyma.20G090000.3.p    HLLTLVEDSSTDRVKKMLNLALLGEEEKNVQFEIKTHGSKMDSGPISLVVNACASRDLRD
Glyma.10G141400.1.p    HLLTLVEDSSTDRVKKMLNLALLGEEEKNVQFEIKTHGSKMDSGPISLVVNACASRDLRD
KAF2307360.1           HLLTLVEESSFDIVKKMLFLALQGKEEQNIQFEIKTHGSKADDGPISLVVNACSSRDIQE
Manes.09G182500.1.p    HLLTLVEESSFDIVKNMLFLALQGKEERNVQFDIKTHGSKAEDGPISLIANACSSRDINE
Casp30444.t1           HLLTLVEDSSLDIVKNMLFSALQGKEEQNIQFELKTYSSKAENCPVSLIVNACASKDVQG
XP_002512596.1         HLLTLVEDGSIDLVKNMLFSALQGKEEQNIQFEIKTHGSKVESGPISLVVNACASRDISE
```

```
                                730       740       750       760       770       780
AT1G09570.3           NVVGVCFVAHDLTGQKTVMDKFTRIEGDYKAIIQNPNPLIPPIFGTDEFGWCTEWNPAMS
Ep_chr4_g08890.t1     NVVGVCFVAQDITGEKTVMDKFTRIEGDYKAIVQNPNPLIPPIFGTDEFGWCSEWNPAME
KAF2324094.1          SVVGVCFVAQDVTGEKIVMDKFTRIEGDYKAIVQNPNPLIPPIFGTDEFGXCSEWNPAMX
Potri.013G000300.2.p  NVVGVCFVGQDITGQKMVMDKFTRIEGDYKAIVQNRNPLIPPIFGTDEFGWCSEWNPAMT
Glyma.19G224200.1.p   NVVGVCFLAQDITAQKTMMDKFTRIEGDYKAIVQNPNPLIPPIFGTDEFGWCCEWNSAMA
Glyma.20G090000.3.p   NVVGVCFVAHDITAQKNVMDKFTRIEGDYKAIVQNRNPLIPPIFGTDEFGWCCEWNPAMT
Glyma.10G141400.1.p   NVVGVCFVAHDITAQKNVMDKFIRIEGDYKAIVQNRNPLIPPIFGTDEFGWCCEWNPAMM
KAF2307360.1          NVVGVCFVAQDVTSQKTVMDKFTRIEGDYKAIVQNPNPLIPPIFGTDEFGWCSEWNPAMT
Manes.09G182500.1.p   NVVGVCFVAQDITSQKTVMDKFTRIEGDYKAIVQNPNPLIPPIFGTDEFGWCSEWNPAMT
Casp30444.t1          NVVGVCFVAQDITGQKTVMDKFTRIEGDYKAIVQNPNPLIPPIFGTDEFGWCSEWNPAMV
XP_002512596.1        NVVGVCFVAQDITGQKTVMDKFTRIEGDYKAIVQNPNPLIPPIFGTDEFGWCSEWNPAMA

                                790       800       810       820       830       840
AT1G09570.3           KLTGLKREEVIDKMLLGEVFGTQKSCCRLKNQEAFVNLGIVLNNAVTSQDPEKVSFAFFT
Ep_chr4_g08890.t1     KLTGWKREEVIDKMLLGEVFGINRACCRLRNRETFVNLGILLNNAMTGQETEKVAFSFLA
KAF2324094.1          ............................................................
Potri.013G000300.2.p  NLTGWKREEVLDKMLLGEVFGLNMACCRLKNQEAFVNLGVVLNTAMTGQESEKVSFGFFA
Glyma.19G224200.1.p   KLTGWKREEVMDKMLLGEVFGTQIACCRLRNHEAVVNFSIVLNTAMAGLETEKVPFGFFA
Glyma.20G090000.3.p   KLTGWKREEVMDKMLLGELFGTHMAACRLKNQEAFVNLGVVLNKAMTGLETEKVPFGFFA
Glyma.10G141400.1.p   KLTGWKREEVMDKMLLGEIFGTQMAACRLKNQEAFVNLGVVLNKAMTGSETEKVPFGFFA
KAF2307360.1          KLTGWKREEVIDKMLLGEVFGINMACCRLKNQEAFLNLGIILNNAMTGQEPEKVHFSFFA
Manes.09G182500.1.p   KLTGWKREEVIDKMLLGEVFGINMACCRLKNQEAFLNLGIVLNNAMTGAEPEKVSFSFSA
Casp30444.t1          KLTGWKREEVIDKMLLGEVFGINRACCRLRNRETFVNLGILLNNAMTGQETEKVTFSFLA
XP_002512596.1        KLTGWKREEVMDKMLLGEVFGINRACCCLKNQEAFVNLGVLINNAMTSQVPEKVSFSFFA

                                850       860       870       880       890       900
AT1G09570.3           RGGKYVECLLCVSKKLDREGVVTGVFCFLQLASHELQQALHVQRLAERTAVKRLKALAYI
Ep_chr4_g08890.t1     RNGKYVECLLCLSKKLNREGAVTGIFCFLQLASQELQQALHIQQLSEQIALKRLKTLAYM
KAF2324094.1          ............................................................
Potri.013G000300.2.p  RTGKYVECLLCVSKKLDREGAVTGVFCFLQLASQELQQALHVQRLSEQTALKRLKALAYL
Glyma.19G224200.1.p   RDGKHVECILSMTKKLDAEGVVTGVFCFLQLASAELQQALHIQRISEQTSLKRLKDLTYL
Glyma.20G090000.3.p   RNGKYVECLLSVSKKLDVEGLVTGVFCFLQLASPELQQALHIQRLSEQTALKRLNALSYM
Glyma.10G141400.1.p   RNGKYVECLLSVSKKLDVEGLVTGVFCFLQLASPELQQALHIQRLSEQTASKRLNALSYM
KAF2307360.1          RNGNWIACYVX.................................................
Manes.09G182500.1.p   RNGNYVECLLCVSKKLDREGAATGVFCFLQLASLELQQALHIQRLSEQTALKRLKTLAYI
Casp30444.t1          RNGKYVECLLCLSKKLNREGSVTGIFCFLQLASQELQQALHIQQLSEQIALKRLKTLAYM
XP_002512596.1        RNKKYVECLLCVSKKLDREGAVTGVFCFLQLASQELQQALHIQRLSEQTALKRLKTLAYI

                                910       920       930       940       950       960
AT1G09570.3           KRQIRNPLSGIMFTRKMIEGTELGPEQRRILQTSALCQKQLSKILDDSDLESIIEGCLDL
Ep_chr4_g08890.t1     KRQIQNPLAGVISAQKMLEGTRLDADQRKFLHSSALCQQQLSKIVESPDFDSITEGYLDL
KAF2324094.1          ............................................................
Potri.013G000300.2.p  KRQIWNPLSGIIFSGKMMEGTELGAEQKELLHTSAQCQCQLSKILDDSDLDSIIEGYLDL
Glyma.19G224200.1.p   KRQIRNPLYGIMFSRKLLEGTELGAEQKQFLQTGIRCQRQISKILDDSDLDSIIDGYMDL
Glyma.20G090000.3.p   KRQIRNPLCGIIFSRKMLEGTALGTEQKQLLRTSAQCQQQLSKILDDSDLDSIIDGYLDL
Glyma.10G141400.1.p   KRQIRNPLCGIVFSRKMLEGTDLGTEQKQLLRTSAQCQQQLSKILDDSDLDTIIDGYLDL
KAF2307360.1          ............................................................
Manes.09G182500.1.p   KRQIRNPLSGIIFSRKMMEATELDAEQNQLLQTSAQCQRQLGKILDDSDLDSIIEGYLDL
Casp30444.t1          KRQIQNPLAGIISAQKMLEDTKLDAEQKKFLGSSSLCQQQLSKVVDSPDFDSITEGYLDL
XP_002512596.1        KRQIQNPLSGIMFSRKLMEITELDAEQKQLLHTSAQCQRQLSKILDDSDIDSIVEGYLDL

                                970       980       990      1000      1010      1020
AT1G09570.3           EMKEFTLNEVLTASTSQVMMKSNGKSVRITNETGEEVMSDTLYGDSIRLQQVLADFMLMA
Ep_chr4_g08890.t1     EMIEFTLHEVLVASMSQVTTKVKAKGMQIVRDLSEEIMHEKLYGDSIRLQQVLADFLAVS
KAF2324094.1          ............................................................
Potri.013G000300.2.p  EMVEFTLREVLVAATSQVMMKSNEKGIRIINDAAEETMAETLYGDSIRLQQVLADFLQMS
Glyma.19G224200.1.p   EMVEFTLHEVLVASLSQVMTKSNAKGIRVVNDVEEKITTETLYGDSIRLQQVLADFLLIS
Glyma.20G090000.3.p   EMAEFTLHEVLVTSLSQVMTKSNGKSIRIVNDVAEQIVMETLYGDSLRLQQVLADFLLIS
Glyma.10G141400.1.p   EMAEFTLHEVLVTSLSQVMTKSNGKSIRIVNDVAGHIMMETLYGDSLRLQQVLADFLLIS
KAF2307360.1          ............................................................
Manes.09G182500.1.p   EMVEFTLREILVASISQVKMKSEGKGIQVSHDASEEIMTEILYGDSLRLQQVLSDFLLVS
Casp30444.t1          EMVEFTLHEVLIASFSQITTKVKAKGMQIVRDVTEEIMYEKLYGDSIRLQQVLADFLTVS
XP_002512596.1        EMVEFTLHEVLIAAISQVTIKSKGKGIRIVNDAAEVIMTETLYGDSIRLQQVLADFLAAS

                               1030      1040      1050      1060      1070      1080
AT1G09570.3           VNFTPSGGQLTVSASLRKDQLGRSVHLANLEIRLTHTGAGIPEFLLNQMFGTEEDVSEEG
Ep_chr4_g08890.t1     VNYAPTGGHLMIATKFTKDQLGKTVHLVHLELRITHAGNGIPEAVLSPMFGGDGGISEES
KAF2324094.1          ............................................................
Potri.013G000300.2.p  VNFTPSGGLLSVSASLTKDQLGQSVYLVHLELRIRHPGAGIPEALLDQMFGEDTDASVEG
Glyma.19G224200.1.p   INFTPTGGQVVVAATLTQQQLGKLVHLANLEFSITHDSFGVPETLLNQMFGRDGHESEEG
Glyma.20G090000.3.p   INFTPNGGQVVVAGTLTKEQLGKSVHLVKLELSITHGGSGVPEALLNQMFGNNGLESEEG
Glyma.10G141400.1.p   INFTPNGGQVVVAGSLTKEQLGKSVHLVKLELSITHGGSGVPEVLLNQMFGNNGLESEEG
KAF2307360.1          ............................................................
Manes.09G182500.1.p   VNYTPTGGQLIIATNLTKDQLGQSVQLVHLELRITHTGGGIPETLLNQMFGSDVDASDEG
Casp30444.t1          VNYAPTGGQLIFATNFTKDQLGKSVHLVHLEFRIMHAGNGIPEAVLSPMFGGDGGISEES
XP_002512596.1        VDFTPPGGQLTIAAKFTKDQLGQSVHLVHLELRITHAGGGIPEPLLNQMFGSDGDVSDEG
```

| | 1090 · 1100 · 1110 · 1120 |
|---|---|
| AT1G09570.3 | LSLMVSRKLVKLMNGDVQYLRQAGKSSFIITAELAAANKX |
| Ep_chr4_g08890.t1 | ISLFVSRKLVKLMNGDVQYLREMGKSTFIISLELAGGSRP |
| KAF2324094.1 | ........................................ |
| Potri.013G000300.2.p | ISLVISRKLVKLMNGDVRYMREAGKSSFIISVELAGGHKS |
| Glyma.19G224200.1.p | ISMLISRKLLKLMNGDVRYLREAGKSSFILSVELAAAHKS |
| Glyma.20G090000.3.p | ISLLISRKLLKLMNGDVRYLREAGKSAFILSAELAAAHNL |
| Glyma.10G141400.1.p | ISLLISRKLLKLMNGDVRYLREAGKSAFILSAELAAAHNL |
| KAF2307360.1 | ........................................ |
| Manes.09G182500.1.p | ISLLISRKLVKHMSGDVLYLREAGQSSFIISAELAAGCKP |
| Casp30444.t1 | ISLFVSRKLVKLMNGDVQYLREAGKSSFIISVELAGGSKP |
| XP_002512596.1 | VSLFISRKLVKLMNGDVQYLREAGKSSFIVTVELAAGRKS |

# PI

```
                   1         10        20        30        40        50        60
KAF2291335.1       MGRGKIEIRRIENPSNRQVTYSKRRNGIMKKAKEITVLCDAQVSLVIFASSGKMHEYCSP
AT5G20240.1        MGRGKIEIKRIENANNRVVTFSKRRNGLVKKAKEITVLCDAKVALIIFASNGKMIDYCCP
Casp33119.t1       MGRGKIEIKKIENVTNRQVTYSKRKNGIIKKAGEISVLCDAQVSLLIFSSSGKIHDYFSP
Ep_chr8_g25689.t1  MGRGKIEIKKIENTSNRQVTFVKRKNGIIKKAGEISKLCDAEVSLMIISSSGKIHDFFSP
Casp33120.t1       MGRGKIEIKRIDNPKNRHVTYSKRKNGIIKKAEQISILCDAKVSLLIFGTSGKIHQYCSP
Ep_chr8_g25690.t1  MGRGKIEIKRIDNQKNRNVTYSKRKDGIIKKAAQIAKLCDAEVSLIIFSSSGKVHEFCSP
Glyma.04G245500.2.p MGRGKIEIKRIENSSNRQVTYSKRKNGILKKAKEISVLCDAQVSLIIFGVSGKMHEYISP
Glyma.06G117600.1.p MGRGKIEIKRIENSSNRQVTYSKRKNGILKKAKEISVLCDAQVSLIIFGVSGKMHEYISP
KAF2296683.1       .............................MKKAKEITVLCDAKVSLVIFASSGKMHEYCSP
Potri.005G182200.1.p MGRGKIEIKRIENASNRQVTYSKRKNGIIKKAKEITVLCDAQVSLVIFASSGRMHEYCSP
Potri.002G079000.1.p MGRGKIEIKRIENSSNRQVTYSKRRSGIIKKAKEITVLCDAQVSLVIFASSGRMHEYCSP
XP_002520308.1     MGRGRIEIKRIENSSNRQVTYSKRRNGIIKKAKEITVLCDAEVSLIIFASSGKMHEYCSP
Manes.02G168500.1.p MGRGKIEIKRIENSSNRQVTYSKRRNGIIKKAKEITVLCDAQVSLVIFASSGKMHEYCSP
Manes.18G082400.3.p MGRGKIEIKRIENLSNRQVTYSKRRNGIMKKAKEITVLCDAKVSLIIFASSGKMHEYCSP
Glyma.13G034100.2.p MGRGKIEIKRIENSSNRQVTYSKRKNGILKKAKEITVLCDAQVSLIIFAASGKMHDYISP
Glyma.14G155100.1.p MGRGKIEIKRIENSSNRQVTYSKRKNGILKKAKEITVLCDAQVSLIIFAASGKMHDYISP

                         70        80        90        100       110       120
KAF2291335.1       STTLIDMLDKYHKQSGKRLWDAKHEDLSNEIDRIKKENDNMQIELRHLKGEDITSLHHEE
AT5G20240.1        SMDLGAMLDQYQKLSGKKLWDAKHENLSNEIDRIKKENDSLQLELRHLKGEDIQSLNLKN
Casp33119.t1       NAPLAQFLEDYQNKTNNKLWDAEHESLKNEIDRIKKENDNMEIELRHLKGLDIGSMHYLE
Ep_chr8_g25689.t1  KADLPQLLEDYQNKMKNKLWDAEHENLKNEIDRIKKENDSMEVHLRHLKGLDIASMHYSE
Casp33120.t1       STNLVDLLDNYQKLPDKRLWDAKHEELSKEIDRIKKENDSMKIKLRHLNGEDISSLHHTE
Ep_chr8_g25690.t1  NTSLVDLLDNYQKQPSKRLWDAKHENLNNEIERIKKENDRMKMKLRQLNGEDIGSLHHTE
Glyma.04G245500.2.p STTLIDVLDRYQRASGKTLWDAKHENLSNEIDRIKKENDSMQIELRHLKGEDITSLNYKE
Glyma.06G117600.1.p YTTLIDVLDRYQRASGKTLWDAKHENLSNEIDRLKKENDSMQIELRHLKGEDITSLNYKE
KAF2296683.1       STTLIDMLDMYHKQSGKRLWDAKHENLSNEIDRIKKENDNMQIELRHLEGQDISSLQPNE
Potri.005G182200.1.p STTVVDLLDKYHKQSGKRLWDAKHENLSKEIDRIKKENDSMQIELRHLKGEDISSLHHTE
Potri.002G079000.1.p STTVVDLLDKYHKQSGKRLWDAKHENLSNEIDRIKKENESMQIELRHLKGQDISSLPHKE
XP_002520308.1     STTLVDMLDKYHKLSGQRLWDAKHENLSNEIDRVKKENDNMQIELRHLNGEDISSLKYPE
Manes.02G168500.1.p STTLIDILDRYHKQSGKRLWDAKHENLSNEIDRIKKENDNMQIELRHLKGEDITSLHHQE
Manes.18G082400.3.p STTLVEILDMYHKHSGKRLWDAKHENLSNEIDRIKKENDNMQIELRHLKGEDISSLQHKE
Glyma.13G034100.2.p STTLIDILERYHKTSGKRLWDAKHENLNGEIERLKKENDSMQIELRHLKGDDINSLNYKE
Glyma.14G155100.1.p STTLIDILERYQKTSGKRIWDAKHENLNGEIERLKKENDSMQIELRHLKGEDINSLNYKE

                             130       140
KAF2291335.1       LGVIEEALEKGLATIRDKQASI................ELX...................
AT5G20240.1        LMAVEHAIEHGLDKVRDHQMEILISKRRNEKMMAEEQRQLTFQLQQQEMASNARGM....
Casp33119.t1       LMDIERSLEQGLDGISKQRIELRRIMDENVENLEEENNRLIFELQQKEMAEKMMENQNPY
Ep_chr8_g25689.t1  LMDIERSLQQGLDGIYDQRMGVKRMMEENIEILEVENNRLICELEQMEMVEKVREMENPY
Casp33120.t1       LQGIEQVLENGIAVVREKRMECYEITENNTKILEEEHRHLNFYLQQQEMAEKMQKMENPY
Ep_chr8_g25690.t1  LGGIEQVLENGLAIVRDKRMECYEMTQNNTKFLEEENRHLNFCLQQQERQ....KLENPY
Glyma.04G245500.2.p LMALEDALENGLSGVREKKMEVHRMFKRNDKILEEQNKELNFLLQQHLALEGVGNM....
Glyma.06G117600.1.p LMALEDALENGLSGVREKKMEVHRMFKRNGKILEEQNKELSFLLQQHLAVEGIGNM....
KAF2296683.1       LLGIEHALENGLACVREKKMEFYKMKKKNEMNLEEENKHLRFILQQQEMAENMREMENPY
Potri.005G182200.1.p LMAIEEALDAGLAAVRKKQMEYHSMLEQNEKMLDEEFKRLQFVLQQQEMAENAMEMENAY
Potri.002G079000.1.p LMAIEEALDTGLAAVRKKQMEFHSMLEQNEKILDEEFKHLQFVLQQQEMAENAMEMENAY
XP_002520308.1     LQALEDALETGLAGIRNKQMEYYKMMKRKDKSLADENQHLSFILHQLEMEQNMREMENPY
Manes.02G168500.1.p LMVIEEALDNGLASVRDKMMEYYNMKKTNDEILEDMNKRLRFFWQQQEMAENVREMENPY
Manes.18G082400.3.p LMAIEEALDHGLASVRHKQMEYYKMKKRNDKLLEDENKELRFVLQQHEMDEHMREMENPY
Glyma.13G034100.2.p LMALEDALETGLVSVREKQMDVYRMLRRNDKILEEENRELNFLWQQRLAEEGAREVDNGF
Glyma.14G155100.1.p LMALEDALETGLVSVREKQMDVYRMFRRNDKILEEENRELTFLWQQRLAVEGAREVDNGF

KAF2291335.1       ..............................
AT5G20240.1        ...MMRDHDGQ..FGYRVQPIQPNLQEKIM
Casp33119.t1       HQYNMKEYNSPMPFAFRIQPIQPNLRERMX
Ep_chr8_g25689.t1  HHHNPMN....MPFAFRIKPTHPDLRERIX
Casp33120.t1       HHQQLNNYNSHMPFAFPVQPIQPNLQDIKX
Ep_chr8_g25690.t1  HHQQKMEYSCNLTFAFSIKPWK.NVX....
Glyma.04G245500.2.p HGQWIX........................
Glyma.06G117600.1.p HEQWIX........................
KAF2296683.1       HQQRVXEYNSQMPFAFRVQPIQPNLQERMX
Potri.005G182200.1.p HQQRMRDYNFQVPFAFRVQPIQPNLQERMX
Potri.002G079000.1.p HQQRVRDYNSQVPLAFRVQPIQPNLQERMX
XP_002520308.1     HQQRVRDYNSQVPFAFRVQPIQPNLQGRMX
Manes.02G168500.1.p HQQRVRDYNSQMPFAFRVQPIQPNLQERMX
Manes.18G082400.3.p HQQRMRDFNYQMPFAFRVQPIQPNLQDRIX
Glyma.13G034100.2.p DQS.VRDYNSHMPFAFRVQPMQPNLQERIX
Glyma.14G155100.1.p DQS.VRDYNSHMPFAFRVQPMQPNLQERIX
```

### SEP1

```
                          1        10        20        30        40        50        60
AT5G15800.2               MGRGRVELKRIENKINRQVTFAKRRNGLLKKAYELSVLCDAEVALIIFSNRGKLYEFCSS
XP_002525916.2            MGRGRVELKRIENKINRQVTFAKRRNGLLKKAYELSVLCDAEVALIIFSNRGKLYEFSSI
Ep_chr2_g04449.t1         MGRGRVELKRIENKINRQVTFAKRRNGLLKKAYELSVLCDAEVALIIFSNRGKLYEFCSS
XP_015583405.1            MGRGRVELKRIENKINRQVTFAKRRNGLLKKAYELSVLCDAEVALIIFSNRGKLYEFCSS
AT2G03710.1               MGRGKVELKRIENKINRQVTFAKRRNGLLKKAYELSVLCDAEIALLIFSNRGKLYEFCSS
AT3G02310.1               MGRGRVELKRIENKINRQVTFAKRRNGLLKKAYELSVLCDAEVSLIVFSNRGKLYEFCST
Ep_chr3_g07686.t1         MGRGRVELKRIENKINRQVTFAKRRNGLLKKAYELSVLCDAEVALIIFSNRGKLYEFSSI
Potri.004G115500.7.p      MGRGRVELKRIENKINRQVTFAKRRNGLLKKAYELSVLCDAEVALIIFSNRGKLYEFCST
KAF2299250.1              MGRGRVELKRIENKINRQVTFAKRRNGLLKKAYELSVLCDAEVALIIFSNRGKLYEFSSI
Glyma.18G273500.1.p       MGRGRVELKRIENKINRQVTFAKRRNGLLKKAYELSVLCDAEVALIIFSNRGKLYEFCSS
Casp01545.t1              ............................................................
Manes.01G025700.1.p       MGRGRVELKRIENKINRQVTFAKRRNGLLKKAYELSVLCDAEVALIVFSNRGKLYEFCST
Casp08433.t1              MGRGRVELKRIENKIN.................................SRGKLFEFCST
Ep_chr6_g17596.t1         MGRGRVELKRIENKINRQVTFAKRRNGLLKKAYELSILCDAEVALIIFSNRGKLFEFSSS
KAF2298928.1              MGRGRVELKRIENKINRQVTFAKRRNGLLKKAYELSVLCDAEVALIIFSNRGKLYEFCST
KAF2298953.1              MGRGRVELKRIENKINRQVTFAKRRNGLLKKAYELSVLCDAEVALIIFSNRGKLYEFCST
Casp07062.t1              ............................................................
Glyma.17G080900.1.p       MGRGRVELKRIENKINRQVTFAKRRNGLLKKAYELSVLCDAEVALIIFSNRGKLYEFSST
Glyma.05G018900.1.p       MGRGRVELKRIENKINRQVTFAKRRNGLLKKAYELSVLCDAEVALIIFSNRGKLYEFSST
Glyma.02G121500.1.p       MGRGKVELKRIENKINRQVTFAKRRNGLLKKAYELSVLCDAEVALIIFSNRGKLYEFCSG
Glyma.01G064100.1.p       MGRGKVELKRIENKINRQVTFAKRRNGLLKKAYELSVLCDAEVALIIFSNRGKLYEFCSG
Potri.017G099700.2.p      MGRGRVELKRIENKINRQVTFAKRRNGLLKKAYELSVLCEAEVALIIFSNRGKLFEFCST
Glyma.08G250700.1.p       MGRGRVELKRIENKINRQVTFAKRRNGLLKKAYELSVLCDAEVALIIFSNRGKLYEFCSS
Glyma.13G052700.1.p       MGRGRVELKRIENKINRQVTFAKRRNGLLKKAYELSVLCDAEVALIIFSTRGKLYEFCST
Glyma.19G034500.1.p       MGRGRVELKRIENKINRQVTFAKRRNGLLKKAYELSVLCDAEVALIIFSTRGKLYEFCST
Manes.02G059200.2.p       MGRGRVELKRIENKINRQVTFAKRRNGLLKKAYELSVLCDAEVALIIFSNRGKLYEFSSI
Manes.01G103100.2.p       MGRGRVELKRIENKINRQVTFAKRRNGLLKKAYELSVLCDAEVALIVFSNRGKLYEFSSI
KAF2313699.1              ............................................................
Manes.06G081900.2.p       MGRGKVVLKRIENKINRQVTFAKRRNGLLKKAYELSVLCDAEIALIIFSTRGKLYEFCST
KAF2312882.1              ............................................................
KAF2282965.1              MGRGKVELKRIENKINRQVTFAKRRNGLLKKAYELSVLCDAEVALIIFSNRGKLYEFCST
Manes.14G088400.1.p       MGRGKVELKRIENKINRQVTFAKRRNGLLKKAYELSVLCDAEVALIIFSNRGKLYEFCSS
Potri.008G098400.2.p      MGRGRVELKRIENKINRQVTFAKRRNGLLKKAYELSVLCDAEVALIIFSNSGKLFEFCSS
Manes.05G111950.1.p       MGRGRVELKRIENKINRQVTFAKRRNGLLKKAYELSVLCDAEVALIIFSNRGKLYEFCST
```

```
                                   70        80        90       100       110       120
AT5G15800.2               SNMLKTLDRYQKCSYGSIEVNKPAKELENSYREYLKLKGRYENLQRQQRNLLGEDLGPLN
XP_002525916.2            SNMLKTLERYQKCSYGAVEVNKPAKELESSYREYLKLKTRFESLQKTQRNLLGEDLGPLS
Ep_chr2_g04449.t1         PSMTKTIDKYQRCSYGTLETNKSMLETQSCYQEYLKLKARVEALQRSQRNFLGEDLGQLH
XP_015583405.1            PSMAKTIEKYQRCSYGALEANQSVHDTQNSYQEYLKLKSRVEALQRSQRHFLGEDLGNLG
AT2G03710.1               SGMARTVDKYRKHSYATMDPNQSAKDLQDKYQDYLKLKSRVEILQHSQRHLLGEELSEMD
AT3G02310.1               SNMLKTLERYQKCSYGSIEVNKPAKELENSYREYLKLKGRYENLQRQQRNLLGEDLGPLN
Ep_chr3_g07686.t1         SNMLKTLDRYQKCSYGAVEVNRPAKELESSYREYLKLKARCESLQRTQRNLLGEDLGPLN
Potri.004G115500.7.p      SNMLKTLERYQKCSYGAEEVNKPAKELESSYREYLKVKARFEALQRTQRNLLGEDLGPLN
KAF2299250.1              SNQLQYPDK..TCSFHLVLVSAPWKGIKSA....AMVRWKSTDLPRSSRNLLGEELGPLS
Glyma.18G273500.1.p       SSMLKTLERYQKCSYGAVEVSKPAKELQSSYREYLKLKARFESLQRTQRNLLGEDLGPLN
Casp01545.t1              ..MTKTIDKYQRCSYGSLETNKSMLETQSCYQEYLKLKAKVEALQRSQRNFLGEDLGHLH
Manes.01G025700.1.p       SSMAKTIDKYQRCSYAPVESNQSMHDTQSCYQEYLKLKAKVEMLQRSQRNLLGEELGDLN
Casp08433.t1              SSMVKTLERYHKSSYGGLEATQPSNETQSNYQDYLKLKARVEVLHRSQRNLLGEDLGALT
Ep_chr6_g17596.t1         SSMVSTLERYHKCSFGGVEARQPSRDTQGNYQDYLKLKARVEVLQRSQRNLLGEDLEALS
KAF2298928.1              SSMAKTIEKYHRCSYAPLEANQSMHDIQNCYQEYLKLKEKVEALQRSQRNLLGEDLGDLN
KAF2298953.1              SSMAKTIEKYHRCSYAPLEANQSMHDIQNCYQEYLKLKEKVEALQRSQRNLLGEDLGDLN
Casp07062.t1              ..MLKTLDRYQKCSYGAVEVNKPAKELESSYREYLKLKARCESLQRTQRNLLGEDLGPLN
Glyma.17G080900.1.p       SSMMKTLEKYQKYSYSALETTRPINDSQN.YQEYLRLKARVEVLQCSQRNLLGEDLAQMN
Glyma.05G018900.1.p       SSMMKTLEKYQKYSYSALETTRPINDTQN.YQEYLRLKARVEVLQRSQRNLLGEDLAQMN
Glyma.02G121500.1.p       HSMAKTLERYHRCSYGALEVQQPEIETQRRYQEYLKLKSRVEALQQTQRNLLGEELEHLD
Glyma.01G064100.1.p       HSTAKTLERYHRCSYGALEVQQPEIETQRRYQEYLKLKSRVEALQQTQRNLLGEELEHLD
Potri.017G099700.2.p      SNMLKTLERYQKCSYGAEEVNKPAKELESSYREYLKVKAKFETLQRTQRNLLGEDLGPLN
Glyma.08G250700.1.p       SSMLKTLERYQKCSYGAVEVTKPAKELQSSYREYLKLKARFESLQRTQRNLLGEDLGPLN
Glyma.13G052700.1.p       NSMLKTLERYQKCSYGAVEVSKPGKELQSSYREYLKLKARFESLQRTQRNLLGEDLGPLN
Glyma.19G034500.1.p       NSMLKTLERYQKCSYGAVEVSKPGKELQSSYREYLKLKARFESLQRTQRNLLGEDLGPLN
Manes.02G059200.2.p       SNMLKTLDRYQKCSYGAVEVNKPAKELESSYREYLKLKAKFESLQRTQRNLLGEDLGPLN
Manes.01G103100.2.p       SNMLKTLERYQKCSYGAVEVNRPAKELESSYREYLKLKGRFESLQRTQRNLLGEELGPLN
KAF2313699.1              ..MLQTLERYQKCSYGAVEVNKPAKELESSYREYLKLKAKFESLQRTQRNLLGEDLGPLN
Manes.06G081900.2.p       SSMMRTLERYQKCSYGAMEASYPGYETQSNHQEYLKLKARFEVLQRSQRNLLGDELGPLN
KAF2312882.1              ..MMRTLERYQKCSYGGLEASQPGYETQSNYQEYLKLKARVEVLQRSQRNLLGEDLGPLN
KAF2282965.1              SSMMRTLERYQKCSYGALEASQPGHETQSNYQEYLKLQARVEVLQRSQRHLLGEDLGPLN
Manes.14G088400.1.p       SSMMRTLERYQKCSYGASEACQPGFETQSNYQEYLKLKARVEILQRSQRNLLGEDLGPLN
Potri.008G098400.2.p      SNMATTIEKYQRFSYGALEGGQSEKETQNNYQEYLKLKTRVDVLQRSQRNLLGEDLGNLG
Manes.05G111950.1.p       SSMAKTIEKYQRCSYAPLGNDQSVHDTQSCYQEYLKLKAKVEVLQHSQRNLHGENLGDLN
```

```
                        130       140       150       160       170       180
AT5G15800.2          SKELEQLERQLDGSLKQVRSIKTQYMLDQLSDLQNKEQMLLETNRALAMKLDDMIGVGGG
XP_002525916.2       TKELEQLERQLESSLKLVRSTRTQFMLDQLSDLQNKEQLLLEANRALTLKLDEISARIRS
Ep_chr2_g04449.t1    TDELEHLETQLDSSLKQIRLNKTTFLLNQLSELQKKEEVLLETNNTLRKKLQETNASMEP
XP_015583405.1       TKDLEQLEHQLDSSLKHVRLTKSNFMLDQLSQLQRKEEMLLQTNNALRKKLEETNAALQP
AT2G03710.1          VNELEHLERQVDASLRQIRSTKARSMLDQLSDLKTKEEMLLETNRDLRRKLEDSDAATQS
AT3G02310.1          SKELEQLERQLDGSLKQVRCIKTQYMLDQLSDLQGKEHILLDANRALSMKLEDMIGVIGG
Ep_chr3_g07686.t1    SKDLEQLERQLESSLKQVRSTKTQYMLDQLSDLQSKEQVLLESNRALQIKLEDISARLRI
Potri.004G115500.7.p TKELEQLERQLESSLNQVRSTKTQYMLDQLADLQNKEHLLLEANRGLTIKLDEISARLRP
KAF2299250.1         TKELEQLERQLESSLKQVRSTKTQFMLDQLSDLQNKEQLLLESNRALAIKLDEISASLRS
Glyma.18G273500.1.p  IKELEHLERQLDSSLKQVRSTKTQFMLDQLSDLQTKEQMLVEANRSLTVKLEEINSRYRQ
Casp01545.t1         TDELEQLECQLDSSLKQIRVTKSTFLLNQLSELQRKEEVLLETNNALRKKLEETNASIES
Manes.01G025700.1.p  TKELKQLEHQLDSSLKQIRSTKTQFMVDQLSELQRKEELLLETNHALRKKLEETDAALQS
Casp08433.t1         ARELEQLEDQLETSLKQIRSTMTQTMVDQLAVLQNREQQLVEVNKALKMKLHGNNSQGGV
Ep_chr6_g17596.t1    GRELEQLEDQLETSLKLIRSTMTQSMLDEVANLQNREQQLVEVNKALKMKLHGSNYQSGV
KAF2298928.1         TTDLKQLEHQLDSSLKQIRSTKTQFMLDRLSELQRKEESLLETNNALRKKLEETDAALQS
KAF2298953.1         TTDLKQLEHQLDSSLKQIRSTKTQFMLDRLSELQRKEESLLETNNALRKKLEETDAALQS
Casp07062.t1         SKDLEQLERQLESSLKQVRSTKTQYMLDQLSDLQSKEQVLLESNRALQIKLEEISARLRI
Glyma.17G080900.1.p  TNELEQLENQLETALKNIRSTKTQFMLDQLSDLHHRETLLIETNNVLRSKLEETNSQVSL
Glyma.05G018900.1.p  TNELEQLENQLEAALRNIRSTKTQFMLDQLSDLHHRETLLIETNNVLRSKLEETDSQVSL
Glyma.02G121500.1.p  VKDLEQLERQLDSSLKQIRSNKTQQMLDQLADLHRKEEMLLETNNILRNKLEEINVALQP
Glyma.01G064100.1.p  VNDLEQLERQLDSSLKQIRSNKTQQMLDQLSDLHRKEEMLLETNNILRNKLEEINVALQP
Potri.017G099700.2.p TKELEQLERHLESSLKQVRSTKTQYMLDQLGDLQNKEHMLLEANRALTIKLDEISARLRP
Glyma.08G250700.1.p  TKELEHLERQLDSSLKQVRSTKTQFMLDQLSDLQTKEQMLVEANRSLTVKLEEINSRYRQ
Glyma.13G052700.1.p  TKDLEQLERQLDSSLKQVRSTKTQFMLDQLADLQNKEHMLVEANRSLTMKLEEINSRYRQ
Glyma.19G034500.1.p  TKDLEQLERQLDSSLKQVRSTKTQFMLDQLADLQNKEHMLVEANRSLTMKLEEINSRYRQ
Manes.02G059200.2.p  TKELEQIERQLESSLKQVRSTRTQFMLDQLSDLQNKEQLLLEANRALTIKLDEISARLRS
Manes.01G103100.2.p  TKELEQLERQLESSLKQVRSTKTQFMLDQLSDLQKKEQLLLEANRALAMKLDEISASLRS
KAF2313699.1         TKELEQIERHLESSLKQVRSTRTQFMLDQLSDLQNKEQLLLEANRALTIKLDEISARLRS
Manes.06G081900.2.p  TKELEQLELQLETSLKRIRFTKTQFMLHQLADLQNKEYLLLETNKALKRKLEEGSGQL.L
KAF2312882.1         TKELEQLELQLETSLKQIRLTMTQFMLDQLADLQNKERLLIETNKALKRKLEEGSGQLRL
KAF2282965.1         TKELEQLEHQLETSLNQIRSKKTQFMLDQLVDLQNREHLLLETNKALKRKLEEGSGQLLL
Manes.14G088400.1.p  TKELEQLEHQLESSLNQIRSTKTQFMLDQLADLQNREHVLLETNKALKRRLEEGSGQLRL
Potri.008G098400.2.p TMELDQLENQLDSSLKQIRSRKGQFVLDELSELQRKEELLLETNNALKRKLEETSAAIRL
Manes.05G111950.1.p  IKDLQQLELQLDSSLKKIRSTKTQFMLDQLSELQRKEELLLQTNKALRKKLVETDAALQS

                        190       200       210       220       230       240
AT5G15800.2          GWEGGEQNVTYAHHQAQSQGLYQPLECNPTLQMGYNPVCSEQITATTQAQQGNGYIPGWM
XP_002525916.2       SWEGGEQSMSYGQQHPQPQELFQPMDCNPTLQIGYNPVGSDQMTATTHAQTVSGFIPGWM
Ep_chr2_g04449.t1    AWEARNHNAGFGHQNMETDDFPDHLECNGHMQISYTAGGTDQVTVASTGQNINGFIPGWM
XP_015583405.1       PWEARDESIPYNRQPGQSSEGFDPLQCSSHFRTGAGE..TDPVTVANTSENINGFIPDWM
AT2G03710.1          FWSAAEQQQQHQQQPIQEAGFFKPLQGNVALQMSYNHNPANATNSATTSQNVNGFFPGWM
AT3G02310.1          GWEGGDQNIAYGHPQAHSQGLYQSLECDPTLQIGYSPVCSEQMAVTVQSQQGNGYIPGWM
Ep_chr3_g07686.t1    SWEGGEQSMSYGQQHPQSQELFQPLECNPTLQIGYNPVGSDQMTATTHAQQVNGYIPGWM
Potri.004G115500.7.p SWEGDDQNMSYGHQHAQSQGLFQALECNPTLQIGYNPVGSDQMTATTHAQQVHGFIPGWM
KAF2299250.1         SWEGGEQSMSYGQQHPHSQELFQPLECNPTLQIGYN...SDQMTATTHAQQVNGFIPGWM
Glyma.18G273500.1.p  SWEAGDQSMPYGHSHSHSQGFFQPLECNPTLHIGYNAVASDQITATTQPQQVSGFIPGWM
Casp01545.t1         TWEARNQNVCFNRQQVHSDDYPNPLECNGHMQISYSNGEPDQVTVASTCQNINGFIPGWM
Manes.01G025700.1.p  SWATREQNVHYNHHPAQPGDFVNPLQCNRNFRIGFNAGETGQVTVATTEQNFNGFIPGWM
Casp08433.t1         SWESETHNFPYPRLLPHSEAFFHTL......QIGYNPMGADHEVGAAEAQHVNGFVPGWM
Ep_chr6_g17596.t1    SWE....TQTYPPLLPHSQAFFHTL......PIGYNPMGADH.QANGGAAHVNGFLPGWM
KAF2298928.1         SWEAREQCVQYNHQPTQPDDFANPLQCNRNLRIGFVFVPSVQKIITKPLESISGLMPGAI
KAF2298953.1         SWEAREQCVQYNHQPTQPDDFANPLQCNRNLRIGFVFVPSVQKIITKPLESISGLMPGAI
Casp07062.t1         SWEGGEQSMSYGQQQQQSQELFQPLECNPTLQIGYNPVGSDQMTATSHAQQVNGYIPGWM
Glyma.17G080900.1.p  ALEAGGPSIQYTNFPPQSEGFFQPMGVNPTLQIGYNQTNPHDANVGASSLSMHGFASEWM
Glyma.05G018900.1.p  ALEAGGPSIQYTNFPPQSEGFFEPVGVNPTLQIGYNQTGPDDTNVGASSLSMHGFASGWM
Glyma.02G121500.1.p  TWETREQNAPYNYHPSQSEGYYETAHCNSTLRIGYDSSGLNEAAAGTSAQNASEFMNGWM
Glyma.01G064100.1.p  TWEAREQNAPYNCHPPQTEGYYETAHCSSTLRIGYDSSGLNEAAAGASAQNASEFMHGWM
Potri.017G099700.2.p SWEGDDQSMSYGHQHAQSQGLFQHLECNPTLQIGYNSVGSDQIAATHAAQQVHGFIPGWM
Glyma.08G250700.1.p  SWEAGDQSMPYGPQNSHSQGFFQPLECNPTLQIGYNDVASDQITATTQPQQVSGFIPGWM
Glyma.13G052700.1.p  TWEAGEQSMSYGTQNAHSQGFFQPLECNPTLQIGYNPEASEQLAATTQAQQVNGFIPGWM
Glyma.19G034500.1.p  TWEAGEQSMPYGTQNAHSQGFFQPLECNPTLQIGYIPEASEQLAATTQAQQVNGFIPGWM
Manes.02G059200.2.p  SWEGGEQSMSYGQQHPQSQELFQPLECNPTLQIGYNPVGTDQMNATTHAQQVNGFIPGWM
Manes.01G103100.2.p  SWEGGEQSMSYGQQHPQSQELFQPLECNPTLQIGYNPVGTDQMSDTTHSQQVNGFIPGWM
KAF2313699.1         SWESGEQSMPYGQQHTQSQELFQPLECNPTLQIGYNPVGTDQMIATTHAQQVNGFIPGWM
Manes.06G081900.2.p  AWEG.........LAAGSDGFFQP......LQIGYNP....EVNVV.HTQHVNGFIPGWM
KAF2312882.1         AWEGGGHTIPYSRLPAHSDGFFQPKGGNSTLQIGYNPEGAEEDNVS.HTHDLNGFIPGFM
KAF2282965.1         AWEDGGQTIPYSRLPAHSEGFFHPLGGNSTLQIGYNPVGADEINVA.HTQHVNGFIPGWM
Manes.14G088400.1.p  AWEGGGQTIPYSRLAVHSEGLFQPLGGNSTLQIGYNPVGADEVNLA.HTQHVNGFTPGWM
Potri.008G098400.2.p SWKVGEQRVPYSFQPVQPYDPIEPLQYNSTFQFGYNPAETDQATVTSSSQNVNGFIPGWM
Manes.05G111950.1.p  SWEASDQSAQYDRQPAQPDGLLDPLQCNRNLRIGFSPGEADELPVATTDQNVNGFIPGWM
```

| | | |
|---|---|---|
| **AT5G15800.2** | L | X |
| **XP_002525916.2** | L | X |
| **Ep_chr2_g04449.t1** | L | X |
| **XP_015583405.1** | L | X |
| **AT2G03710.1** | V | X |
| **AT3G02310.1** | L | X |
| **Ep_chr3_g07686.t1** | L | X |
| **Potri.004G115500.7.p** | L | X |
| **KAF2299250.1** | L | X |
| **Glyma.18G273500.1.p** | L | X |
| **Casp01545.t1** | L | X |
| **Manes.01G025700.1.p** | L | X |
| **Casp08433.t1** | L | X |
| **Ep_chr6_g17596.t1** | L | X |
| **KAF2298928.1** | L | A |
| **KAF2298953.1** | L | A |
| **Casp07062.t1** | L | X |
| **Glyma.17G080900.1.p** | L | X |
| **Glyma.05G018900.1.p** | L | X |
| **Glyma.02G121500.1.p** | N | X |
| **Glyma.01G064100.1.p** | N | X |
| **Potri.017G099700.2.p** | L | X |
| **Glyma.08G250700.1.p** | L | X |
| **Glyma.13G052700.1.p** | L | X |
| **Glyma.19G034500.1.p** | L | X |
| **Manes.02G059200.2.p** | L | X |
| **Manes.01G103100.2.p** | L | X |
| **KAF2313699.1** | L | X |
| **Manes.06G081900.2.p** | L | X |
| **KAF2312882.1** | L | X |
| **KAF2282965.1** | L | X |
| **Manes.14G088400.1.p** | L | X |
| **Potri.008G098400.2.p** | L | X |
| **Manes.05G111950.1.p** | L | X |

### SEP3

```
                    1        10        20        30        40        50        60
KAF2292923.1        MGRGRVELKRIENKINRQVTFAKRRNGLLKKAYELSVLCDAEVALIIFSNRGKLYEFCSS
KAF2292906.1        MGRGRVELKRIENKINRQVTFAKRRNGLLKKAYELSVLCDAEVALIIFSNRGKLYEFCSS
AT1G24260.2         MGRGRVELKRIENKINRQVTFAKRRNGLLKKAYELSVLCDAEVALIIFSNRGKLYEFCSS
Casp18261.t1        ............................................................
Ep_chr6_g20383.t1   MGRGRVELKRIENKINRQVTFAKRRNGLLKKAYELSVLCDAEVALIIFSNRGKLYEFCSS
Manes.13G009600.2.p MGRGRVELKRIENKINRQVTFAKRRNGLLKKAYELSVLCDAEVALIIFSNRGKLYEFCSS
KAF2286473.1        MGRGRVELKRIENKINRQVTFAKRRNGLLKKAYELSVLCDAE..................
Manes.12G009200.2.p MGRGRVELKRIENKINRQVTFAKRRNGLLKKAYELSVLCDAEVALIIFSNRGKLYEFCSS
XP_015572264.1      MGRGRVELKRIENKINRQVTFAKRRNGLLKKAYELSVLCDAEVALIIFSNRGKLYEFCSS
Potri.003G169600.1.p MGRGRVELKRIENKINRQVTFAKRRNGLLKKAYELSVLCDAEIALIIFSNRGKLYEFCSS
Potri.001G058400.1.p MGRGRVELKRIENKINRQVTFAKRRNGLLKKAYELSVLCDAEVALIIFSNRGKLYEFCSS
Glyma.08G105500.1.p MGRGRVELKRIENKINRQVTFAKRRNGLLKKAYELSVLCDAEVALIIFSNRGKLYEFCSS
Glyma.05G148800.1.p MGRGRVELKRIENKINRQVTFAKRRNGLLKKAYELSVLCDAEVALIIFSNRGKLYEFCSS
Glyma.11G252300.1.p MGRGRVELKRIENKINRQVTFAKRRNGLLKKAYELSVLCDAEVALIIFSNRGKQYEFCSG
Glyma.18G004700.4.p MGRGRVELKRIENKINRQVTFAKRRNGLLKKAYELSVLCDAEVALIIFSNRGKQYEFCSG

                             70        80        90       100       110       120
KAF2292923.1        SSMLKTLERYQKCNYGAPETNISTREALELSSQQEYLKLKARYEALQRSQRNLLGEDLGP
KAF2292906.1        SSMLKTLERYQKCNYGAPETNISTREALELSSQQEYLKLKARYEALQRSQRNLLGEDLGP
AT1G24260.2         SSMLRTLERYQKCNYGAPEPNVPSREALELSSQQEYLKLKERYDALQRTQRNLLGEDLGP
Casp18261.t1        ..MLKTLERYQKCNYGAPETNVSAREALELSSQQEYLKLKARYEALQRSQRNLLGEDLGP
Ep_chr6_g20383.t1   SSMLKTLERYQKCNYGAPETNVSAREALELSSQQEYLKLKARYEALQRSQRNLLGEDLGP
Manes.13G009600.2.p SSMLKTLERYQKCNYGAPETNVSAREALELSSQQEYLKLKARYEALQRSQRNLLGEDLGP
KAF2286473.1        ............................ELSSQQEYLKLKARYEALQRSQRNLLGEDLGP
Manes.12G009200.2.p SSMLKTLERYQKCNYGAPETNVSAREALELSSQQEYLKLKARYEALQRSQRNLLGEDLGP
XP_015572264.1      SSMLKTLERYQKCNYGAPEPNVSAREALELSSQQEYLKLKARYEALQRSQRNLLGEDLGP
Potri.003G169600.1.p SSMLKTLERYQKCNYGAPEPNVSAREALELSSQQEYLKLKARYEALQRSQRNLLGEDLGP
Potri.001G058400.1.p SSMLKTLERYQKCNYGAPEPNVSAREALELSSQQEYLKLKARYEALQRTQRNLLGEELGP
Glyma.08G105500.1.p SSMLKTLERYQKCNYGAPEANVSTREALELSSQQEYLKLKARYESLQRSQRNLMGEDLGP
Glyma.05G148800.1.p SSMLKTLERYQKCNYGAPEANVSTREALELSSQQEYLKLKARYEALQRSQRNLMGEDLGP
Glyma.11G252300.1.p SSMLKTLERYQKCNYGAPEDNVATNEALELSSQQEYLRLKARYEALQRSQRNLMGEDLGP
Glyma.18G004700.4.p SSMLKTLERYQKCNYGAPEDNVATKEALELSSQQEYLRLKARYEALQRSQRNLMGEDLGP

                            130       140       150       160       170       180
KAF2292923.1        LSSKELESLERQLDMSLKQIRSTRTQCMLDQLTDLQRKEHMLNEANKTLKQRLVEGYQIN
KAF2292906.1        LSSKELESLERQLDMSLKQIRSTRTQCMLDQLTDLQRKEHMLNEANKTLKQRLVEGYQIN
AT1G24260.2         LSTKELESLERQLDSSLKQIRALRTQFMLDQLNDLQSKERMLTETNKTLRLRLADGYQM.
Casp18261.t1        LNSKELESLERQLDMSLKQIRSTRTQFMLDQLTDLQRREHMLNEANRSLKQRLMEGYQVN
Ep_chr6_g20383.t1   LSSKELESLERQLDMSLKQIRSTRTQYMLDQLTDLQRREHMLNEANRSLKQRLVEGYQIN
Manes.13G009600.2.p LSSKELESIERQLDMSLKQIRSTRTQCMLDQLTDLQRKEHMLNEANKTLKQRLVEGYQIN
KAF2286473.1        LSSKELESLERQLDMSLKQIRSTRTQYMLDQLTDLQRKEHMLNEANKTLKQRLVEGYQVN
Manes.12G009200.2.p LSSKELESLERQLDMSLKQIRSTRTQYMLDQLTDLQRKEQMLNEANKTLKQRLVEGYQIN
XP_015572264.1      LSSKDLESLERQLDMSLKQIRSTRTQYMLDQLTDLQRKEHMLNEANKTLKQRLVEGYQVN
Potri.003G169600.1.p LSSKELESLERQLDMSLKQIRSTRTQYMLDQLNDLQHKEHMLTAANKSLRERLMEGYEVN
Potri.001G058400.1.p LSSKELESLERQLDMSLKQIRSTRTQYMLDQLHDLQHKEHMLTAANKSLKERLMEGYQLN
Glyma.08G105500.1.p LSSKELESLERQLDSSLKQIRSTRTQFMLDQLSDLQRKEHLLSEANRSLRQRQLEGYQIN
Glyma.05G148800.1.p LSSKELESLERQLDSSLKQIRSTRTQFMLDQLSDLQRKEHLLSEANRSLRQRQLEGYQIN
Glyma.11G252300.1.p LSSKELESLERQLDSSLKQIRSIRTQFMLDQLSDLQRKEHFLGESNRDLIQRL.EEFQIN
Glyma.18G004700.4.p LSSKELESLERQLDSSLKQIRSIRTQFMLDQLSDLQRKEHFLGESNRDLRQRL.EEFQIN

                            190       200       210       220       230
KAF2292923.1        SMQLNPSAEEVEFGRQAAQPQGDG.FHPLGCEPTLQIGFRAMLVTVMTAGPQREKDMRRI
KAF2292906.1        SMQLNPSAEEVEFGGQAAQPQGDGFFHPLGCEPTLQIGYAX...................
AT1G24260.2         PLQLNPNQEEVHYGRHQQQQHSQAFFQPLECEPILQIGYQGQQ.DGMGAGPSVNNYMLGW
Casp18261.t1        PLQLT.GQDEVAFGRQGGQSQGEGFFHPLECEPTLQIGYPHDPITVVTAGPSVNNYMPGW
Ep_chr6_g20383.t1   PMQLT.GQEEVGFGRQGGQPQGDGFYHPLECEPTLQIGYPHDQMTVVTAGPSVNNYMPGW
Manes.13G009600.2.p SMQLNPSAEDVGFGRQEAQPQGDGFFHPLDCEPTLQIGYQHDQITVVTAGPSMNNYMPGW
KAF2286473.1        SMQLNPSAEDVGFGRQAA..HGDVFFHALECEPTLQIGYQPDPITVVTAGPSVNNYMPGW
Manes.12G009200.2.p TMQLNPSAEEVAFGRQAPQHHGDVFFHPLECEPTLQIGYQPDPITVVTAGPSVNNYMPGW
XP_015572264.1      AMQLNPSAEDVGFGRQAAQPQGDGFFHPLDCEPTLQIGYHPDQI.VVTAGPSVNNYMSGW
Potri.003G169600.1.p SLQLNLSAEDVGFSRQQAQPQGYGFFHPLECEPTLQIGYQPDSITVVTSGPSMTAYMPGW
Potri.001G058400.1.p SLQLNPSAEDVEYARQQAQPQGDGFFHALECEPTLQIGYQPENITMVTAGPSMTTYMPGW
Glyma.08G105500.1.p PLQLNPGVEEMGYGRNPAQTHGEALFQQMECEPTLQIGYQPDPVSVVTAGPSMSNYMAGW
Glyma.05G148800.1.p PLQLNPGVEEMGYGRHPAQTHGEALFQQMECEPTLQIGYQPDPVSVVTAGPSMSNYMAGW
Glyma.11G252300.1.p PLQLNPSAEEMGHGRYPGQPQGHALFQPLDCEPTLQIGYHPDPVSVVSEGPSMNNYMAGW
Glyma.18G004700.4.p PLQLNPSAEDMGYGRHPGQPQGHALFQPLECEPTLQIGYHPDPVSVVTEGPSMNNYMAGW

                  240
KAF2292923.1        LPS
KAF2292906.1        ...
AT1G24260.2         LPY
Casp18261.t1        LPX
Ep_chr6_g20383.t1   LPX
Manes.13G009600.2.p LPX
KAF2286473.1        LPX
Manes.12G009200.2.p LPX
XP_015572264.1      LPX
Potri.003G169600.1.p LPX
Potri.001G058400.1.p LAX
Glyma.08G105500.1.p LPX
Glyma.05G148800.1.p LPX
Glyma.11G252300.1.p LPX
Glyma.18G004700.4.p LPX
```

### UFO

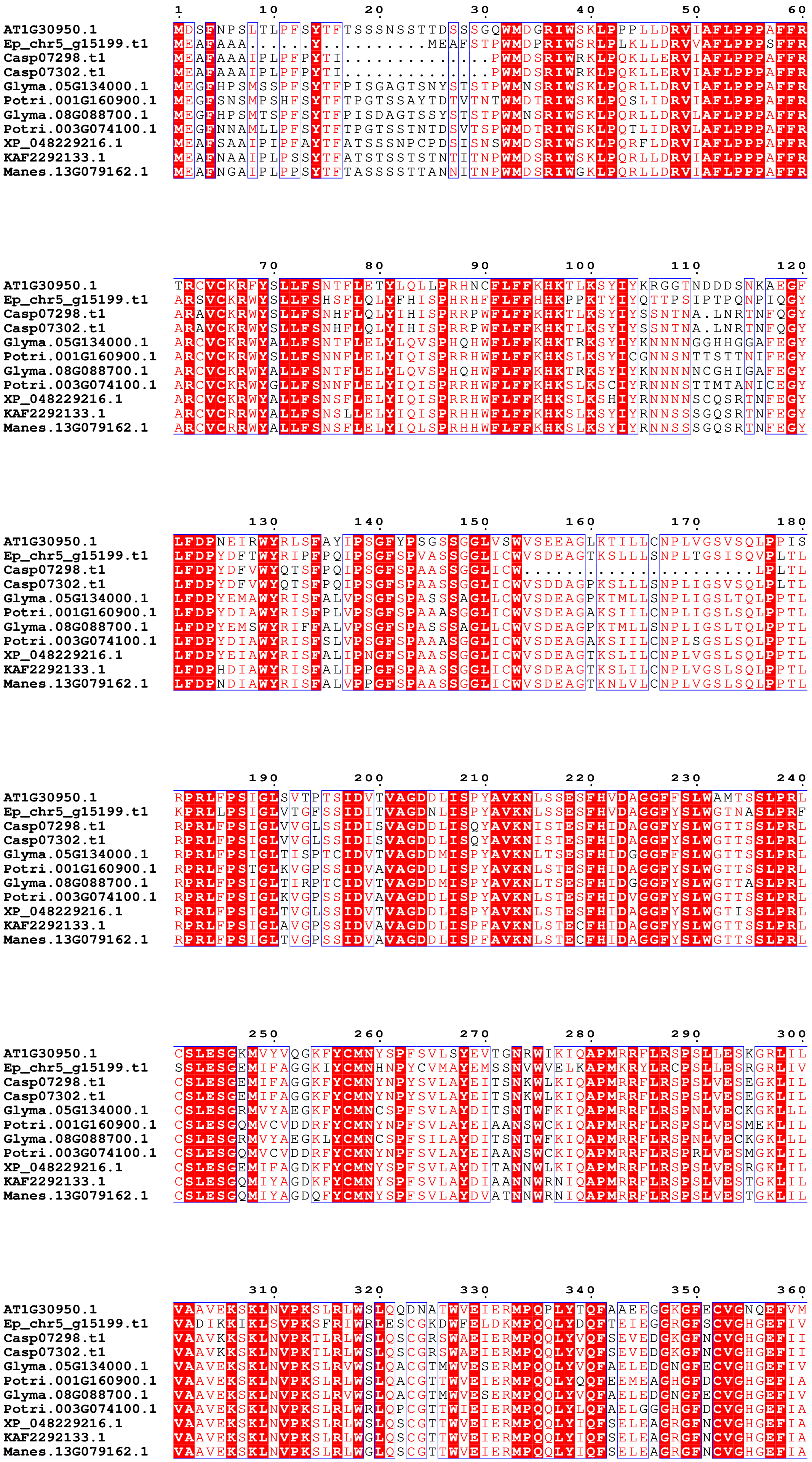
AT1G30950.1
Ep_chr5_g15199.t1
Casp07298.t1
Casp07302.t1
Glyma.05G134000.1
Potri.001G160900.1
Glyma.08G088700.1
Potri.003G074100.1
XP_048229216.1
KAF2292133.1
Manes.13G079162.1

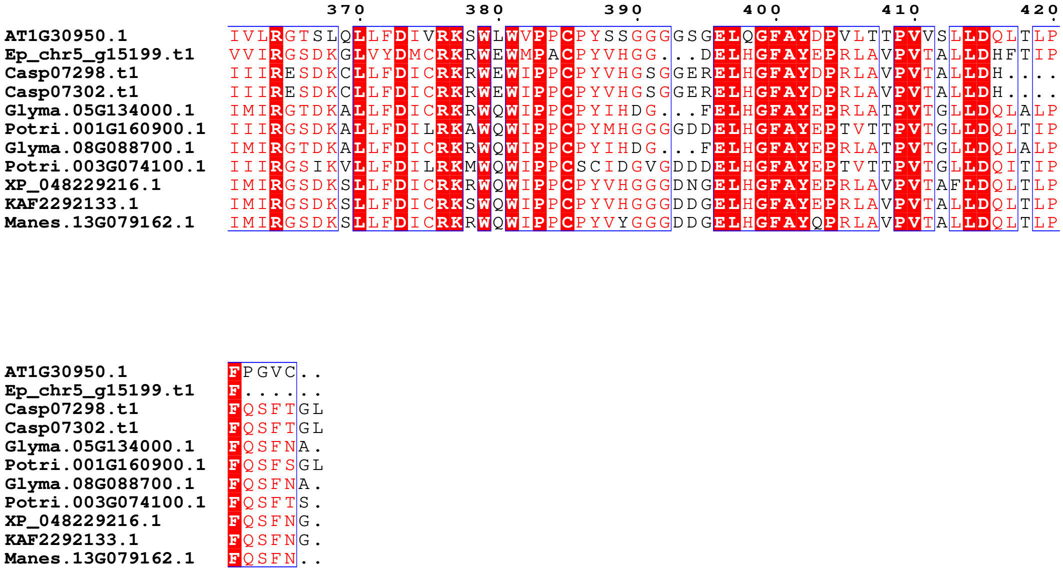
370 380 390 400 410 420
AT1G30950.1 IVLRGTSLQLLFDIVRKSWLWVPPCPYSSGGGGSGELQGFAYDPVLTTPVVSLLDQLTLP
Ep_chr5_g15199.t1 VVIRGSDKGLVYDMCRKRWEWMPACPYVHGG...DELHGFAYEPRLAVPVTALLDHFTIP
Casp07298.t1 IIIRESDKCLLFDICRKRWEWIPPCPYVHGSGGERELHGFAYDPRLAVPVTALLDH....
Casp07302.t1 IIIRESDKCLLFDICRKRWEWIPPCPYVHGSGGERELHGFAYDPRLAVPVTALLDH....
Glyma.05G134000.1 IMIRGTDKALLFDICRKRWQWIPPCPYIHDG...FELHGFAYEPRLATPVTGLLDQLALP
Potri.001G160900.1 IIIRGSDKALLFDILRKAWQWIPPCPYMHGGGGDDELHGFAYEPTVTTPVTGLLDQLTIP
Glyma.08G088700.1 IMIRGTDKALLFDICRKRWQWIPPCPYIHDG...FELHGFAYEPRLATPVTGLLDQLALP
Potri.003G074100.1 IIIRGSIKVLLFDILRKMWQWIPPCSCIDGVGDDDELHGFAYEPTVTTPVTGLLDQITIP
XP_048229216.1 IMIRGSDKSLLFDICRKRWQWIPPCPYVHGGGDNGELHGFAYEPRLAVPVTAFLDQLTLP
KAF2292133.1 IMIRGSDKSLLFDICRKSWQWIPPCPYVHGGGDDGELHGFAYEPRLAVPVTALLDQLTLP
Manes.13G079162.1 IMIRGSDKSLLFDICRKRWQWIPPCPYVYGGGDDGELHGFAYQPRLAVPVTALLDQLTLP
AT1G30950.1 FPGVC..
Ep_chr5_g15199.t1 F......
Casp07298.t1 FQSFTGL
Casp07302.t1 FQSFTGL
Glyma.05G134000.1 FQSFNA.
Potri.001G160900.1 FQSFSGL
Glyma.08G088700.1 FQSFNA.
Potri.003G074100.1 FQSFTS.
XP_048229216.1 FQSFNG.
KAF2292133.1 FQSFNG.
Manes.13G079162.1 FQSFN..
